# A genetically linked pair of NLR immune receptors show contrasting patterns of evolution

**DOI:** 10.1101/2021.09.01.458560

**Authors:** Motoki Shimizu, Akiko Hirabuchi, Yu Sugihara, Akira Abe, Takumi Takeda, Michie Kobayashi, Yukie Hiraka, Eiko Kanzaki, Kaori Oikawa, Hiromasa Saitoh, Thorsten Langner, Mark J. Banfield, Sophien Kamoun, Ryohei Terauchi

## Abstract

Throughout their evolution, plant nucleotide-binding leucine-rich-repeat receptors (NLRs) have acquired widely divergent unconventional integrated domains that enhance their ability to detect pathogen effectors. However, the functional dynamics that drive the evolution of NLRs with integrated domains (NLR-IDs) remain poorly understood. Here, we reconstructed the evolutionary history of an NLR locus prone to unconventional domain integration and experimentally tested hypotheses about the evolution of NLR-IDs. We show that the rice (*Oryza sativa*) NLR Pias recognizes the effector AVR-Pias of the blast fungal pathogen *Magnaporthe oryzae*. Pias consists of a functionally specialized NLR pair, the helper Pias-1 and the sensor Pias-2, that is allelic to the previously characterized Pia pair of NLRs: the helper RGA4 and the sensor RGA5. Remarkably, Pias-2 carries a C-terminal DUF761 domain at a similar position to the heavy metal–associated (HMA) domain of RGA5. Phylogenomic analysis showed that Pias-2/RGA5 sensor NLRs have undergone recurrent genomic recombination within the genus *Oryza*, resulting in up to six sequence-divergent domain integrations. Allelic NLRs with divergent functions have been maintained trans-species in different *Oryza* lineages to detect sequence-divergent pathogen effectors. By contrast, Pias-1 has retained its NLR helper activity throughout evolution and is capable of functioning together with the divergent sensor-NLR RGA5 to responds to AVR-Pia. These results suggest that opposite selective forces have driven the evolution of paired NLRs: highly dynamic domain integration events maintained by balancing selection for sensor NLRs, in sharp contrast to purifying selection and functional conservation of immune signaling for helper NLRs.

**Significance statement:** Plants have evolved sophisticated defense mechanisms to fend off pathogens. Plant nucleotide-binding leucine-rich repeat receptor (NLR) proteins play crucial roles in detecting pathogen molecules inside plant cells and mounting defense responses. Here, we identified the *Pias* gene from rice, which encodes the NLR pair Pias-1 “helper” and Pias-2 “sensor.” These proteins function together to detect the pathogen molecule AVR-Pias of *Magnaporthe oryzae* and defend against rice blast disease. *Pias* is allelic to the previously reported *Pia* gene. A comparison of Pias/Pia alleles among *Oryza* species showed that Pias/Pia helper is evolutionarily and functionally conserved, whereas the Pias/Pia sensor shows highly dynamic evolution, with various host domains integrated into similar positions, allowing it to detect a wide variety of pathogen molecules.

## Introduction

Plants are continually attacked by a multitude of microbial pathogens. Pathogens secrete effector molecules to enable the invasion of their hosts (Hogenhout et al. 2009). To counter this, plants have evolved a surveillance system that detects pathogen effectors inside the plant cell, leading to effector-triggered immunity (ETI; Jones and Dangl, 2006). Nucleotide-binding leucine-rich-repeat receptors (NLRs) play pivotal roles in ETI, which frequently leads to hypersensitive response (HR)–mediated cell death (Kourelis and van der Hoorn, 2018; Adachi et al. 2019a). NLR genes underwent lineage-specific expansions in most plant genomes (∼150 in *Arabidopsis thaliana* and ∼500 in rice (*Oryza sativa*)) and are among the most variable genes in plants, pointing to strong selection pressure from pathogens (Clark et al. 2007; Van de Weyer et al. 2019). NLR proteins are characterized by a conserved nucleotide-binding (NB) domain and a leucine-rich-repeat (LRR) domain. NLRs are divided into two major groups depending on the type of N-terminal domain: NLRs with the N-terminal coiled-coil (CC) domain are called CNLs (CC-NLRs), and those with the N-terminal Toll-like domain are called TNLs (TIR-NLRs). CNLs are widespread in the plant kingdom. TIRs are grouped into the canonical TIR and TIR2 subclasses. Though TIR2-NB proteins are found in monocot plants, TNLs with canonical TIR domains have been detected in dicot but not in monocot plants (Pan et al. 2000; Sarris et al. 2016). ADP/ATP exchange at the NB domain (Bernoux et al. 2016; Wang et al. 2019a) and oligomerization of NLRs (Ma et al. 2020; Martin et al. 2020; Sharif et al. 2019; Tenthorey et al. 2017; Wang et al. 2019b; Zhang et al. 2015) trigger ETI signaling. The Arabidopsis CNL ZAR1 forms a pentamer “resistosome” complex after binding to a host protein and a pathogen effector. This leads to the protrusion of the N-terminal alpha helix, which perturbs the plasma membrane to trigger the HR (Wang et al. 2019ab), a process potentially mediated by Ca^2+^ influx (Bi et al. 2021). The Arabidopsis TNL RPP1 forms a tetramer “resistosome” upon binding to its cognate effector ATR1, followed by an increase in its NAD^+^ase activity (Ma et al. 2020), which triggers cell death (Wan et al. 2019; Horsefield et al. 2019). These recent breakthroughs have started to reveal the biochemical links between NLR molecular structure and HR induction (Adachi et al. 2019b).

NLRs can function as singletons, in pairs or in networks (Adachi et al. 2019a). Arabidopsis ZAR1 (Wang et al. 2019a), RPP1 (Krasileva et al. 2010), and many other NLRs function as singleton NLRs and recognize AVRs directly or indirectly. Two NLR proteins encoded by genetically linked genes function together as paired NLRs. One of the paired NLRs frequently has a non-canonical domain called the integrated domain (ID). IDs are thought to have been derived from other host proteins (Kroj et al. 2016; Sarris et al. 2016). Examples of paired NLRs include RPS4/RRS1 in Arabidopsis (Williams et al. 2014; Le Roux et al. 2015) and RGA4/RGA5 (Okuyama et al. 2011; Cesari et al. 2013), Pik1/Pik2 (Ashikawa et al. 2008; Maqbool et al. 2015), and Pii1/Pii2 in rice (Takagi et al. 2013; Fujisaki et al. 2017). Multiple NLRs encoded by unlinked genes may function together; these network NLRs (Wu et al. 2017) include NRC2, NRC3, and NRC4 in Solanaceae plants. When NLRs function as a pair or network, one NLR is involved in the recognition of the AVR effector (the “sensor NLR”), whereas the other plays a role in signaling (the “helper NLR”). Recent studies have expanded our understanding of the genetic architecture of NLR pairs and networks; however, how they function together and how they evolved remain elusive.

A phylogenetic study of 4,184 NLRs in the genomes of seven Poaceae species including rice, wheat, and other grass species grouped the NLRs into 24 major clades, including three clades (MIC1, MIC2, and MIC3) containing the majority of NLRs with IDs (Bailey et al. 2018). NLRs in clade MIC1 (including RGA5) are characterized by a wide variety of IDs integrated into similar positions in the NLR after the LRR domain. Bailey et al. (2018) reported a conserved 43–amino acid motif between LRR and ID named the CID (conservation and association with IDs) motif. The authors proposed a possible evolutionary mechanism of ID generation and ID shuffling mediated by the CID motif. However, this study examined only a few genomes from each species, with a focus on inter-species diversity within the Poaceae, and provided limited information about the recent diversification of NLRs within a single genus. In addition, experimental validation of the hypotheses underpinning the functional diversity of NLRs with IDs remains limited.

Rice blast disease caused by blast fungus (*Magnaporthe oryzae* [syn. *Pyricularia oryzae*]) is a major disease of rice that threatens world food security (Pennisi 2010). The best way to control this disease is to deploy cultivars with resistance (*R*) genes. To date, 40 *R*-genes against blast have been reported, and 25 have been cloned, most of which encode NLRs (Liu et al. 2010; Wu et al. 2015). However, the molecular interactions between rice NLRs and blast AVRs are understood for only a small number of cases, including Pita and AVR-Pita (Orbach et al. 2000; Jia et al. 2000), Pia and AVR-Pia/AVR1-CO39 (Cesari et al. 2013), Pik and AVR-Pik (Kanzaki et al. 2012; Maqbool et al. 2015), Pii and AVR-Pii (Takagi et al. 2013; Fujisaki et al. 2015; 2017), and Pizt and AVR-Pizt (Park et al. 2016, Wang et al. 2016).

The rice Pia pair, consisting of the NLR helper RGA4 and the NLR sensor RGA5, is one of the most well-characterized NLR pairs (Okuyama et al. 2011; Cesari et al. 2013). Overexpression of *RGA4* in *Nicotiana benthamiana* and rice protoplasts caused HR-like cell death in the respective plant species, suggesting that the helper RGA4 is responsible for HR signaling. RGA4-mediated cell death was suppressed by co-expression of RGA5, indicating that RGA5 negatively regulates RGA4-mediated defense signaling. Finally, co-expression of RGA5, RGA4, and AVR-Pia triggered cell death via the direct binding of AVR-Pia to HMA ID of the sensor RGA5 (Cesari et al. 2013). This study, together with a study of the Arabidopsis RPS4/RRS1 pair (Williams et al. 2014; Le Roux et al. 2015), supported a negative regulation model for paired NLRs in which helper NLRs function in HR signaling and sensor NLRs in the suppression of helper NLR activity. Upon binding or modification of the sensor-NLR IDs by pathogen AVRs, this suppression is released, allowing HR signaling to proceed. However, a recent study of rice Pikp paired NLRs suggests a helper-sensor cooperation model. Expression of the helper NLR alone did not cause cell death in *N. benthamiana* and co-expression of the helper and sensor was required for triggering effector-dependent HR-like cell death (Zdrzałek et al. 2020).

Here, we describe the rice NLR pair Pias, which recognize the *M. oryzae* effector AVR-Pias. *Pias* encodes the helper Pias-1 and the sensor Pias-2 and is allelic to the previously characterized *Pia* gene, encoding the NLR pair helper RGA4 and sensor RGA5. Pias-2 carries a C-terminal DUF761 domain at a similar position to the heavy metal–associated (HMA) domain of RGA5. We show that the Pias/Pia helper NLR lineage is evolutionarily and functionally conserved, while its sensor NLR lineage shows highly dynamic evolution with various host domains integrated into similar positions, possibly allowing it to detect a wide variety of pathogen molecules.

## Results

### Isolation of rice *Pias* NLR genes and the matching gene *AVR-Pias* from *Magnaporthe oryzae*

As part of a large genetic screen to identify novel rice blast resistance genes, we crossed *japonica-*type rice cultivar Hitomebore to 20 different rice cultivars representing the worldwide genetic diversity of rice, resulting in the generation of recombinant inbred lines (RILs) of the F_7_-F_9_ generations. We inoculated the parental rice lines with a panel of *M. oryzae* isolates and recorded their resistance or susceptibility to each isolate. Since Hitomebore was susceptible to *M. oryzae* isolate 2012-1 but *indica*-type accession WRC17 (cultivar Keiboba; Kojima et al. (2005)) was resistant to this isolate, we set out to isolate the resistance genes in WRC17 rice against the 2012-1 pathogen. The 58 RILs derived from a cross between Hitomebore and WRC17 segregated into 52 resistant and 6 susceptible lines (Fig. 1A), indicating that WRC17 likely contains more than one locus conferring resistance against 2012-1.

**Figure 1.**
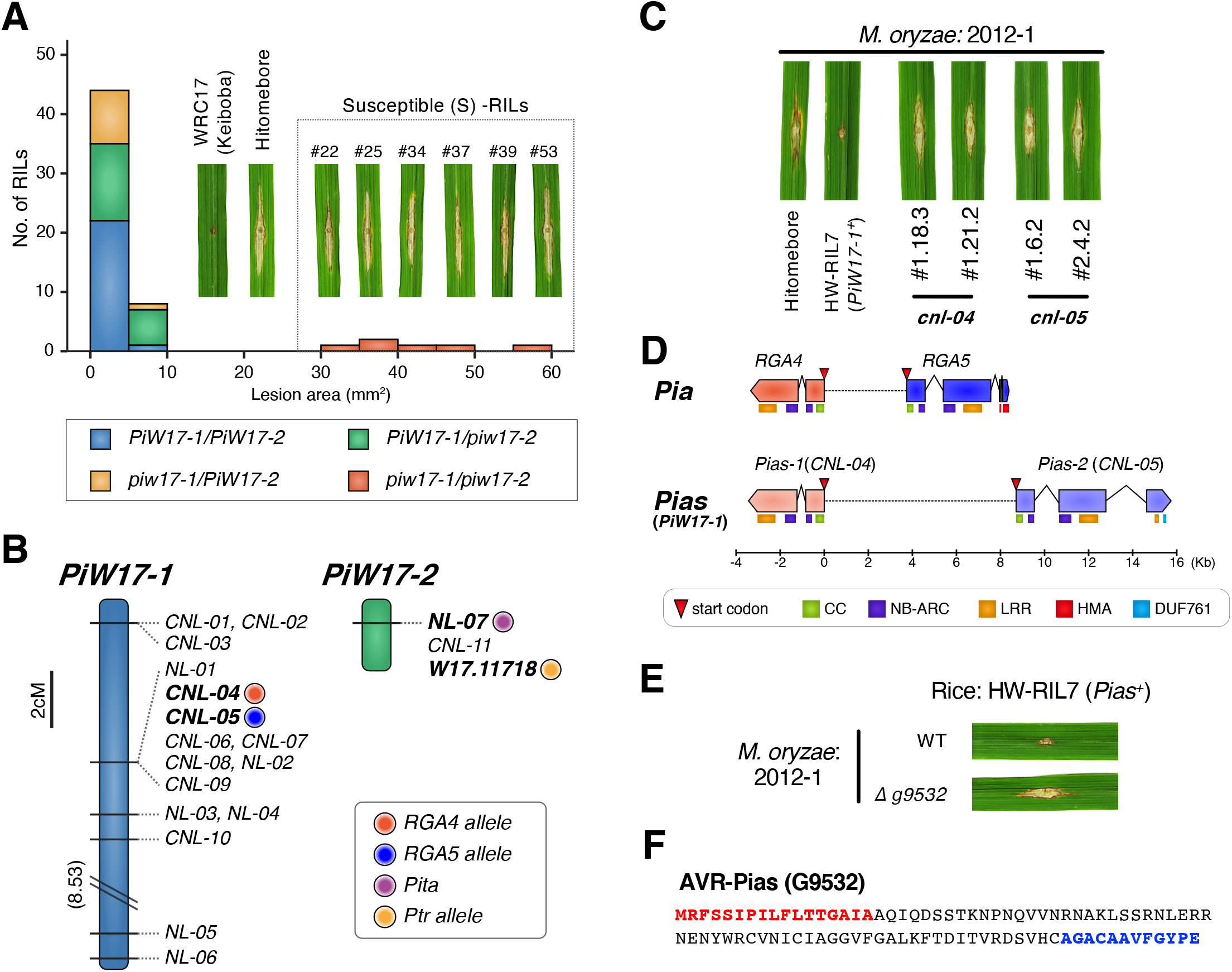
The *Pias* gene of rice line WRC17 (cultivar Keiboba) encodes a CC-NLR protein and is allelic to *Pia*. (A) Segregation of the resistance and susceptibility traits among the 58 RILs derived from a cross between WRC17 (cultivar Keiboba) and Hitomebore. Disease symptoms of WRC17, Hitomebore, and six RILs showing a susceptible phenotype after punch inoculation of *Magnaporthe oryzae* isolate 2012-1 (leaf photographs) and the frequency distribution of disease lesion area of the 58 RILs (bar graphs). (B) Linkage maps of candidate NLR genes at the *PiW17-1* and *PiW17-2* loci. (C) Both *CNL-04* and *CNL-05* are required for *PiW17-1*-mediated resistance against *M. oryzae* 2012-1. HW-RIL7 contains only *PiW17-1* and is resistant to *M. oryzae* 2012-1. Knockout of *CNL-04* (*cnl-04*) and *CNL-05* (*cnl-05*) in HW-RIL7 rendered plants susceptible to 2012-1. (D) Gene structures of *Pia* consisting of *RGA4* and *RGA5* and *Pias* consisting of *Pias-1* and *Pias-2*. The positions of protein domains (CC, NB-ARC, LRR, HMA, and DUF761) encoded by the NLRs are indicated. (E) The *M. oryzae* 2012-1 *AVR-Pias* knockout mutant became virulent to HW-RIL7. (F) Amino acid sequence of G9532 protein (AVR-Pias). The secretion signal is indicated by red letters and the Toxin18-like motif by blue letters. The Toxin18-like motif was annotated by Pfam (https://pfam.xfam.org/).

To identify the resistance genes, we performed whole-genome sequencing and RNA-seq of WRC17 and conducted an association study using a bioinformatics pipeline we named “RaIDeN” (https://github.com/YuSugihara/RaIDeN) using the RILs segregating for these phenotypes (Fig. 1; Suppl. Fig. 1). We sequenced the genomes WRC17, Hitomebore, and six RILs showing susceptibility to the isolate 2012-1 on an Illumina DNA sequencer (**Suppl. Table 1**) and subjected the short reads of WRC17 to *de novo* assembly with *DISCOVAR* (https://www.broadinstitute.org/software/discovar/blog/), resulting in the WRC17 reference genome sequence (**Suppl. Table 2**). We also performed RNA-seq of WRC17 leaves that had been inoculated with *M. oryzae* (2012-1). The RNA-seq reads were mapped to the WRC17 reference genome, revealing 22,561 genes expressed from the WRC17 genome. The short reads obtained from Hitomebore and six susceptible (S-) RILs were aligned to the genome sequences of the 22,561 expressed genes. We reasoned that Hitomebore and the six S-RILs share the same DNA sequences in the candidate genes responsible for their resistance that are different from the sequences in resistant WRC17 (Suppl. Fig. 1). Two types of DNA polymorphisms were considered: (1) presence/absence of the genes and (2) SNPs in the genes. We identified 14 genes that were present in WRC17 but absent from Hitomebore and the six S-RILs, in addition to 839 genes with shared SNPs among Hitomebore and the S-RILs that were different from the sequences of WRC17.

From this group of 853 genes (**Suppl. Dataset 1**), we selected Resistance Gene Analogs (RGAs) using “RGAugury” (Li et al. 2016), which predicts genes encoding putative NLRs, Receptor-Like Kinases (RLKs), and Receptor Like Proteins (RLPs). This analysis identified 38 RGAs as the candidate genes (**Suppl. Table 3**). Since most rice resistance genes against *M. oryzae* reported to date are *NLR*s, we focused on 18 *NLR* (11 *CNL* and 7 *NL*) genes as the candidate resistance genes of WRC17 against *M. oryzae* isolate 2012-1. Among the 18 NLR genes, only one (*NL-04*) showed presence/absence polymorphisms, and the rest contained SNPs associated with the phenotypes. We developed DNA markers in the candidate NLR genes and studied their association with phenotypes using 58 RILs segregating for resistance and susceptibility to the isolate 2012-1 (Suppl. Fig. 2A). This analysis showed that 16 NLRs were tightly linked to each other and that the two other *NLR*s were linked to each other, suggesting that the two loci (designated *Pi-W17-1* and *Pi-W17-2*) are involved in the resistance of WRC17 against 2012-1 (Fig. 1B). Genotyping of the 52 RILs using the markers located in the two loci, *Pi-W17-1* (*CNL-04*) and *Pi-W17-2* (*NL-07* and *W17.11718*), suggested that the lines became resistant when either of the two loci contained the WRC17-type allele (Fig. 1A; Suppl. Fig. 2A).

The genomic position of *Pi-W17-1* corresponds to that of the previously reported *Pia* locus (Okuyama et al. 2011), and the position of *Pi-W17-2* corresponds to that of the *Pita* (=*NL-07*) and *Ptr* (=*W17.11718*) loci (Zhao et al. 2018; Orbach et al. 2000; Meng et al. 2020). Among the progeny showing resistance against 2012-1, we selected 19 RILs containing *PiW17-1* but not *PiW17-2*. These lines shared 10 candidate NLR genes within the *PiW17-1* region. (Fig. 1B; Suppl. Fig. 2B). We performed RNAi-mediated gene silencing of eight of these genes (encoding proteins over 900 amino acids long) (Suppl. Fig. 2C) using the RIL HW-RIL7, which contains *Pi-W17-1* but lacks *Pi-W17-2*. When *CNL-04* or *CNL-05* was silenced, its resistance against 2012-1 became compromised (Suppl. Fig. 2CD). We confirmed this result by generating *CNL-04* and *CNL-05* knockout mutant lines by CRISPR/Cas9-mediated genome editing (Fig. 1C; Suppl. Fig. 3). These data suggest that *Pi-W17-1*-mediated resistance requires both the neighboring NLRs *CNL-04* and *CNL-05*. Indeed, the position of *CNL-04* corresponds to that of *RGA4*, whereas the position of *CNL-05* corresponds to that of *RGA5* of *Pia* (Fig. 1D; Okuyama et al. 2011). Interestingly, *CNL-04* is similar to *RGA4* in terms of both structure and DNA sequence (96.6% DNA sequence identity), whereas *CNL-05* has a distinct structure and a DNA sequence that diverged from *RGA5* (59.8% DNA sequence identity) (Suppl. Fig. 4). *Pia RGA5* encodes a protein with a heavy metal–associated (HMA) domain in its C terminus (Okuyama *et al.* 2011), whereas *CNL-05* encodes a protein with a 19–amino acid motif corresponding to domain of unknown function 761 (DUF761) near its C terminus. The physical distance between *CNL-04* and *CNL-05* is 8.7 kb, which is longer than that between *RGA4* and *RGA5* (3.7 kb) (Fig. 1D). In view of the substantial differences between *CNL-05* and *RGA5*, we decided to name this WRC17 allele *Pias*, *CNL-04* as *Pias-1*, and *CNL-05* as *Pias-2* (Suppl. Fig. 5).

To isolate the *AVR-Pias* avirulence gene cognate of rice NLR *Pias*, we performed an association study of expressed genes encoding candidate effector proteins (See Suppl. Fig. 6 and **Suppl. Table 4, 5** for details). This analysis identified three genes (*G9141*, *G9435*, and *G9532*) as candidates of *AVR-Pias*. We selected *M. oryzae* isolate Ao92-06-2, which is compatible with HW-RIL7, transformed it with each of the candidate genes, and tested their interactions with HW-RIL7 (Suppl. Fig. 7). Transformation with one of the candidate genes, *G9532*, rendered Ao92-06-2 incompatible with HW-RIL7 (Suppl. Fig. 7A), suggesting that *G9532* is *AVR-Pias*. To validate this result, we generated a knockout mutant of *G9532* in the 2012-1 background, which became compatible with HW-RIL7 (Fig. 1E; Suppl. Fig. 8). These results indicate that *G9532* is *AVR-Pias*, which is recognized by *Pias*. Both *Pias-1* and *Pias-2* are required for the recognition of *AVR-Pias*, as knockout of either NLR gene abrogated resistance (Suppl. Fig. 7B). AVR-Pias is a 91–amino acid protein with a secretion signal peptide (Fig. 1F). This protein contains a 12–amino acid Toxin18-like motif (a feature of proteins belonging to the conotoxin O superfamily) in its C terminus.

### Various host domains are integrated into Pias/Pia sensor NLRs

*Pias* and *Pia* are allelic to each other, and both are composed of a pair of NLRs. The helper NLRs (*Pias-1* and *RGA4*) are conserved, whereas the sensor NLRs (*Pias-2* and *RGA5*) are divergent, with different integrated domains. To explore the diversity and evolution of *Pias/Pia* NLRs in the entire *Oryza* genus, we obtained the genomic DNA sequences of the *Pias/Pia* locus from 171 accessions representing 11 *Oryza* species as well as four non-*Oryza* species of Poaceae, including *Setaria italica* (foxtail millet), *Panicum hallii* (Hall’s panicgrass), *Hordeum vulgare* (barley), and *Aegilops tauschii* (Tausch’s goatgrass) (**Suppl. Table 6**). To validate the gene structures, we used *RGA4/RGA5* and *Pias-1/Pias-2* genes as well as gene models supported by RNA-seq for 10 *Oryza* accessions as queries to infer the gene models of 167 *Oryza* accessions and four non-*Oryza* species using Exonerate software (http://www.ebi.ac.uk/~guy/exonerate) (**Suppl. Table 7, 8**). The details of the gene model prediction pipeline and the results of RNA-seq alignment to the Pias-2/RGA5 genes are given in Suppl. Fig. 9. Remarkably, Pias-2/RGA5 sensor proteins contain a wide variety of IDs, with up to nine different domains, including HMA, DUF761, DUF677, Zinc_ribbon_12, PKc_MAPKK (PKc_M), PKc_like, and WRKY, inserted into the identical position after the LRR domain (Fig. 2A). Around the junction of the ID and LRR domains, we identified a conserved 145– to 146–amino acid motif named the LRR-ID intervening (LII) region (Fig. 2B), which spans four LRR domains. The LII region partially overlaps but is different from the previously reported CID motif (Bailey et al. 2018; Suppl. Fig. 10).

**Figure 2.**
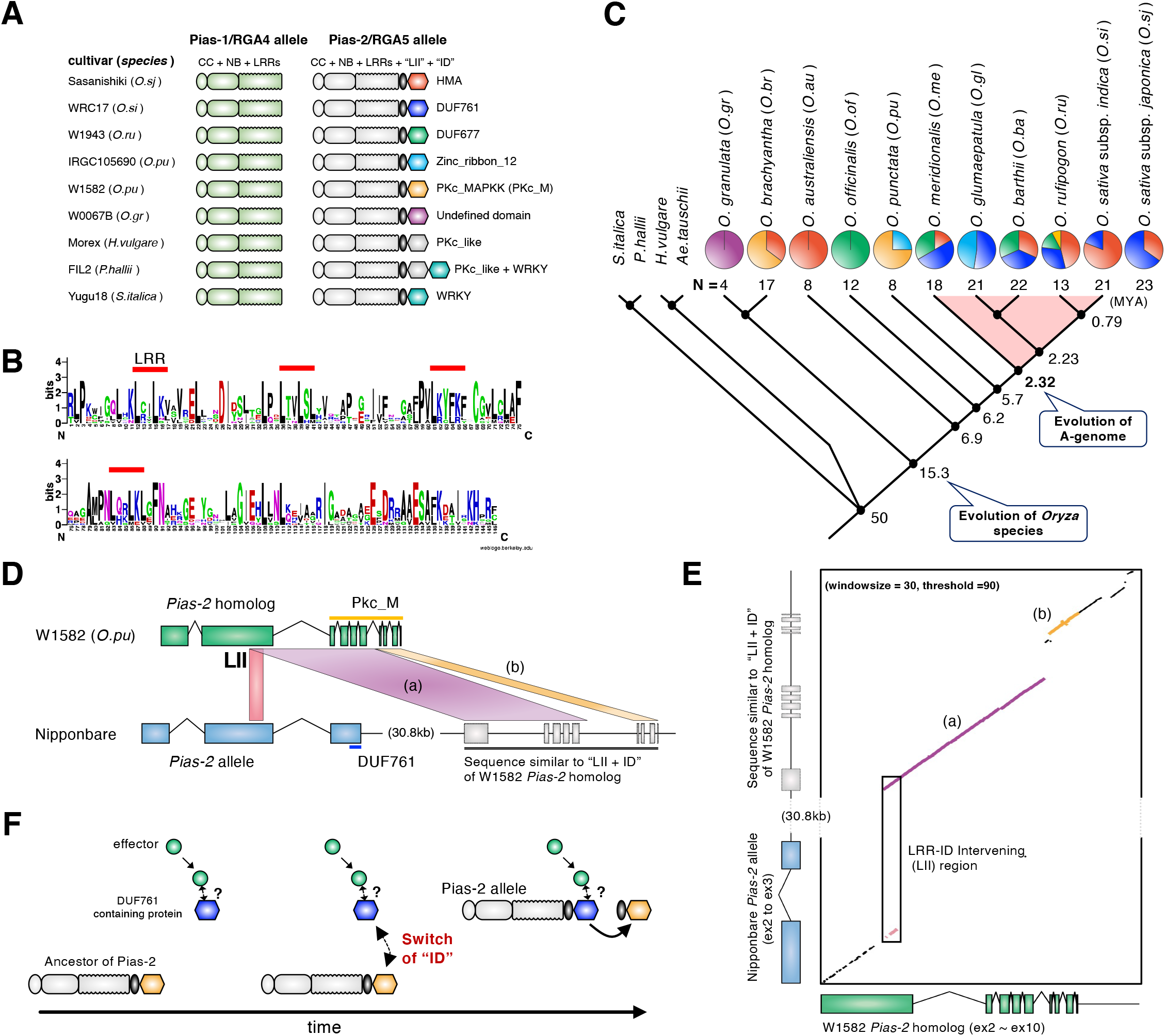
Recurrent integration of extraneous domains in Pias/Pia sensor NLRs. (A) A simplified scheme of the structures of the Pias/Pia NLR pairs. Pias-1/RGA4 helper NLRs are shown in green, and Pias-2/RGA5 sensor NLRs are shown in white. The conserved LRR-ID intervening motif (LII) is indicated by black ellipses. Integrated domains are shown by different-colored hexagons. (B) A sequence logo showing conserved amino acids of the LRR-ID Intervening (LII) motif. The red lines indicate the LRR in LII motif. (C) Distribution of ID motifs among *Oryza* species. The pie charts show the frequencies of different ID motifs in a given species. The colors correspond to the ID colors in (A). The numbers below the pie charts indicate the sample numbers. A cladogram showing the phylogenetic relationships of 11 *Oryza* species and four other Poaceae species (*Setaria italica*, *Panicum hallii*, *Hordeum vulgare*, and *Aegilops tauschii*) based on Time Tree, the time scale of life web database (http://www.timetree.org/). The numbers on the branches indicate the estimated time of the splitting of lineages (MYA: million years ago). (D) DNA sequence similarity between *O. punctata Pias-2/RGA5 sensor NLR* and the downstream sequence of *O. sativa* (Nipponbare) *Pias-2/RGA5.* LII: LRR-ID intervening motif. (E) Dot-plot analysis of the *O. punctata* (W1582) *Pias-2/RGA5 sensor NLR* and *O. sativa* (Nipponbare) *Pias-2/RGA5 NLR* downstream sequences using the Dotmatcher tool (http://emboss.bioinformatics.nl/cgi-bin/emboss/dotmatcher/). (F) Possible evolutionary process of ID replacement that might have occurred between the *O. punctata* and the *O. sativa Pias-2*/*RGA5* lineages. We still do not know the mode of interaction between the AVR-Pias effector and DUF761-containing protein, so it is indicated by “?”.

We studied the frequency of different IDs in the 11 *Oryza* species used to reconstruct a phylogenetic tree based on whole-genome sequences (Fig. 2C; **Suppl. Table 6, 8**). The IDs of the cultivated rice *O. sativa* (44 samples) are shared by the HMA and DUF761 domains with similar frequencies. *O. rufipogon* (13 samples), the wild progenitor of *O. sativa*, contains DUF677 and PKc_M in addition to HMA and DUF761. *Oryza* species belonging to the A-genome group (*O. sativa*, *O. rufipogon*, *O. barthii*, *O. glumaepatula*, and *O. meridionalis*) contain a higher proportion of the DUF761 ID, suggesting its importance in their defense. However, outside the A-genome species, the DUF761 ID is absent. Instead, PKc_M (*O. brachyantha* and *O. punctata*), DUF677 (*O. officinalis*), and HMA (*O. australiensis*) are dominant, indicating that the integration of the DUF761 domain into Pias-2/RGA5 likely occurred in the immediate ancestor of the A-genome species that diverged from other *Oryza* lineages over 2.3 million years ago (TimeTree; http://www.timetree.org/).

A comparison of the genome sequences around *Pias/Pia* sensor-NLR genes between *O. punctata* and *O. sativa* cultivar Nipponbare revealed an interesting conserved region (Fig. 2DE). Both of their coding regions contain conserved LII sequences, whereas the IDs downstream of the LII are different: PKc_M for *O. punctata* W1582 and DUF761 for *O. sativa* Nipponbare. However, in the 30.8-kb region downstream of the *O. sativa* Nipponbare *Pias-2* gene, two regions (block **a** and **b**) share high DNA sequence similarity with the LII and ID sequences of *O. punctata* W1582 Pias-2 homolog (Fig. 2DE; Suppl. Fig. 11): Block **a** corresponds to region LII to exon-6, and block **b** corresponds to exon-7 to −10 of the *O. punctata* W1582 *Pias-2* homolog. Perhaps this conserved sequence downstream of the *O. sativa* Nipponbare *Pias* sensor is a footprint of the replacement of the ID from PKc_M to DUF761 via homologous recombination at the LII region (Fig. 2F). A survey of 51 A-genome *Oryza* accessions with the DUF761 ID revealed that the *O. punctata* W1582 LII-ID-like sequence is widely conserved in *O. sativa*, *O. rufipogon*, *O. meridionalis*, and an accession of *O. barthii* (Accession W1702), but not in the majority of *O. barthii* and *O. glumaepatula* accessions (**Suppl. Table 9**; Suppl. Fig. 12). These results indicate that this possible recombination occurred only once and that the footprint was probably lost in the latter two species. We did not detect similar footprints in other sensor *Pias/Pia* NLRs with non-DUF761 IDs.

### Contrasting patterns of evolution of *Pias/Pia* sensors and helpers

To explore the evolutionary patterns of the *Pias/Pia* NLR locus, we reconstructed phylogenetic trees of *Pias*/*Pia* NLR pairs separately for the helper NLR (Pias-1 and RGA4) and the sensor NLR (Pias-2 and RGA5) using the amino acid sequences of the helper NLRs (full-length protein) and sensor NLRs (full-length protein except the ID domain) of 22 *Oryza* accessions and four accessions from other Poaceae genera. Note that Pias/Pia helper and sensor NLRs belong to distantly related NLR clades (Bailey et al. 2018). Overall, the helper NLR tree (Fig. 3A) is consistent with the species tree (Fig. 2C) with a few exceptions. Species in the AA genome group (*O. sativa* subsp. *japonica*, *O. sativa* subsp. *indica*, *O. rufipogon*, *O. barthii*, *O. glumaepatula*, and *O. meridionalis*) clustered together. Only the placement of *O. australiensis* (accession W0008) is not congruent between the Pias-1/RGA4 helper NLR tree and the species tree. On the other hand, the sensor-NLR tree is quite distinct from the helper NLR tree and is not congruent to the species tree. Remarkably, the sensor NLRs show a higher level of divergence than the helper NLRs (Fig. 3A). The sensor phylogeny includes two major clades (C1and C2) separated by deep branches. RGA5 of *O. sativa* cv. Sasanishiki (Okuyama et al, 2011) belongs to the C1 clade, whereas Pias-2 of *O. sativa* WRC17 belongs to the C2 clade. Four species of the AA genome group (*O. sativa* subsp*. japonica*, *O. sativa* subsp*. indica*, *O. rufipogon*, *O. barthii*, and *O. meridionalis*) and *O. punctata* (BB genome) have sensor-NLR alleles from both the C1 and C2 clades, whereas *O. australiensis* (EE genome), *O. granulata* (GG genome), and *O. brachyantha* (FF genome) have C1 alleles, and *O. glumaepatula* (AA genome) and *O. officinalis* (CC genome) have C2 alleles. Therefore, phylogenetic analysis of the sensor NLR of the *Pias/Pia* locus pointed to trans-species polymorphism, which is reminiscent of the major histocompatibility (MHC) gene polymorphism in vertebrates (Edwards and Hedrick, 1998).

**Figure 3.**
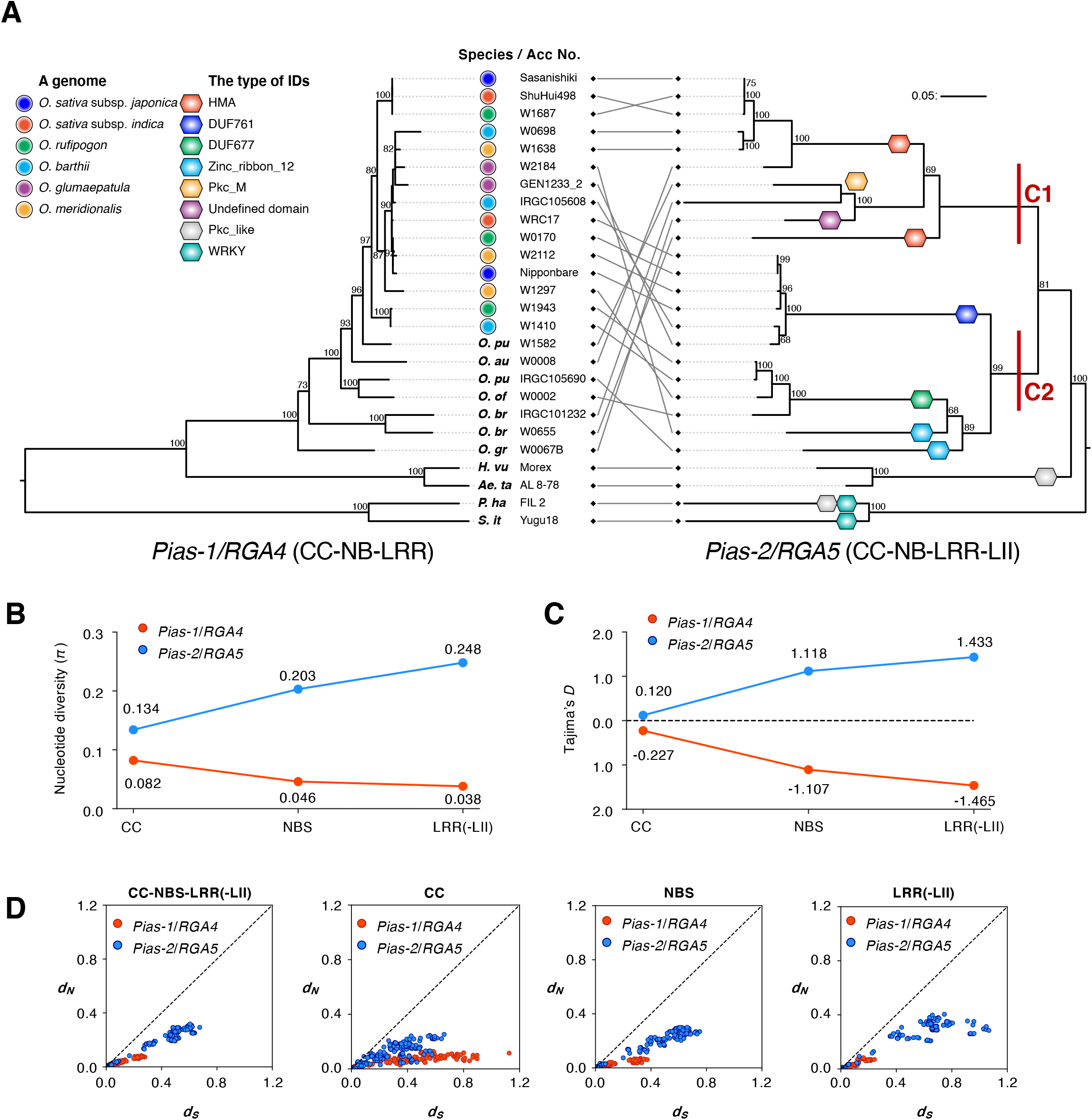
Contrasting evolutionary patterns of the helper and sensor NLRs of the *Pias/Pia* locus. (A) Phylogenetic tree of the *Pias-1/RGA4* helper NLR gene (left) and *Pias-2/RGA5* sensor NLR (right) gene based on the full-length amino acid sequence of Pias-1/RGA4 and the sequence in the region CC to LII for Pias-2/RGA5*. Pias-2/RGA5* sensor NLRs form two major clades (C1 and C2). The numbers indicate bootstrap values. (B) Nucleotide diversity (*π*) of the CC, NBS and LRR(-LII) domains of *Pias-1*/*RGA4* helper NLR gene and *Pias-2*/*RGA5* sensor NLR gene in 22 *Oryza* samples. (C) Tajima’s *D* of the CC, NBS and LRR(-LII) domains of *Pias-1*/*RGA4* helper NLR gene and *Pias-2*/*RGA5* sensor NLR gene in 22 *Oryza* samples. (D) Pairwise *d_N_* and *d_S_* values of CC-NBS-LRR(-LII), CC, NBS and LRR domains of the *Pias-1*/*RGA4* helper and *Pias-2*/*RGA5* sensor NLR gene in 22 *Oryza* samples.

To determine whether the paired NLRs have accumulated mutations at different rates, we calculated the nucleotide diversity and Tajima’s *D* (Tajima, 1989). The nucleotide diversities of the *Pias-2/RGA5* sensor was markedly higher than that of the *Pias-1/RGA4* helper for each of the CC, NBS and LRR(-LII) domains (Fig. 3B). Also, the three domains of *Pias-1/RGA4* helper and *Pias-2/RGA5* sensor had contrasting negative and positive Tajima’s *D*, respectively (Fig. 3C), indicating purifying selection in *Pias-1/RGA4* and balancing selection in *Pias-2/RGA5*, especially in the LRR domain. These results suggest that the genetically linked helper and sensor NLRs of the *Pias/Pia* locus in the genus *Oryza* have undergone contrasting modes of evolution.

To determine the types of nucleotides substitutions in the *Pias-1/RGA4* helper and *Pias-2/RGA5* sensor, we calculated *d_N_* (non-synonymous mutations) and *d_S_* (synonymous mutations) (Fig. 3D). Consistent with higher nucleotide diversity and Tajima’s *D*, we noted overall higher *d_N_* and *d_S_* in *Pias-2/RGA5*, notably in NBS and LRR domains (Fig. 3D). We also observed that the pairwise *d_N_* and *d_S_* of *Pias-1/RGA4* was constrained by the low genetic divergence in many of the examined pairs. Nonetheless, *d_S_* was higher in *Pias-1/RGA4* for CC domain than in *Pias-2/RGA5* (Fig. 3D; Suppl. Fig. 13). These results suggest that different evolutionary patterns affect the different domains of *Pias-1/RGA4* and *Pias-2/RGA5*.

### Pias-1 helper functions with the RGA5 sensor to recognize AVR-Pia

Pia comprises the helper NLR RGA4 and the sensor-NLR RGA5. The expression of RGA4 triggers HR cell death in rice as well as *N. benthamiana*, and RGA5 suppresses RGA4-mediated cell death (Césari et al. 2014). Upon the binding of AVR-Pia to HMA ID of RGA5, this suppression is released, and HR-like cell death is triggered (Césari et al. 2014). To address the functional conservation of helper/sensor NLRs of the *Pias/Pia* locus, we examined the functions of helper NLRs by transiently overexpressing five Pias/Pia helper NLRs in *N. benthamiana* using agroinfiltration (Suppl. Fig. 14). Only RGA4 supported strong cell death, and other alleles including Pias-1 caused weaker cell death, indicating that the role of RGA4 as a strong HR inducer is not typical among the tested helper NLRs (Suppl. Fig. 14). Next, we tested the effects of co-expressing Pias-1 and Pias-2, which surprisingly resulted in stronger cell death than that caused by Pias-1 expression alone (Suppl. Fig. 15). Co-expression of *AVR*-*Pias* with *Pias-1* and *Pias-2* did not alter the level of cell death (Suppl. Fig. 15). A co-immunoprecipitation analysis showed that Pias-1 and Pias-2 interact (Suppl. Fig. 16), and a yeast two-hybrid assay showed that the CC-domains of Pias-1 and Pias-2 form homo- and heterodimers (Suppl. Fig.16). These results suggest that Pias-1 and Pias-2 physically interact like RGA4/RGA5, but their mode of action is different from that of the RGA4/RGA5 pair.

We next examined the effects of exchanging various helper and sensor Pias/Pia NLRs by performing a transient expression assay in *N. benthamiana* leaves (Fig. 4A, Suppl. Fig. 17). Co-expression of the sensor RGA5 with the two heterologous helpers (Pias-1 and RGA4-Ogr from *O. granulata*) suppressed HR-like cell death, indicating that RGA5 can suppress the cell death induced by helper NLRs other than RGA4 (Fig. 4A, Suppl. Fig. 17). Furthermore, this RGA5-mediated suppression of the Pias-1-triggered HR was released via co-expression of AVR-Pia, which resulted in cell death (Fig. 4A; Suppl. Fig. 17). A similar result was obtained using RGA4-Ogr helper (Fig. 4A; Suppl. Fig. 17). These results indicate that the sensor RGA5 properly functions with helpers other than RGA4 to recognize AVR-Pia and mount the HR in *N. benthamiana*.

**Figure 4.**
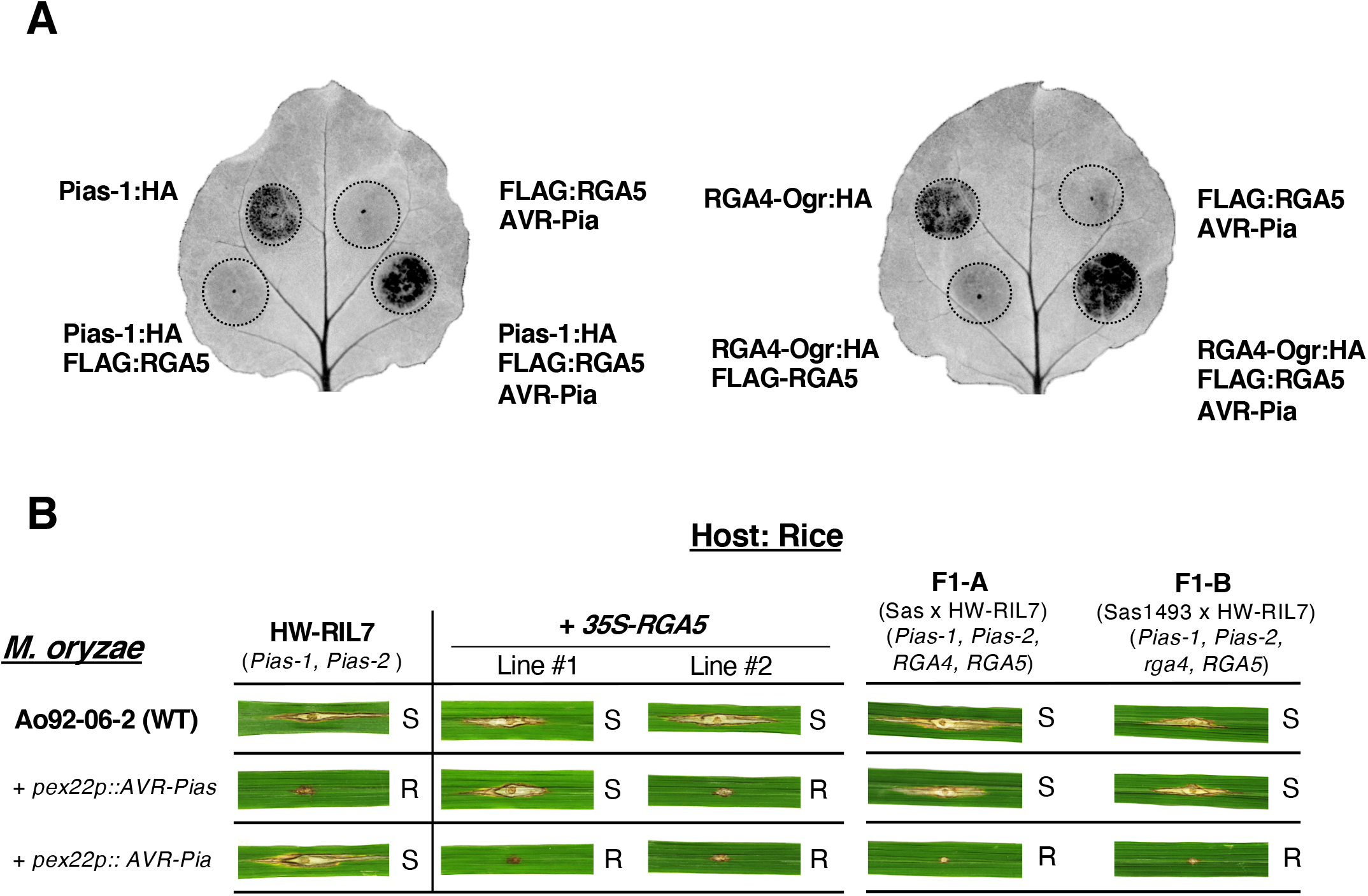
The NLR helper Pias-1 is functionally conserved. (A) Representative images of *N. benthamiana* leaves after agroinfiltration with Pias-1:HA, Pias-1:HA/FLAG:RGA5, FLAG:RGA5/AVR-Pia, and Pias-1:HA/FLAG:RGA5/AVR-Pia (left) and RGA4-Ogr:HA derived from *O. granulata*, RGA4-Ogr:HA/FLAG:RGA5, FLAG:RGA5/AVR-Pia, and RGA4-Ogr:HA/FLAG:RGA5/AVR-Pia (right). Autofluorescence under UV light is shown. (B) Pias-1 cooperates with RGA5 to recognize AVR-Pia and induces resistance in rice. The rice line HW-RIL7 with *Pias* (*Pias-1* and *Pias-2*) recognizes the Ao-92-06-2 strain with *AVR-Pias* (Ao92-06-2+*pex22p*:*AVR-Pias*) and induces resistance. However, HW-RIL7 cannot recognize the Ao02-06-2 strain with *AVR-Pia* (Ao92-06-2+*pex22p*:*AVR-Pia*). Two lines (Line #1 and #2) contain the *35S*-*RGA5* transgene in the HW-RIL7 background. F1-A is a progeny derived from a cross between Sasanishiki with *Pia* (*RGA4* and *RGA5*) and HW-RIL7. F1-B is a progeny derived from a cross between a Sasanishiki mutant (Sas1493) with *pia* (*rga4* and *RGA5*) and HW-RIL7.

Finally, we generated transgenic rice plants (HW-RIL7:35S-RGA5) expressing the *RGA5* transgene driven by the 35S cauliflower mosaic virus (CaMV35S) promoter in the *Pias* (*Pias-1* and *Pias-2*) background. We challenged these lines with two *M. oryzae* isolates: Ao92-62-2 harboring the *AVR-Pia* transgene and Ao92-62-2 harboring the *AVR-Pias* transgene (Fig. 4B; Suppl. Fig. 18). Remarkably, HW-RIL7:35S-RGA5 exhibited resistance against *M. oryzae* containing *AVR-Pia* (Fig. 4B). Although the HW-RIL7:35S-RGA5 lines consistently showed resistance against *M. oryzae* containing *AVR-Pia*, their resistance against *M. oryzae* containing *AVR-Pias* varied. We crossed an *rga4* mutant in the Sasanishiki background (Sas1493; Okuyama et al. 2011) with HW-RIL7 and obtained F1-B plants. These plants, harboring intact *RGA5* as well as *Pias-1* and *Pias-2*, now recognized and triggered resistance against *M. oryzae* containing *AVR-Pia*, but not *AVR-Pias* (Fig. 4B). Similar results were obtained for F1-A plants generated by crossing Sasanishiki (WT) with HW-RIL7 (Fig. 4B). These results suggest that *Pia* function is dominant over *Pias* function in terms of the recognition of *AVR-Pia* and *AVR-Pias*.

These results suggest that Pias-1 helper functions together with RGA5 in rice to recognize and mount resistance against *M. oryzae* containing *AVR-Pia*, suggesting that the helper function has been conserved over the long history of Pias/Pia evolution.

## Discussion

Our study investigated the evolution of a pair of genetically linked NLRs in the genus *Oryza* and provided experimental evidence that the two paired NLRs have evolved in dramatically contrasting fashions. This study points to the evolution of a modular architecture of paired NLRs. Division of roles between a conserved helper NLR for signaling and a divergent sensor NLR with a cassette-like receptor domain for pathogen sensing may have given plants the ability to efficiently fend off rapidly evolving microbe pathogens.

We identified and functionally characterized the rice R-gene *Pias*. This gene encodes the paired NLRs Pias-1 helper and Pias-2 sensor, which recognizes the *M. oryzae* effector AVR-Pias. *Pias* is allelic to the well-studied *R*-gene *Pia*, encoding the NLRs RGA4 helper and RGA5 sensor (Okuyama et al. 2011), which recognizes the effectors AVR-Pia (Yoshida et al. 2009) and AVR1-CO39 (Cesari et al. 2013). The allelic sensor-NLRs Pias-2 and RGA5 carry different domains at their C termini. In RGA5, the integrated domain HMA directly binds to and recognizes two *M. oryzae* effectors, AVR-Pia and AVR1-CO39 (Cesari et al. 2013). We have not yet detected direct binding between the DUF761 ID of Pias-2 and AVR-Pias despite several attempts. Perhaps the recognition of AVR-Pias by Pias-2 requires other host components; indeed, the recognition of AVR-Pii by Pii-2 requires the rice protein OsExo70-F3, which binds to both AVR-Pii and Pii-2 (Fujisaki et al. 2015, 2017).

What is the origin of the integrated DUF761 domain of Pias-2? The rice sensor-NLRs RGA5 and Pik-1 contain HMA domains as IDs. The ID HMA shares high amino acid sequence similarity with rice small heavy metal–associated domain proteins (sHMAs: Oikawa et al. 2020; Białas et al. 2021). We revealed that the *M. oryzae* effector AVR-Pik binds to and stabilizes sHMA proteins, likely to promote pathogen infection (Oikawa et al. 2020; Maidment et al. 2021). To identify the proteins that provide the DUF761 domain to Pias-2 ID, we performed BLAST searches against the rice protein database using the short Pias-2 DUF761 sequence (19 amino acids) as a query (Suppl. Fig.19). This identified 15 proteins with a similarity threshold of E < 10. Most of the 15 proteins contained DUF761 at their C termini. Only a few functional studies of DUF761-containing proteins have been performed. A study in cotton (*Gossypium hirsuta*) showed that GhCFE1A, containing DUF761 and DUF4408 domains, binds to actin proteins and localizes to the endoplasmic reticulum (ER) upon overexpression in *N. benthamiana* (Lv et al. 2015). Knockdown of *GhCFE1A* did not cause any phenotypic changes, while its overexpression led to delayed cotton fiber cell elongation. These results suggest that GhCFE1A is a linker protein that mediates the formation of the ER network and actin cytoskeleton. Arabidopsis *A70*, encoding a DUF761-containing protein, is specifically induced in the incompatible interaction with *Pseudomonas syringae*, but not in the compatible interaction (Truman et al. 2007). Knockout of the Arabidopsis DUF761-containing protein gene *DUF761-1* did not have any phenotypic effects, whereas overexpressing *DUF761-1* altered plant morphology and resulted in a constitutive defense response, leading to enhanced resistance against *P. syringae* (Zhang et al. 2019). These findings suggest that DUF761-containing proteins function in defense, presumably mediated by the actin-ER network. Previous studies of the NLRome of Arabidopsis (Van de Weyer et al, 2019) and NLR-ID of various members of the plant kingdom (Sarris et al. 2016) showed that DUF761 is one of the most common domains integrated into NLR as the ID. In Arabidopsis, DUF761 was integrated into TIR-NLR, and the NLRs with DUF761-ID are in almost all cases paired with helper NLRs (Van de Weyer et al, 2019). These findings suggest that the DUF761 domain is a major target of pathogen effectors and has been frequently integrated into NLR. Future studies should investigate how AVR-Pias is recognized by Pias and how AVR-Pias interferes with host cellular processes by its possible interaction with DUF761-containing proteins.

### Divergent sensor NLRs in the Pias/Pia lineage

The Pias-2/RGA5 sensor-NLR lineage is extremely divergent among *Oryza* species, with up to six different ID motifs integrated at their C termini (Fig. 2). These hugely divergent IDs may mediate the detection of a diversity of effectors from the blast fungus and possibly other pathogens. We hypothesize that the diversity of the Pias-2/RGA5 lineage has been maintained by natural selection to maintain various IDs that recognize the invasion of pathogens by directly binding to effectors or guarding host factors that are modified by effectors.

Within *Oryza* species, there are two major clades, C1 and C2. Notably, alleles from both clades are maintained within the species *O. sativa*, *O. rufipogon*, *O. barthii*, *O. meridionalis*, and *O. punctata*. The observed trans-species allelic divergence and their roles in detecting pathogen molecules are similar to those of the major host incompatibility (MHC) locus of vertebrates (Edwards and Hedrick, 1998). Individuals with higher heterozygosity at the MHC locus might have higher fitness (overdominance) due to their ability to bind to a larger number of pathogen peptides (Takahata and Nei, 1990; Piertney and Oliver, 2006). Perhaps in the ancestral outcrossing *Oryza* species, heterozygous plants with a larger repertoire of NLRs with different IDs had a selective advantage against a multitude of pathogens. It is also possible that frequency-dependent selection helped maintain this polymorphism. When the frequency of a pathogen effector in a population increases, the frequency of an allele for a cognate sensor NLR will increase, resulting in a reduction in the frequency of pathogen alleles in the population. In turn, the frequency of another effector gene will increase in the pathogen, and the frequency of the cognate sensor NLR will increase. According to this Red Queen model, the allele frequencies of both effector and sensor NLRs oscillate and may be maintained for a long time by balancing selection (Takahata and Nei, 1990; Woolhouse et al. 2002; Terauchi and Yoshida, 2010). In summary, the highly divergent evolution of Pias/Pia sensor NLRs with variable IDs seems driven by the fitness gain obtained by an enhanced recognition capability of pathogen effectors.

### Genetic mechanism of ID switching in Pias/Pia NLRs

The Pias/Pia sensor NLRs contain various ID sequences at the identical position downstream of the LRR domain. This suggests the presence of a mechanism for “cassette”-like exchange of IDs between sensor NLRs. Between the LRR and ID, we identified a highly conserved stretch of 145–amino acid sequences named LII. Bailey et al. (2018) reported a 43–amino acid CID motif conserved in the region between the LRR and ID of the MIC1 NLR clade by studying the NLRs of seven Poaceae species. The LII motif identified encompasses the CID motif (Suppl. Fig. 10) but extends to the N-terminal direction by approximately 100 amino acids. Bailey hypothesized that the CID could serve as a recombination point of integration of genomic sequences. Despite the difference in the conserved motif, our data basically support Bailey (2018)’s hypothesis that this region serves as the recombination point for the integration of endogenous sequences matching various protein domains. In support of this idea, the downstream sequence of a Pias sensor NLR with DUF761 contains a DNA sequence similar to the LII and ID regions of the *O. punctata* Pias sensor homolog. Perhaps the PKc_M ID of the original *O. punctata* Pias was replaced by the DUF761 ID, and this switch caused the translocation of PKc_M to the region downstream of the sensor NLR gene (Fig. 2F). In view of the high sequence conservation between *O. punctata* PKc_M ID and the downstream sequence, it is also possible that the downstream sequence was functional until the recent past, the Pias sensor contained the dual IDs DUF761 and PKc_M in the same molecule, or Pias switched between these two IDs, possibly via alternative splicing. Future studies should address the mechanism of LII-mediated recombination.

### Function of *Pias/Pia* paired NLRs

Pia has been extensively studied (Césari et al. 2014) and serves as a paradigm for paired NLRs together with the paired NLRs RPS4/RRS1 (Williams et al. 2014). In these two cases, the helper NLR is regarded as a cell death inducer and the sensor NLR as a suppressor that maintains the complex in an inactive state when the pathogen is absent. Once the sensor has been modified by direct binding (RGA5) or modification (RRS1) of the ID by pathogen effectors, the suppression of the helper is released and HR-like cell death occurs. However, Pias NLRs do not function according to this model. In the *N. benthamiana* assay, Pias-1 functioned as a weak cell death inducer and Pias-2 did not suppress Pias-1-mediated cell death. Indeed, Pias-1 and Pias-2 together triggered stronger cell death than that caused by Pias-1 alone. A recent functional study of rice Pikp, another paired NLR, also showed that the helper NLR alone does not cause cell death in the *N. benthamiana* system and that the helper and sensor cooperate to trigger HR (Zdrzałek et al. 2020). Therefore, the hypothesized functional roles of the helper as a cell death inducer and the sensor as a cell death suppressor as well as a detector of effector molecules may not be as prevalent as assumed. The system of negative regulation of a cell death inducer by a suppressor encoded by genetically separate factors carries tremendous risks given that a loss-of-function mutation in the suppressor gene kills the carrier cells and incurs a genetic load. Therefore, such an extreme negative regulation system is unlikely to be maintained over a long period of evolution. We predict that cooperative NLRs in pairs or networks are more prevalent (Wu et al. 2017). It is also possible that the Pias/Pia-paired NLR system is regulated by additional components in rice cells that are absent from *N. benthamiana*. It would be interesting to experimentally determine what proportions of paired NLRs function in negative regulation and in cooperation using *N. benthamiana* transient expression assays. Further studies are needed to decipher the full regulatory network of Pias/Pia-paired NLR-mediated resistance.

### Helper NLRs in the Pias/Pia lineage are functionally conserved

Phylogenetic reconstruction of the Pias/Pia NLR locus revealed that the helper NLR Pias-1/RGA4 is conserved, whereas the sensor NLR Pias-2/RGA5 is highly divergent (Fig. 3). An Arabidopsis NLRome study showed that some NLR pairs coevolved, with the phylogenetic trees of helper and sensor NLRs corresponding (Van de Weyer, 2019). Our findings for Pias/Pia NLRs do not align with these observations. A functional study in *N. benthamiana* showed that Pias-1-mediated cell death was suppressed by RGA5 and that Pias-1 together with RGA5 function in the recognition of AVR-Pia, leading to cell death (Fig. 4A). Similarly, an RGA4 homolog of *O. granulata* that is phylogenetically distant from *O. sativ*a also functions with RGA5 to recognize AVR-Pia (Fig. 4A). Moreover, a HW-RIL7 rice line harboring *Pias* as well as an *RGA5* transgene recognized AVR-Pia (Fig. 4B). These results suggest that the function of the Pias-1/RGA4 helper lineage is conserved, which is in line with its conserved amino acid sequences. It is possible that the separation of the roles of NLRs between the conserved helper and divergent sensor allowed for higher flexibility of pathogen recognition compared to singleton NLRs. This flexibility would allow the plant to cope with the rapid evolution of pathogens, which exhibit larger population sizes and shorter generation times than the host plants. The functional understanding of the modular structure of paired NLRs revealed here provides a basis for engineering NLRs to detect various effectors and to confer resistance to crops against pathogens.

## Materials and Methods

### Rice pathogenicity assays

Rice leaf blade punch inoculation was performed using the *M. oryzae* isolates. A conidial suspension (3 × 10^5^ conidia mL^−1^) was punch inoculated onto a rice leaf 1 month after seed sowing. The inoculated plants were placed in a dew chamber at 27°C for 24 h in the dark and transferred to a growth chamber with a 16-h-light/8-h-dark photoperiod. Disease lesions were scanned 10 days post-inoculation (dpi), and lesion size was measured using Image J software (Schneider et al. 2012).

### RNA-seq of rice and barley leaves infected with *M. oryzae* 2012-1 isolate

Total RNA was extracted from rice and barley infected leaves using an SV Total RNA Isolation System (Promega, WI, USA). One microgram of total RNA was used to prepare each sequencing library with an RNA Sample Prep Kit v. 2 (Illumina, CA, USA). The two types of libraries, created from infected barley and rice leaves, were sequenced by paired-end (PE) and single-end (SE) sequencing using the NextSeq 500 platform.

### DNA-seq for the RaIDeN pipeline

Genomic DNA was extracted from WRC17, Hitomebore, and S-RIL leaves using a NucleoSpin Plant II Kit (Macherey Nagel Co, Düren, Germany). Libraries for PE short reads were constructed using an Illumina TruSeq DNA LT Sample Prep Kit (Illumina, CA, USA). The PE library of WRC17 and other libraries were sequenced on the Illumina MiSeq and HiSeq 4000 platforms.

### Generation of *PiW17-1* with knocked-down expression of candidate NLR genes in rice and quantitative RT-PCR

The eight types of gene knockdown (RNAi) constructs (pANDA-*PiW17-1* candidate NLRs) were generated by PCR amplification of a specific fragment of each *PiW17-1* candidate NLR gene from WRC17 WT cDNA. The sequences were cloned into the Gateway vector pENTR/D-TOPO (Invitrogen, CA, USA) and transferred into recombination sites of the pANDA vector (Miki and Shimamoto, 2004) using LR Clonase (Invitrogen, CA, USA). The resulting vectors with eight types of pANDA-*PiW17-1* candidate NLR genes were introduced into *Agrobacterium tumefaciens* (strain EHA105) and used for *A. tumefaciens–*mediated transformation of HW-RIL7 following the method of Okuyama et al. (2011). Total RNA was extracted from leaves using an SV Total RNA Isolation System (Promega, WI, USA) and used for quantitative RT-PCR (qRT-PCR). cDNA was synthesized from 500 ng total RNA using a Prime Script RT Reagent Kit (Takara Bio, Otsu, Japan). qRT-PCR was performed using a StepOne Real-time PCR Instrument (Applied Biosystems, CA, USA) with KAPA SYBR FAST PCR Master Mix (Kapa Biosystems, MA, USA). Melting curve analysis (from 60 to 95°C) was included at the end of the cycles to ensure the consistency of the amplified products. The comparative Ct (ΔΔCt) method was used to calculate the expression of *CNL-04* (*CNL-05*) relative to the rice *ACTIN* gene (*LOC_Os03g50885*) as an internal control. The data presented are the average and standard deviations from three experimental replications. The primers used to generate the RNAi construct and for qRT-PCR are listed in **Suppl. Table 10**.

### Generation of rice mutants of *CNL-04* and *CNL-05* by CRISPR/Cas9-mediated genome editing

Rice knockout mutants of *CNL-04* and *CNL-05* were generated using the CRISPR/Cas9 system developed by Mikami et al. (2015). Sense and antisense target sequences were designed using the web-based service CRISPR direct (http://crispr.dbcls.jp), annealed, and cloned into the pU6::ccdB::gRNA cloning vector following digestion with *Bbs*I as the target sequence. The target sequence with the *OsU6* promoter was cloned into the pZH::gYSA::MMCas9 vector following digestion with *Asc*I and *Pac*I. The resulting vectors (pZH::gYSA::MMCas9-*CNL-04* and -*CNL-05*) were introduced into *A. tumefaciens* (strain EHA105) and used for *A. tumefaciens–*mediated transformation of HW-RIL7 following the method of Okuyama et al. (2011). The resulting regenerated T0 plants were sequenced, and the mutation type was confirmed using primers listed in **Suppl. Table 10**.

### Genetic complementation of the candidate AVR-Pias

Three candidate gene constructs (pCB1531-*AVR-Pias* candidate) were generated by PCR amplification of the coding sequences of the *AVR-Pias* candidate genes from cDNA prepared from *M. oryzae* 2012-1-infected barley leaves. The sequences were digested with *Xba*I and *BamH*I and cloned into pCB1531-pex22p-EGFP (Yoshida et al. 2009) that had been linearized by digestion with *Xba*I and *Bam*HI. The resulting vectors were used to transform Ao92-06-2 (lacking *AVR-Pias*) following the method of Sweigard et al. (1997). *M. oryzae* isolate 2012-1 mutated in *G9532* was generated using the CRISPR/Cas9 system developed by Arazoe et al. (2015). Sense and antisense target sequences were designed using the web-based service CRISPR direct (http://crispr.dbcls.jp), annealed, and cloned into the pCRISPR/Cas-U6-1 cloning vector following the method of Arazoe et al. (2015). To generate the targeting vector TV-*G9532*, the 5ʹ flanking region of *G9532* was amplified and cloned into pCB1636 (Sweigard et al. 1997) containing a hygromycin resistance gene that had been linearized by inverse PCR using primers pCB1636iv2fwd and pCB1636iv2rev as described by Shimizu et al. (2019) using in-fusion cloning (Clontech, Madison, WI, USA). Subsequently, the 3′ flanking region was amplified and cloned into a plasmid containing the 5′ flanking region that had been linearized by inverse PCR using primers pCB1636iv1fwd and pCB1636iv1rev as described by Shimizu et al. (2019) using in-fusion cloning (Clontech, Madison, WI, USA). The resulting vectors were used to transform the 2012-1 isolate (containing *AVR-Pias*) following the method of Sweigard et al. (1997). The primers used for construct generation are listed in **Suppl. Table 10**.

### Genome sequences used for the study

Genome sequences of 171 accessions of Poaceae plants were used, including 167 *Oryza* accessions as well as one accession each from *Aegilops tauschii*, *Hordeum vulgare*, *Panicum hallii*, and *Setaria italica*. These also included sequences of *O. sativa* ‘WRC17’ (this study) and *O. sativa* ‘Sasanishiki’ (Okuyama et al. 2011). Genome sequences of 66 accessions were obtained from public databases: 52 accessions of *O. sativa* and *O. rufipogon* (Zhao et al. 2018), *O. sativa* subsp. *indica* ‘ShuHui498’ (Du et al. 2017), *O. sativa* subsp. *japonica* ‘Nipponbare’ (IRGSP. 2004), *O. rufipogon* ‘W1943’ (National Center for Gene Research), 5 accessions from *O. barthii*, *O. glumaepatula*, *O. meridionalis*, *O. punctata*, and *O. brachyantha* (The Oryza Map Alignment Project (OMAP)), *O. officinalis* ‘W0002’ (National Institute of Genetics, Japan), *O. granulata* ‘W0067B’ (Wu et al. 2018), *Ae. tauschii* ‘AL8/78’ (Zimin et al. 2017), *H. vulgare* ‘Morex’ (Mascher et al. 2017), *P. hallii* ‘FIL2’ (DOE Joint Genome Institute), and *S. italica* ‘Yugu18’ (Bennetzen et al. 2012) (see **Suppl. Table 6** for details). For 101 accessions of the wild *Oryza* species (*O. barthii*, *O. glumaepatula*, *O. meridionalis*, *O. punctata*, *O. officinalis, O. brachyantha*, and *O. granulata*), NGS reads (fastq format) of whole-genome sequences were retrieved from the NGS NCBI database (**Suppl. Table 6**) and used for *de novo* assembly by MaSuRCA (Zimin et al. 2013). For the two wild *Oryza* accessions, *O. glumaepatula* W2184 and *O. punctata* W1582, DNA sequencing was performed using Oxford Nanopore Technology (ONT) using genomic DNA extracted from their leaves. Sequencing was performed using the PromethION system with a FLO-PRO002 flow cell (ONT). Base calling of ONT reads was performed on FAST5 files using Guppy (ONT). Subsequently, low-quality reads were filtered out, and *de novo* assembly was performed using NECAT software (https://github.com/xiaochuanle/NECAT/). To further improve the accuracy of the assembly, Racon software (https://github.com/lbcb-sci/racon) was applied twice, and Medaka (https://github.com/nanoporetech/medaka) was used to correct mis-assembly. One round of consensus correction was performed using BWA (Li and Durbin, 2010) and HyPo (https://github.com/kensung-lab/hypo) on Illumina short reads for the accessions.

### Gene annotation and detection of Pias/Pia orthologs

To infer the protein coding regions of *Pias-1/RGA4* and *Pias-2/RGA5* genes, and to obtain information about their ID sequences, we used the pipeline shown in Suppl. Fig. 9. We retrieved *Pias-1*/*RGA4* and *Pias-2* /*RGA5* gene models from the genome sequences and RNA-seq data publicly available for seven *Oryza* samples (*O. barthii* ‘IRGC105608’, *O. glumaepatula* ‘GEN1233_2’, *O. meridionalis* ‘W2112’, *O. punctata* ‘IRGC105690’, *O. australiensis* ‘W0008’, *O. brachyantha* ‘IRGC101232’, and *O. granulata* ‘W0067B’). For *O. rufipogon* ‘W1943’ and *O. officinalis* ‘W0002’, genome sequences were publicly available, but we performed new RNA-seq analyses to improve gene prediction. For the sample *O. punctata* ‘W1582’, we performed genome sequencing and RNA-seq analyses. We also used the *RGA4/RGA5* gene model of *O. sativa* cv. Sasanishiki (Okuyama et al. 2011) and the *Pias-1/Pias-2* gene model of *O. sativa* cv. Keiboba (this study). The gene models of *Pias-1/RGA4* and *Pias-2/RGA5* of these 12 *Oryza* samples were used as queries to annotate IDs in the genome assembly of 167 *Oryza* samples using Exonerate (http://www.ebi.ac.uk/~guy/exonerate). However, 10 samples of *O. glumaepatula* and six samples of *O. brachyantha* did not match to known domains. Therefore, we incorporated RNA-seq data for each sample for the two species, resulting in the annotation of the Zinc_ribbon_12 (*O. glumaepatula*) and HMA (*O. brachyantha*) IDs. In the next round, we used the gene models of 12 samples used in the first round of Exonerate as well as two new samples (*O. glumaepatula* W2184 and *O. brachyantha* W0655) as queries to infer IDs in the assembled genomes of 167 *Oryza* samples (**Suppl. Table 6, 7, 8**). This resulted in the identification of five known domains and one unknown domain in the *Oryza* ID sequences.

### Amino acid sequences and accession numbers of Pias/Pia orthologs

Amino acid sequences of Pias/Pia homologs used in this study were retrieved from the Eukaryotic Genome Annotation of the NCBI database (Du et al. 2017; Mascher et al. 2017); accession numbers for Pias-1/RGA4 homologs and Pias-2/RGA5 homologs are as follows: XP_015617251.1 and XP_015617810.1 for *O. sativa* subsp. *japonica* ‘Nipponbare’, OsR498G1119642600.01 and OsR498G1119642700.01 for *O. sativa* subsp. *indica* ‘ShuHui498’, XP_020148260.1 and XP_020148256.1 for *Aegilops tauschii* ‘AL8/78’, HORVU.MOREX.r2.4HG0288000.1 and HORVU.MOREX.r2.4HG0288010.1 for *Hordeum vulgare* ‘Morex’, XP_025826635.1 and XP_025827327.1 for *Panicum hallii* ‘FIL2’, and XP_004979045.1 and XP_004979046.2 for *Setaria italica* ‘Yugu18’.

### Phylogenetic analysis of Pias-1/RGA4 and Pias-2/RGA5 orthologs

The protein sequences of the Pias-1/RGA4 and Pias-2/RGA5 orthologs were aligned using webPRANK (https://www.ebi.ac.uk/goldman-srv/webprank/) (Löytynoja et al. 2010). We used the whole amino acid sequences of the Pias-1/RGA4 orthologs but only the partial amino acid sequences (CC-NB-LRR-LII domains) of the Pias-2/RGA5 orthologs due to the very low sequence similarity after the LII domains. A maximum likelihood tree was constructed with IQ-TREE v2.0.3 (Minh et al. 2020) using 1,000 ultrafast bootstrap replicates (Hoang et al., 2018). The models were automatically selected by ModelFinder (Kalyaanamoorthy et al. 2017) in IQ-TREE (Minh et al. 2020). ModelFinder (Kalyaanamoorthy et al. 2017) selected “JTT + G4” for Pias-1/RGA4 orthologs and “JTT + R2” for Pias-2/RGA5 orthologs. Finally, the phylogenetic trees were drawn with FigTree v1.2.3 (http://tree.bio.ed.ac.uk/software/figtree/).

### Analysis of DNA polymorphisms, *dN*, and *dS*

The coding DNA sequences (CDSs) of the *Pias-1/RGA4* and *Pias-2/RGA5* orthologs were aligned using the codon-based aligner MACSE v2.05 (Ranwez et al. 2018). We applied MACSE v2.05 not only to the entire CDS of *Pias-1/RGA4* and CC-NBS-LRR-LII of *Pias-2/RGA5* but also to each domain (CC, NBS, and LRR[-LII]) using default parameters. We evaluated DNA polymorphisms of the *Pias-1/RGA4* and *Pias-2/RGA5* orthologs, calculating the nucleotide diversity (π) and Tajima’s *D* (Tajima, 1989) using MEGAX v10.2.4 (Kumar et al. 2018). Then, for each alignment, the maximum likelihood trees were constructed using IQ-TREE v2.0.3 with 1,000 ultrafast bootstrap replicates (Hoang et al., 2018) and ModelFinder (Kalyaanamoorthy et al. 2017). Based on these alignments and trees, we calculated the pairwise *dN* and *dS* using the YN00 program (Yang and Nielsen, 2000) in PAML v4.8 (Yang 1997).

### Expression constructs used in the cell death assay

Expression constructs for five types of helper-NLRs, Pias-1 (*O. sativa* subsp. *Indica*, WRC17), RGA4 (*O. sativa* subsp. *Japonica*, Sasanishiki), RGA4-Oru (*O. rufipogon* accession W1943), RGA4-Oau (*O. australiensis* accession W0008), and RGA4-Ogr (*O. granulata* accession W0067B) (pCambia1300S-“helper-NLR”:HA), were generated by PCR amplification of the coding sequences from cDNA generated from leaf material and cloned into the binary vector pCambia1300S (http://www.cambia.org) that had been linearized by digestion with *Pst*I and *Spe*I by In-fusion cloning. Expression constructs for two types of sensor NLRs, Pias-2 (*O. sativa* subsp. *Indica*, WRC17) and RGA5 (*O. sativa* subsp. *Japonica*, Sasanishiki) (pCambia1300S-FLAG: “sensor NLR”), were generated by PCR amplification of the coding sequences from cDNA generated from leaf material and cloned into the binary vector pCambia1300S (http://www.cambia.org) that had been linearized by digestion with *Sal*I and *Pst*I by In-fusion cloning. The resulting vectors were introduced into *A. tumefaciens* (strain GV3101). The primers used to generate the expression constructs are listed in **Suppl. Table 10**.

### Immunoblot analysis

Proteins were expressed in *Nicotiana benthamiana* leaves and extracted from leaf tissue (approximately 100 mg) in 200 µL of extraction buffer (50 mM Tris-HCl [pH 7.5], 150 mM NaCl, and 0.1 % NP-40 [v/v]). The extracts were subjected to SDS-PAGE, followed by immunoblotting using anti-FLAG-HRP antibody, anti-HA-HRP antibody, or anti-AVR-Pia antibody. Anti-mouse IgG-HRP was used as a secondary antibody following incubation with anti-AVR-Pia antibody. Antibody-antigen complexes were detected using a luminescent image analyzer (ImageQuant LAS-4000) (Cytiva, Tokyo, Japan). Equal loading of proteins in the PAGE gel was confirmed by Coomassie blue staining.

### Cell death assay in *N. benthamiana*

Transient expression of Pias/Pia allelic NLR and AVR-Pia was performed by infiltrating 4- to 5-week-old *N. benthamiana* plants with *A. tumefaciens* carrying the expression vector. *A. tumefaciens* suspensions in infiltration buffer (10 mM MES, 10 mM MgCl_2_, and 150 µM acetosyringone, pH 5.6) were adjusted to the densities shown in **Suppl. Table 11**. The autofluorescence value under UV light was scored using a luminescent image analyzer (ImageQuant LAS-4000) (Cytiva, Tokyo, Japan).

### Yeast two-hybrid assay

To examine the protein–protein interactions between Pias-1:CC_182_ and Pias-2:CC_177_, a yeast two-hybrid assay was performed as described previously (Kanzaki et al. 2012). The CC domain of Pias-1 (Pias-1:CC_182_) was amplified by PCR, digested with *Eco*RI and *Bam*HI, and cloned into pGADT7 (prey) or pGBKT7 (bait) vector (Clontech, Madison, WI, USA) that had been linearized by digestion with *Eco*RI and *Bam*HI by in-fusion cloning. The CC domain of Pias-2 (Pias-2:CC_177_) was amplified by PCR, digested with *Sfi*I, and cloned into pGADT7 or pGBKT7 vector that had been linearized by digestion with *Sfi*I. GFP (as negative control) was amplified by PCR and cloned into pGADT7 or pGBKT7 vectors that had been linearized by digestion with *Eco*RI and *Bam*HI by in-fusion cloning. The various combinations of bait and prey vectors were transformed into yeast strain AH109 using the PEG/LiAc method. To detect protein–protein interactions, a 10-fold dilution series (×1, ×10^-1^, ×10^-2^) of yeast cells (×1: OD_600_ = 1.0) was spotted onto basal medium lacking Trp, Leu, Ade, and His (-HTLA) but containing 5-bromo-4-chloro-3-indolyl α-D-galactopyranoside (Clontech, Madison, WI, USA). As a control, yeast growth on basal medium lacking Trp and Leu (–TL) was also checked. To assess the protein accumulation in yeast cells, each transformant was propagated in liquid basal medium lacking Trp and Leu with gentle shaking at 30°C overnight. Yeast cells from 10 mL medium were collected, and 100 mg of yeast cells was treated with 400 µL of 0.3 N NaOH for 15 min at room temperature. The resulting yeast extracts were used for immunoblot analysis using anti-Myc (MBL, Woburn, MA, United States) for bait proteins and anti-HA (Roche, Switzerland) for prey proteins. The primers used to generate the constructs are listed in **Suppl. Table 10**.

### Co-immunoprecipitation assay

Expression constructs for two types of helper-NLRs, Pias-1 (MHD) and RGA4 (MHD) (pCambia1300S-“helper-NLR”:HA), were generated by PCR and cloned into the binary vector pCambia1300S (http://www.cambia.org) that had been linearized by digestion with *Pst*I and *Spe*I by in-fusion cloning. The resulting vectors were introduced into *A. tumefaciens* (strain GV3101). Pias-1(MHD):HA and FLAG:Pias-2 or FLAG:GFP were transiently co-expressed in *N. benthamiana.* Similarly, RGA4 (MHD):HA and FLAG:RGA5 or FLAG:GFP were transiently co-expressed in *N. benthamiana*. The co-expressed proteins were extracted with 50 mM sodium phosphate buffer containing 10 % (v/v) glycerol and 150 mM NaCl. Extracted proteins were incubated with FLAG-agarose beads in the presence of protease inhibitor, 10 mM DTT, and 2% (w/v) blocking reagent (Cytiva, Tokyo, Japan) at 4°C for 1 h. The beads were washed with the same buffer four times, and then bound proteins were extracted with the same buffer containing 1% (w/v) SDS by boiling. Bound fractions were analyzed by immunoblotting using anti-FLAG and anti-HA antibodies. The primers used to generate constructs are listed in **Suppl. Table 10**.

### Data Availability

The nucleotide sequences of *Pias-1*, *Pias-2*, and *AVR-Pias* were deposited in the DNA Databank of Japan (*Pias-1*: LC672059, *Pias-2*: LC672060, and *AVR-Pias*: LC672061). The DNA-seq and RNA-seq data from this study are listed in **Suppl. Dataset 2** and were deposited in the DNA Databank of Japan (DDBJ) under the Bioproject accession numbers PRJDB9440, PRJDB12353, PRJDB12884, PRJDB12891, and PRJDB12902. All data pertaining to this study are included in the article and/or supporting information.

## Acknowledgements

This study was supported by JSPS KAKENHI 15H05779, 20H00421 and 16H06279 (PAGS) to RT and 21K14834 to MS, the Royal Society UK-Japan International exchange grants JPJSBP120215702 and IEC\R3\203081 to RT, SK and MB, the Biotechnology and Biological Sciences Research Council (UKRI-BBSRC, UK, grant BB/P012574), the European Research Council (ERC BLASTOFF project, 743165), the John Innes Foundation and the Gatsby Charitable Foundation. Computations were partially performed on the NIG supercomputer at ROIS National Institute of Genetics. We also thank the National Institute of Genetics (National Bioresource Project), Japan, and the Genebank at the National Agriculture and Food Research Organization, Japan, for providing the seeds of wild rice and WRC17, respectively.

## Legends to Supplementary Figures

**Suppl. Fig. 1.**
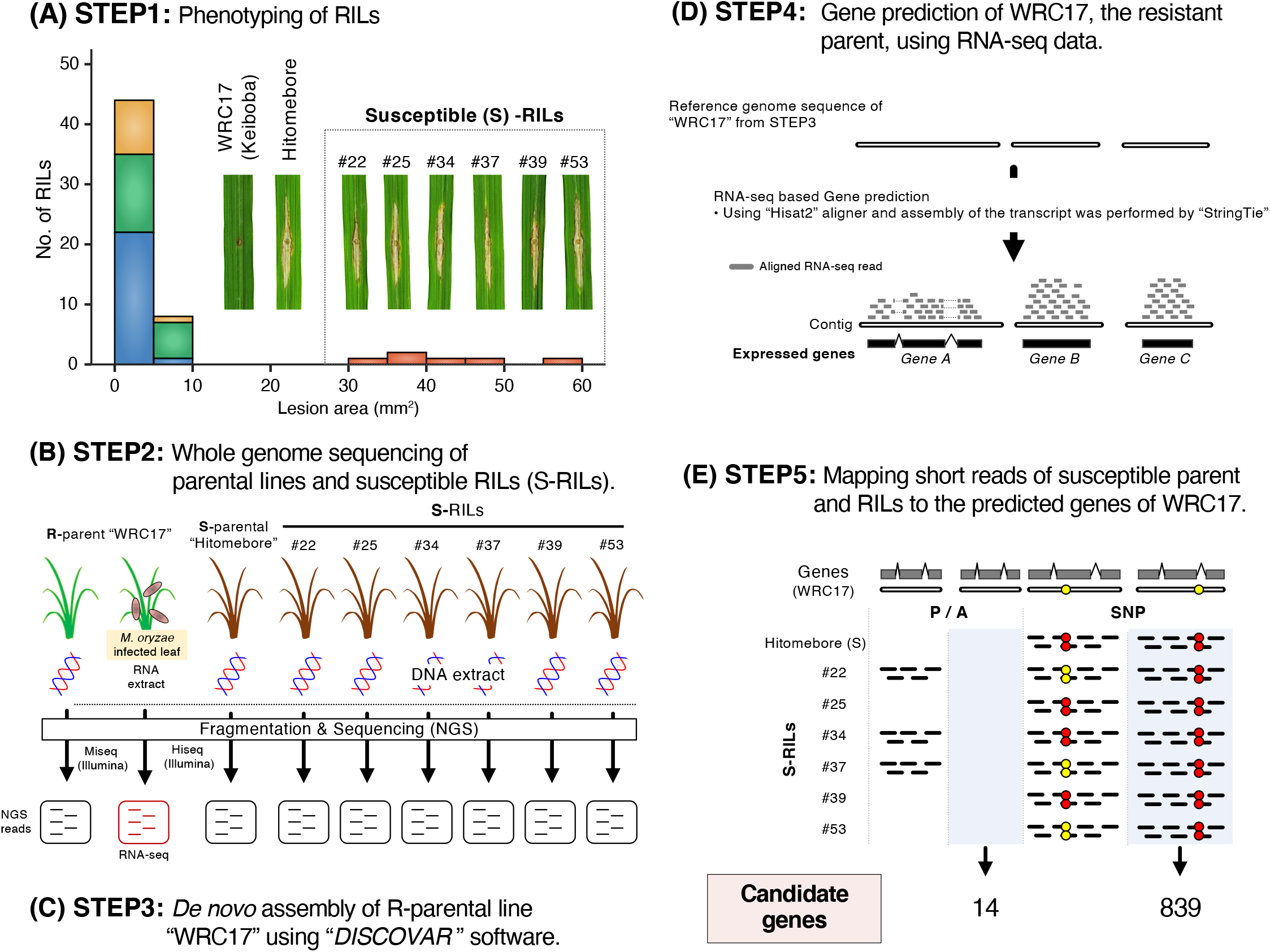
The RaIDeN method used to identify candidate genes responsible for the resistance of rice line WRC17 to *M. oryzae* 2012-1. (A) Segregation of the resistance and susceptibility traits among the 58 RILs derived from a cross between WRC17 (cultivar Keiboba) and Hitomebore. (B) The whole-genome sequence of the resistant parent WRC17 was obtained by Illumina sequencing. RNA-seq of WRC17 leaves inoculated with *M. oryzae* was also performed. Next, short reads of whole-genome sequences of the susceptible parent Hitomebore and the six susceptible RILs were obtained by Illumina sequencing. (C) *De novo* assembly of R-parental line WRC17 was performed using *DISCOVAR* software. (D) Expressed genes of WRC17 were identified by mapping RNA-seq reads to the WRC17 reference genome. (E) Mapping of short reads of the susceptible parent (Hitomebore) and susceptible RILs to the predicted genes identified DNA polymorphisms shared by the susceptible lines using the RaIDeN pipeline (the script of the RaIDeN pipeline and details about the pipeline are available at https://github.com/YuSugihara/RaIDeN).

**Suppl. Fig. 2.**
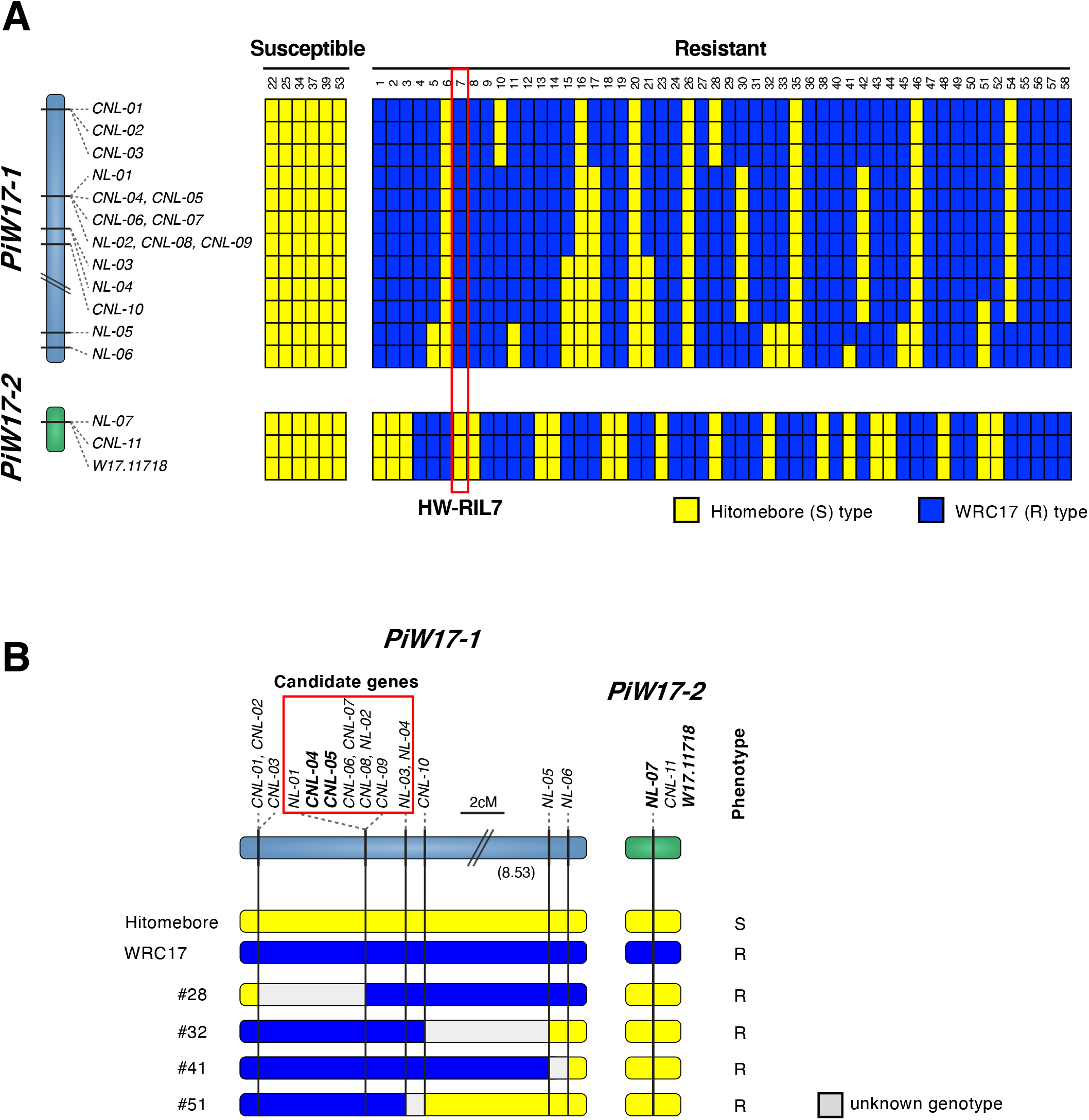
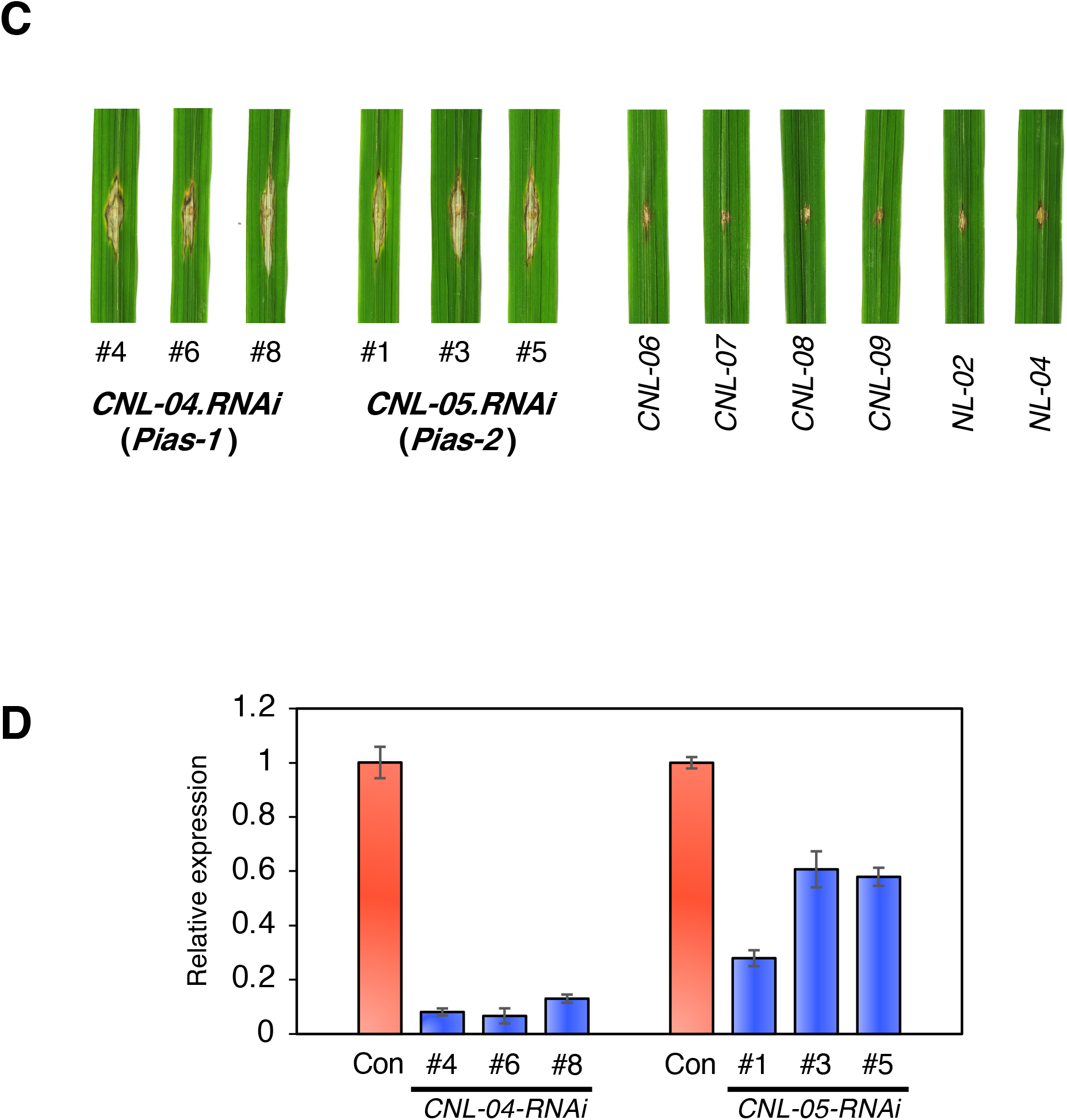
Mapping of *NLR*s in the *PiW17-1* and *PiW17-2* loci that confer resistance against *M. oryzae* 2012-1 to WRC17. (A) Association between resistance/susceptible phenotypes and genotypes among the RILs (the primers used for genotyping are shown in Suppl. Table 9). *CNL-04* and *CNL-05*, *CNL-06* and *CNL-07*, and *CNL-08*, *NL-2*, and *CNL-09* are located on the same contigs. (B) Fine maps of candidate NLR genes in *PiW17-1* and *PiW17-2*. Blue and yellow indicate WRC17-and Hitomebore-type genotypes, respectively. The 10 NLRs shown in the red rectangle were considered to be the candidates of *Pi-W17-1.* **Results of RNAi-mediated silencing of candidate NLR genes.** (C) RNAi-mediated silencing of eight candidate genes encoding proteins over 900 amino acids long was performed. Gene silencing of *CNL-04* and *CNL-05* made plants susceptible to *M. oryzae* 2012-1. (D) Gene expression levels of *CNL-04* and *CNL-05* in plants transformed with the RNAi constructs. Control (Con) is WRC17 transformed with empty vector.

**Suppl. Fig. 3.**
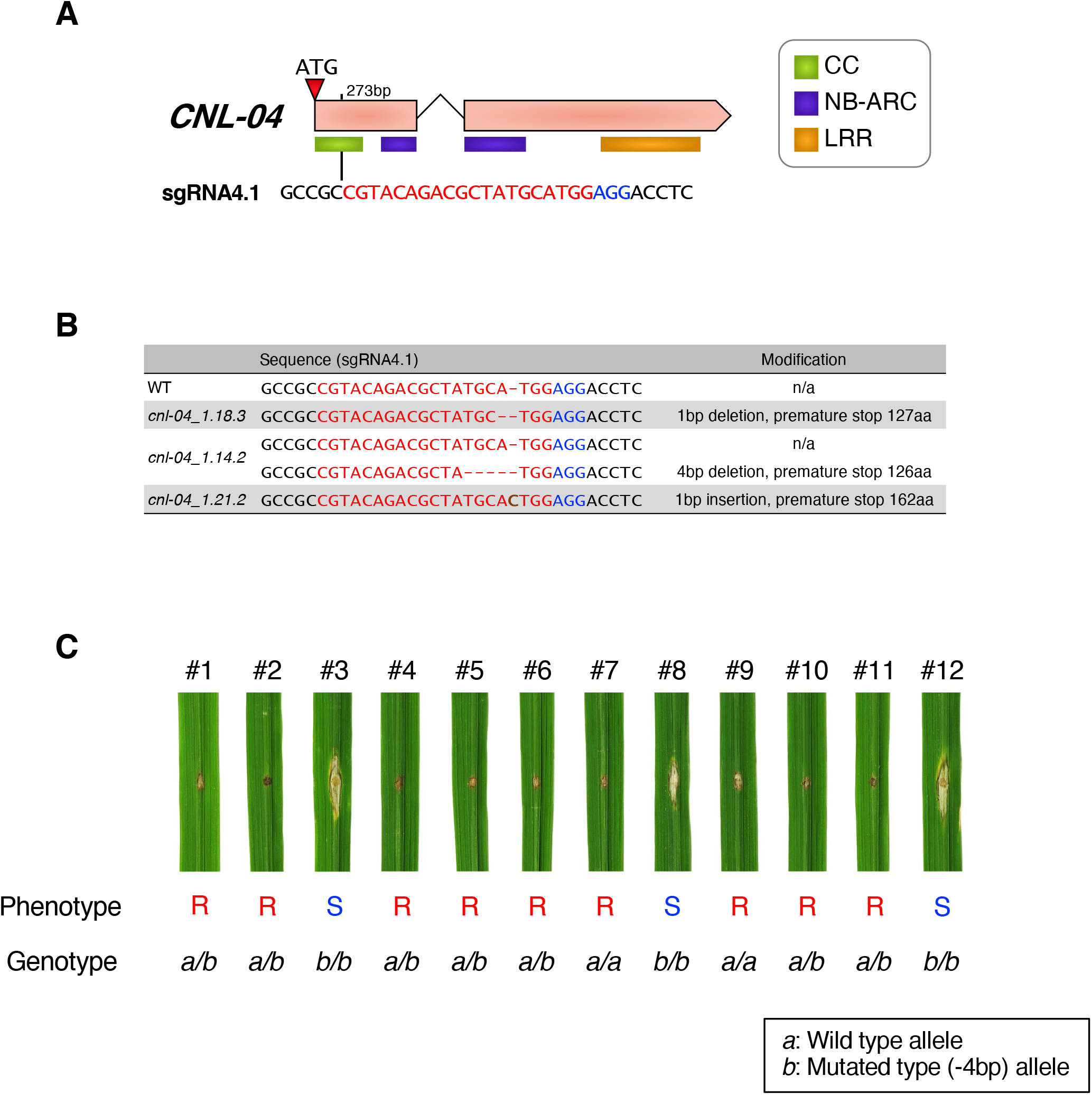
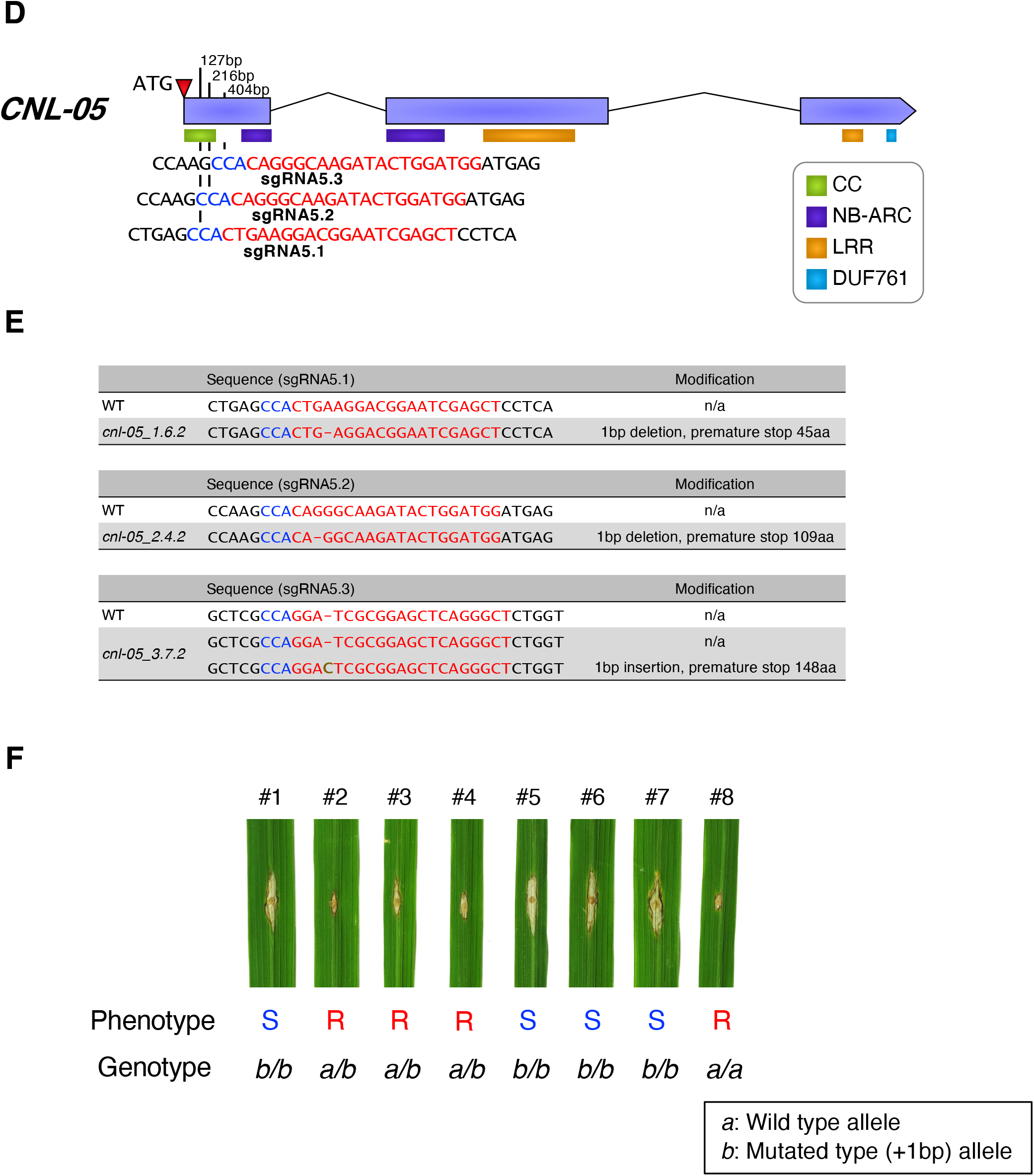
CRISPR/Cas9-mediated knockout of *CNL-04* abolishes *PiW17-1*-mediated resistance against *M. oryzae* isolate 2012-1. (A) The position of guide DNA targeted to *CNL-04*. The PAM is marked with blue letters, and the sgRNA sequence is marked with red letters. (B) Sequences of the sgRNA positions in the *CNL-04* knockout lines. (C) Resistance/susceptible phenotypes of the selfed progeny of the *cnl-04_1.14.2* line heterozygous for wild-type (*a*) and mutated (*b*: 4-bp deletion) alleles. The *b/b* homozygous plants became susceptible. **CRISPR/Cas9-mediated knockout of *CNL-05* abolishes *PiW17-1* mediated resistance against *M. oryzae* isolate 2012-1.** (D) The positions of guide DNAs targeted to *CNL-05*. The PAM is marked with blue letters, and the sgRNA sequence is marked with red letters. (E) Sequences of the sgRNA positions in the *CNL-05* knockout lines. (F) Resistance/susceptible phenotypes of the selfed progeny of *cnl-05_3.7.2* line heterozygous for wild-type (a) and mutated (*b*: 1-bp insertion) alleles. The *b/b* homozygous plants became susceptible.

**Suppl. Fig. 4.**
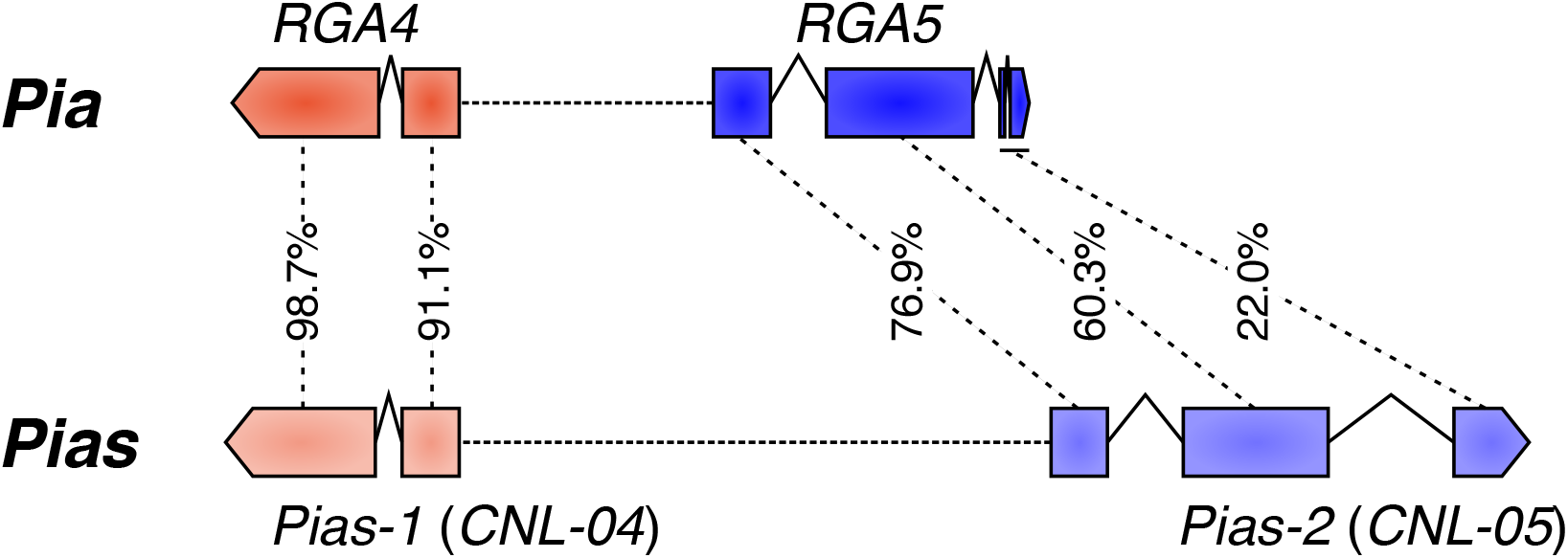
Comparison of the gene structures of *Pia* (*RGA4* and *RGA5*) and *Pias* (*Pias-1* and *Pias-2*). DNA sequence similarities obtained by ClustalW are shown.

**Suppl. Fig. 5.**
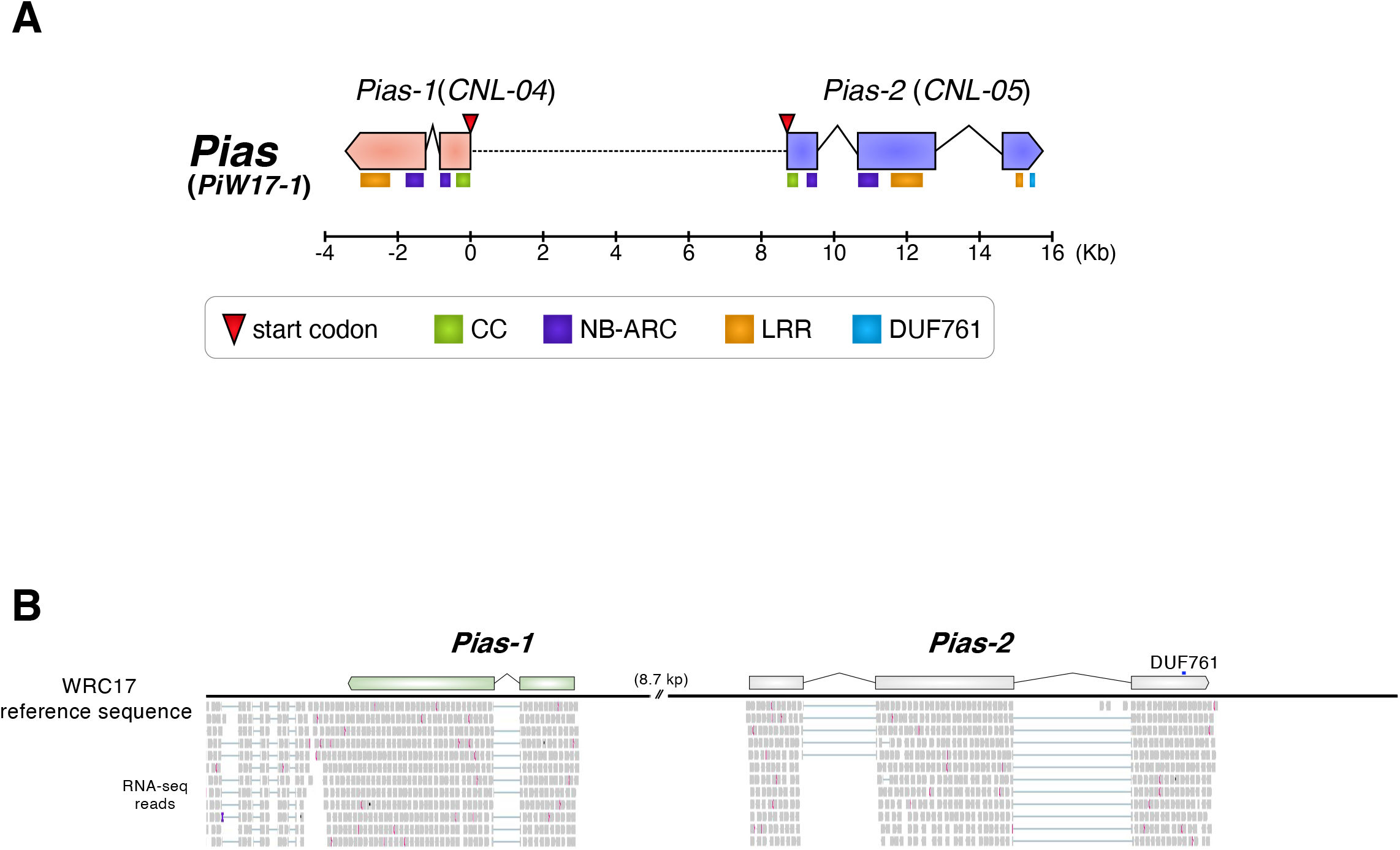
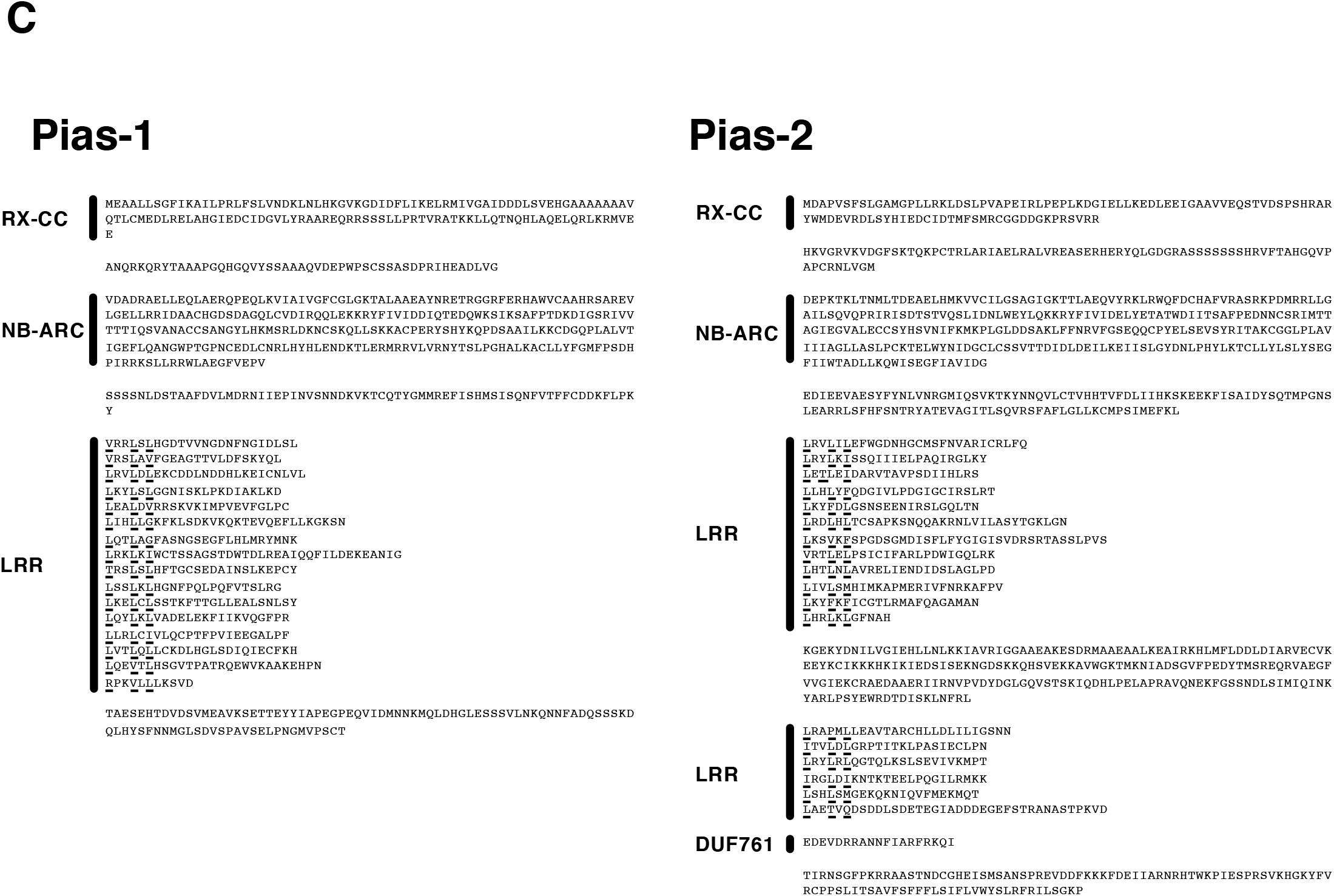
Gene structures and predicted amino acid sequences of *Pias-1* and *Pias-2*. (A) Gene structures of *Pias-1* and *Pias-2*. (B) RNA-seq read alignment across the genome sequence of the *Pias* region as visualized by IGV (Robinson et al. 2011). (C) Amino acid sequences and putative domains of Pias-1 and Pias-2. The RX-CC and NB-ARC domains of Pias-1 and Pias-2 were annotated by a CD-search in NCBI (https://www.ncbi.nlm.nih.gov/Structure/cdd/wrpsb.cgi), and the LRR was predicted using LRR predictor (Martin et al 2020).

**Suppl. Fig. 6.**
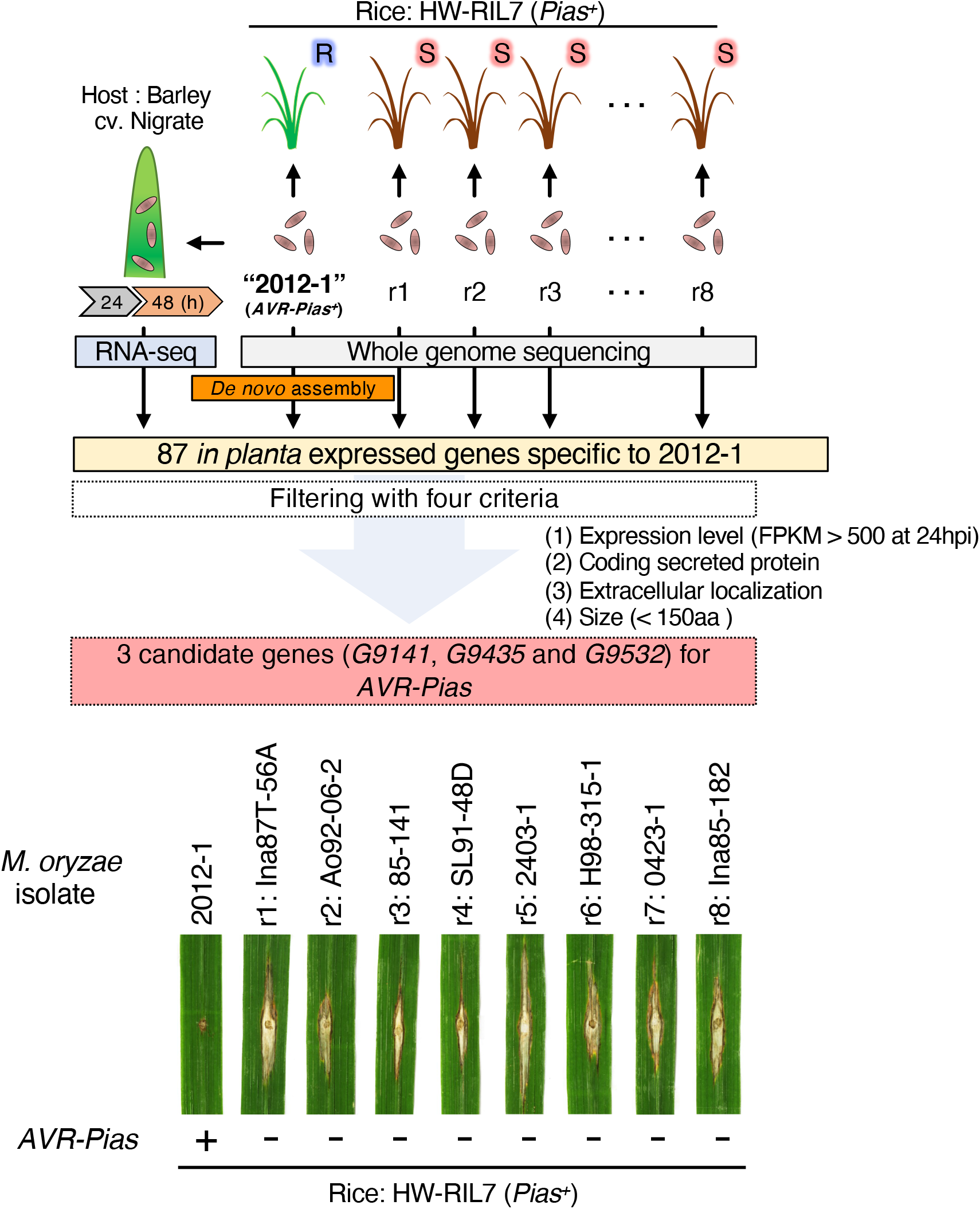
Isolation of *AVR-Pias* from *M. oryzae* isolate 2012-1. A simplified scheme showing the procedure used to isolate *AVR-Pias*. First, the genome of isolate 2012-1 was sequenced on the Illumina platform, and the short reads were used for *de novo* assembly with *DISCOVAR*, resulting in the 2012-1 genome reference (**Suppl. Table 4, 5**). Next, the isolate 2012-1 was used to inoculate barley (*Hordeum vulgare*) cultivar ‘Nigrate’, which is highly susceptible to *M. oryzae* (Hyon et al. 2012), and the infected leaves were subjected to RNA-seq (Shimizu et al. 2019), resulting in the identification of 10,991 genes in 2012-1 that were expressed during host infection. Next, eight *M. oryzae* isolates (r1–r8) were selected and used for inoculation of rice line HW-RIL7 with *Pias* but without *Pi-W17-2*. The eight isolates were compatible with HW-RIL7 (result shown in the bottom), suggesting that they lack *AVR-Pias*. The eight isolates were then subjected to genome resequencing on the Illumina platform, and the RaIDeN method was used to identify only presence/absence polymorphisms (**Suppl. Table 4**). The short reads of eight isolates were aligned to the expressed genes of the 2012-1 isolate, resulting in the identification of 87 expressed genes that were specific to isolate 2012-1. These transcripts were further filtered based on four criteria: (1) genes showing a higher level of expression 24 h after inoculation (FPKM > 200) and genes encoding (2) a putative secreted protein, (3) a non-transmembrane protein, and (4) a small protein (<150 amino acids). This analysis identified three transcripts (*G9141*, *G9435*, and *G9532*) as the candidates of *AVR-Pias*.

**Suppl. Fig. 7.**
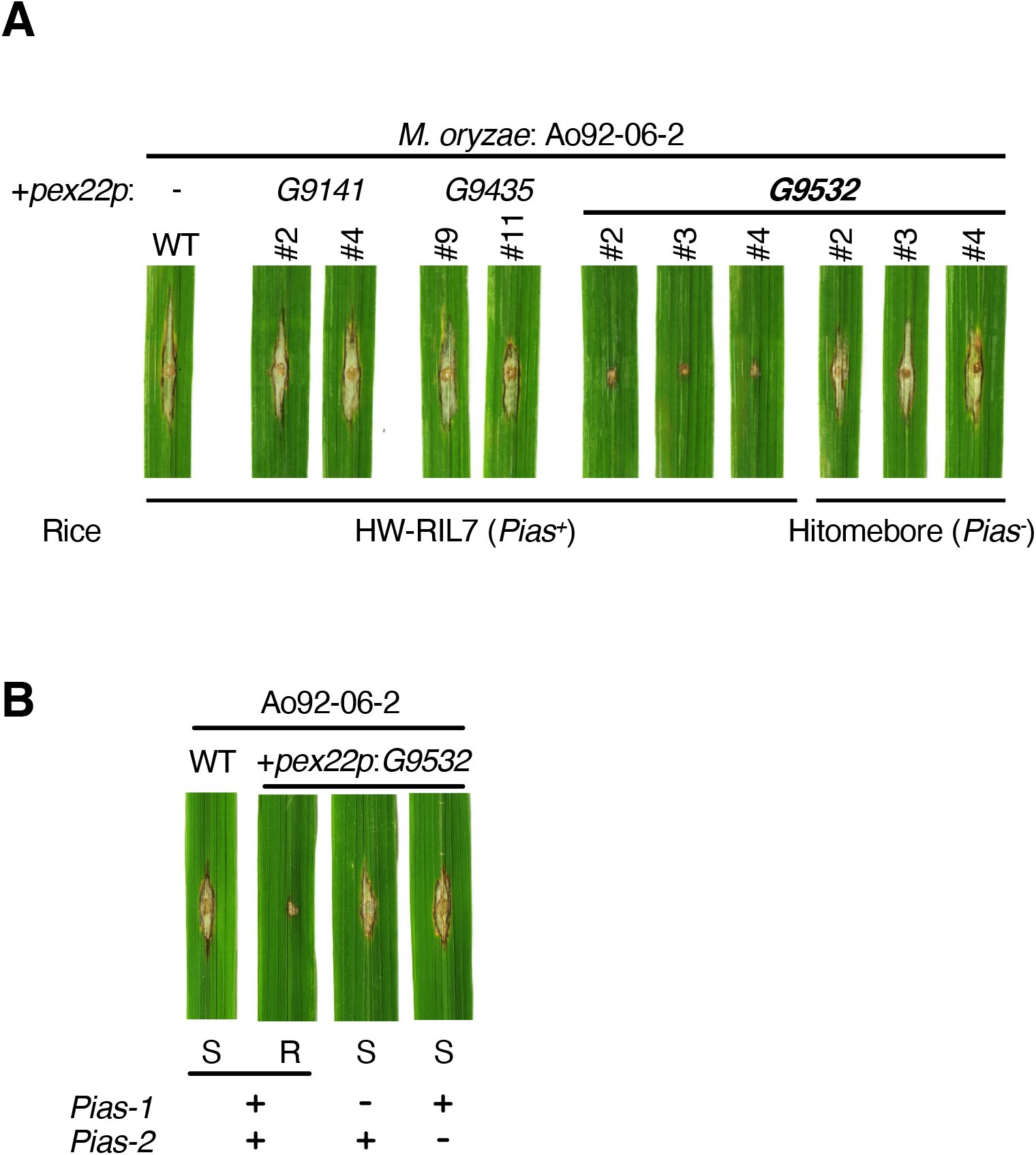
The expressed gene *G9532* of *M. oryzae* isolate 2012-1 is *AVR-Pias*. (A) Results of inoculation of rice line HW-RIL7 with *Pias* or Hitomebore without *Pias* with *M. oryzae* isolate Ao92-06-2 wild type (WT) or Ao92-06-2 with the *G9141*, *G9435*, or *G9532* transgene, all driven by the *pex22* promoter. When Ao92-06-2 contained *G9532*, the interaction became incompatible, indicating that *G9532* is *AVR-Pias*. (B) Introduction of *pex22p*:*G9532* into *M. oryzae* isolate Ao92-06-2 confers avirulence to the pathogen against HW-RIL7 with *Pias-1* and *Pias-2*. Knockout of the host gene *Pias-1* or *Pias-2* compromises the avirulence caused by *G9532*.

**Suppl. Fig. 8.**
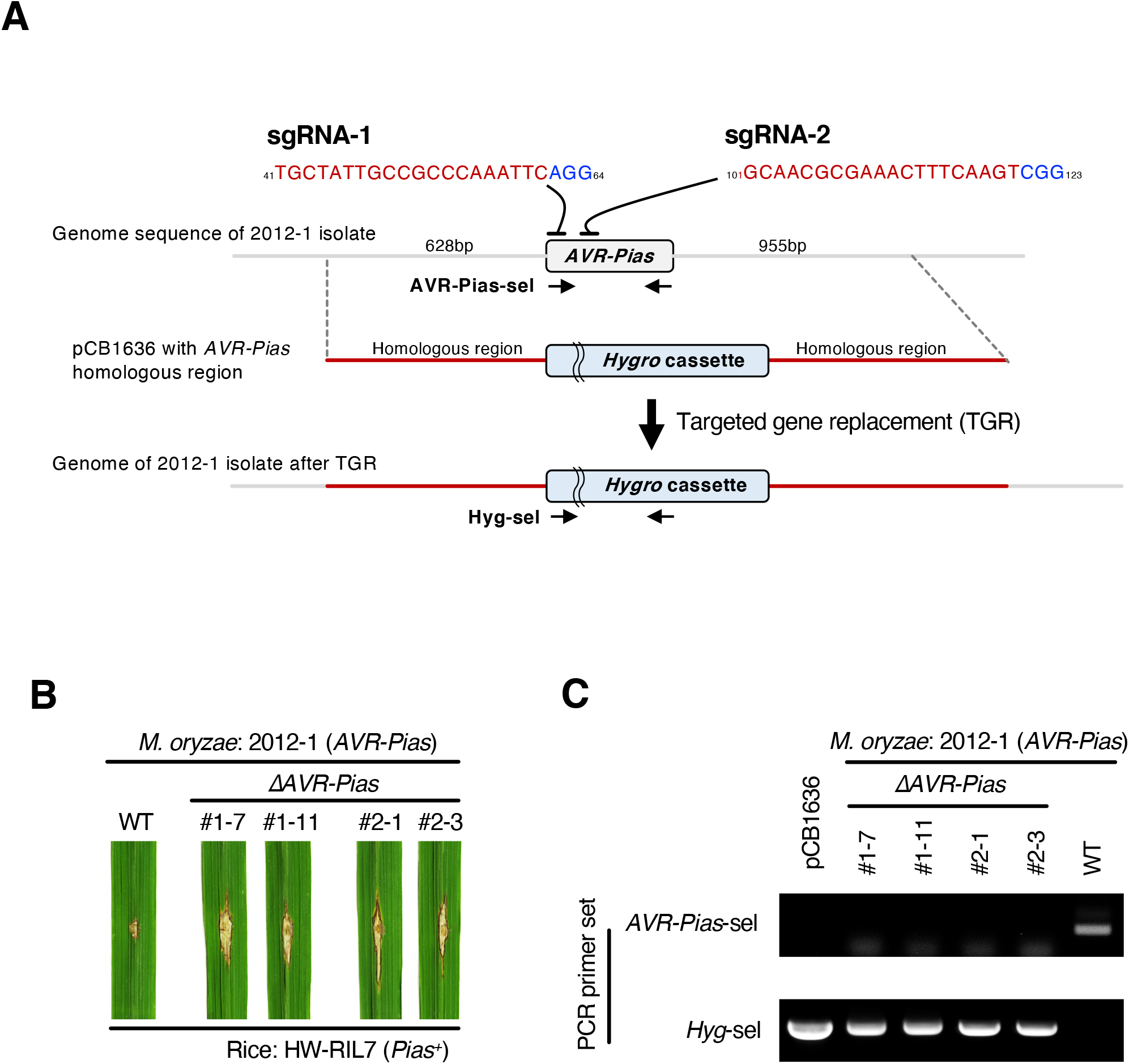
Deletion of *AVR-Pias* from *M. oryzae* isolate 2012-1 causes a loss of avirulence against rice line HW-RIL7 with *Pias*. (A) Schematic overview of the target gene replacement (TGR) strategy at the *AVR-Pias* locus using RNA-guided nuclease. The PAM is marked with blue letters, and the sgRNA sequence is marked with red letters. The primers used for PCR analysis are indicated by horizontal arrows. (B) Inoculation assay of rice line HW-RIL7 with *Pias* wild type (WT) and TGR transformants. (C) PCR analysis of TGR events at the *AVR-Pias* locus. Upper and lower images show PCR results using *AVR-Pias-*and *Hygromycin* (*Hyg*)-specific primers, respectively. pCB1636: Replacement vector containing the hygromycin resistance gene and genomic regions neighboring *AVR-Pias*. The isolates corresponding to lanes 2–5 show compatibility with rice line HW-RIL7 with *Pias*, whereas the wild-type isolate (2012-1) in lane 6 is incompatible with HW-RIL7.

**Suppl. Fig. 9.**
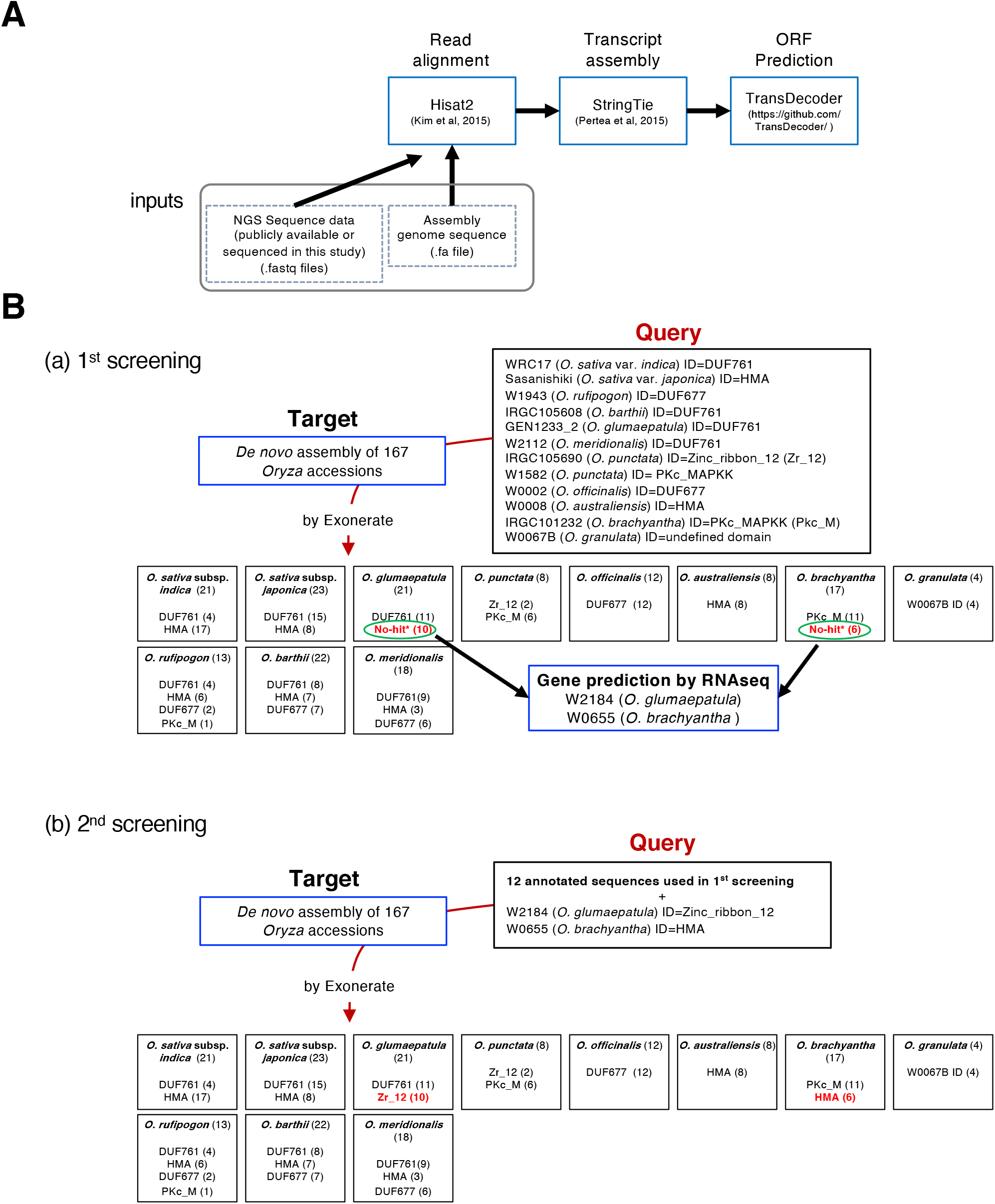
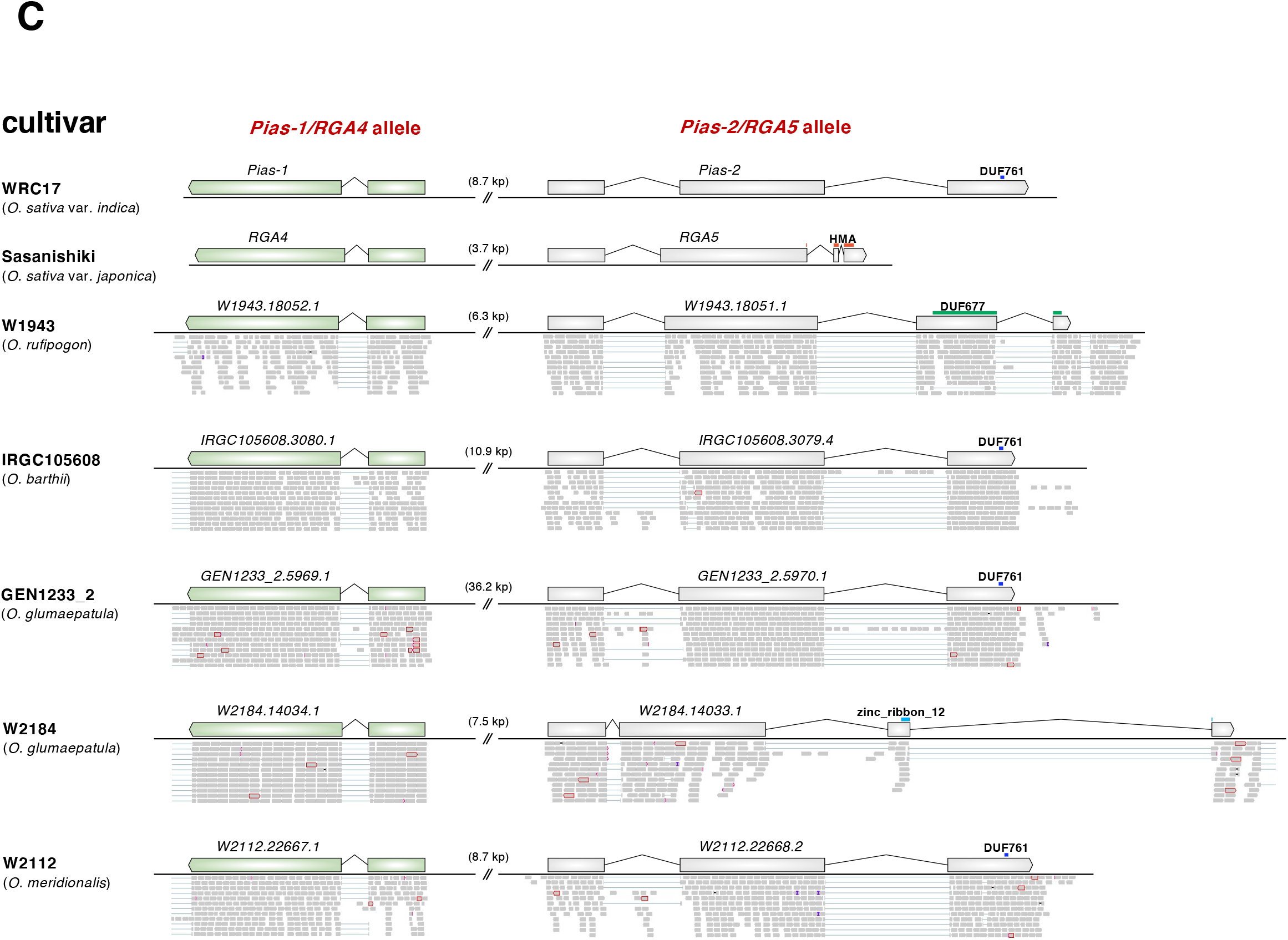
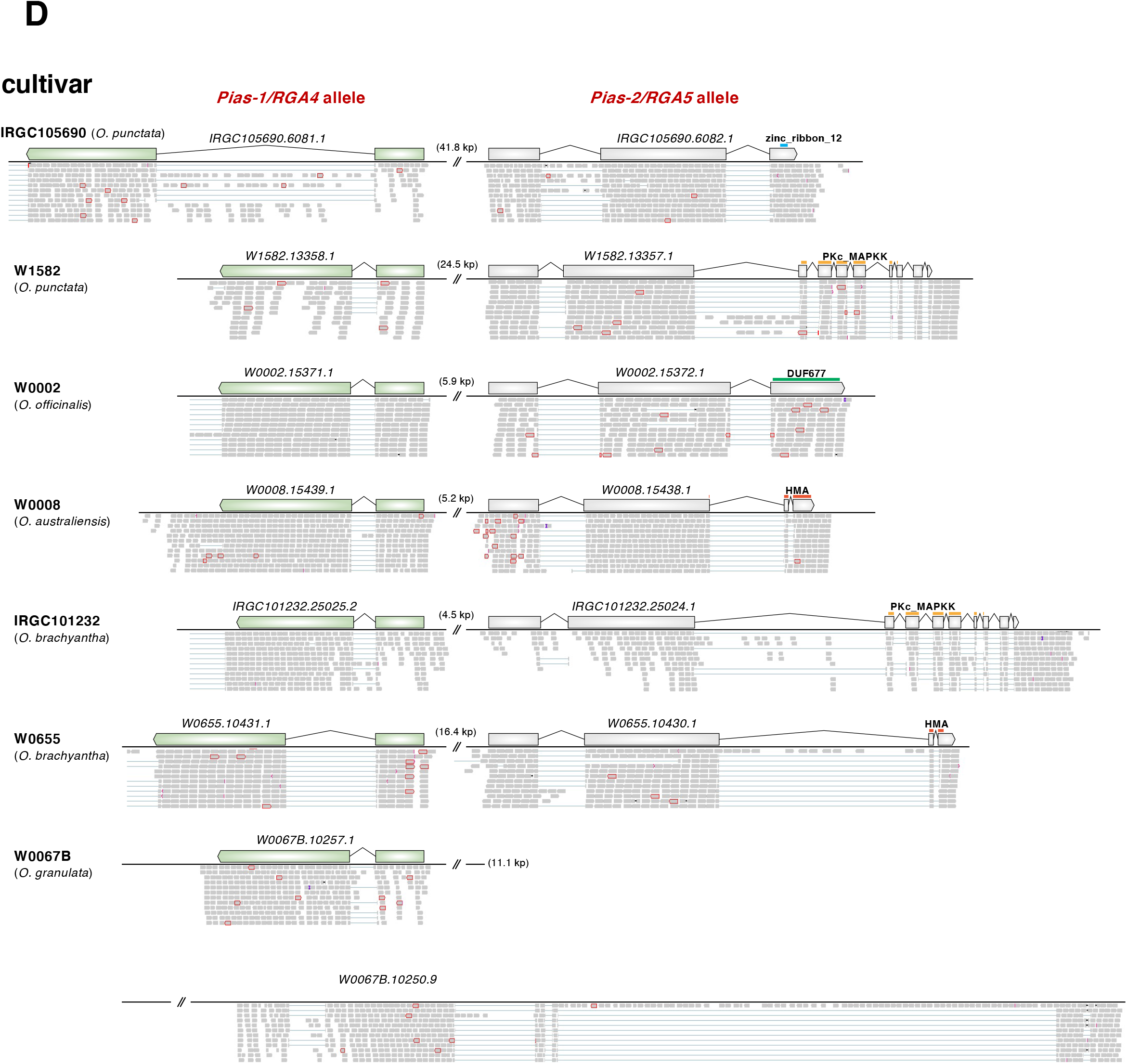
Prediction of ID sequences in the Pias/Pia sensor NLRs of the *Oryza* species. (A) Overview of gene prediction methods used in this study. (B) Gene models of *Pias-1/RGA4* and *Pias-2/ RGA5* of 12 *Oryza* samples supported by RNA-seq data were used as queries to annotate IDs in the genome assemblies of 167 *Oryza* samples using Exonerate (http://www.ebi.ac.uk/~guy/exonerate). However, 10 samples of *O. glumaepatula* and six samples of *O. brachyantha* did not match known domains. Therefore, we incorporated RNA-seq data of each sample from the two species, resulting in the annotation of the Zinc_ribbon_12 (*O. glumaepatula*) and HMA (*O. brachyantha*) IDs. In the second round, we used the 12 samples used in the first round of Exonerate as well as two new samples (*O. glumaepatula* W2184 and *O. brachyantha* W0655) as queries to infer IDs in the assembled genomes of the 167 *Oryza* samples. (C) RNA-seq read alignment across the genome sequence of the *Pias/Pia* homologous region in the AA genome *Oryza* species as visualized by IGV (Robinson et al. 2011). (D) RNA-seq read alignment across the genome sequence of the *Pias/Pia* homologous region in the non-AA genome *Oryza* species as visualized by IGV (Robinson et al. 2011).

**Suppl. Fig. 10.**
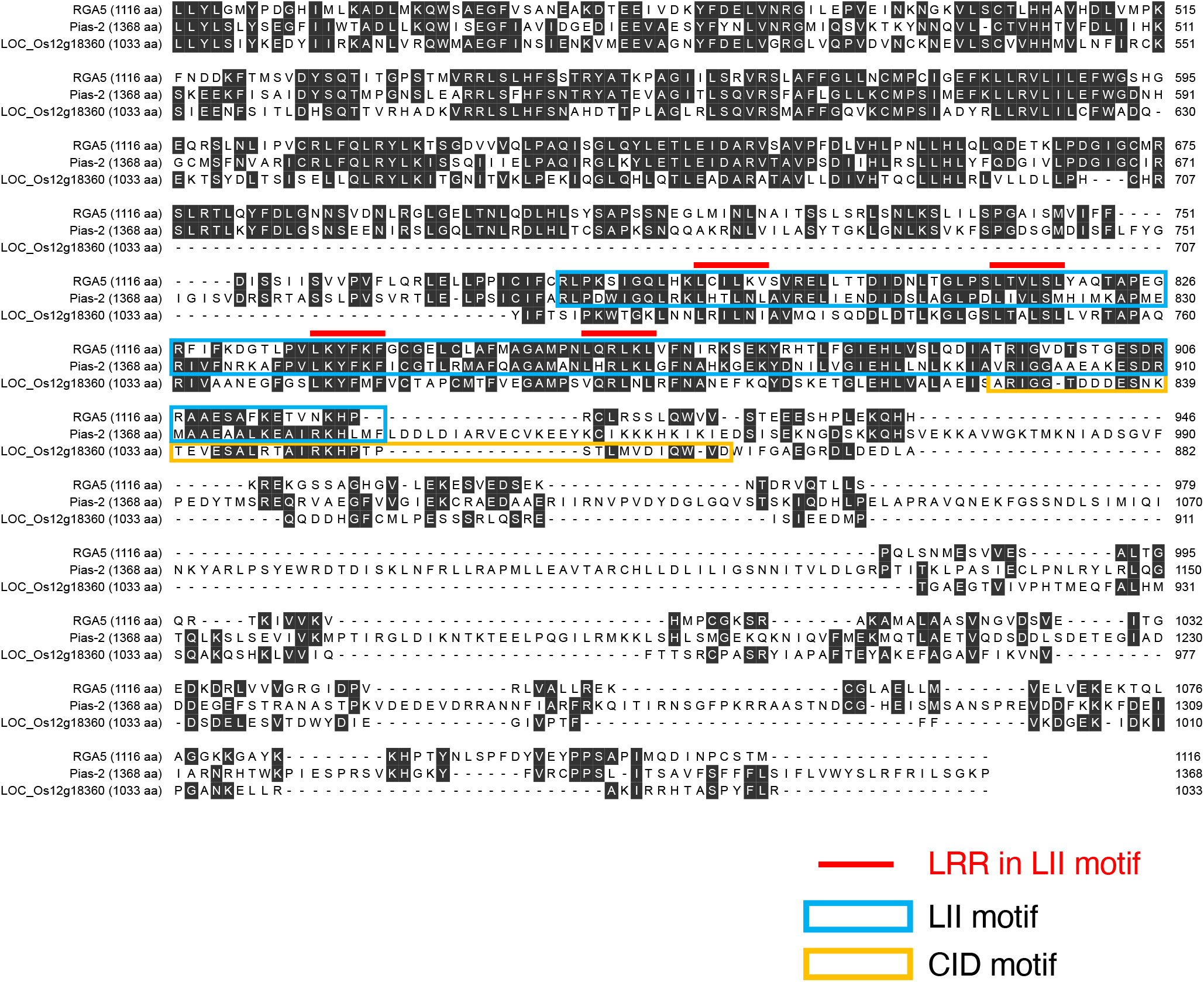
Positions of the LII motif (this study) and the CID motif (Bailey et al. 2018). Amino acid sequence alignment of RGA5, Pias-2, and LOC_Os12g18360 and the positions of the LII motif (this study) and the CID motif as described by Bailey et al. (2018). The aqua and orange boxes indicate the LII and CID motif regions, respectively. The red lines indicate LRR in the LII motif.

**Suppl. Fig. 11.**
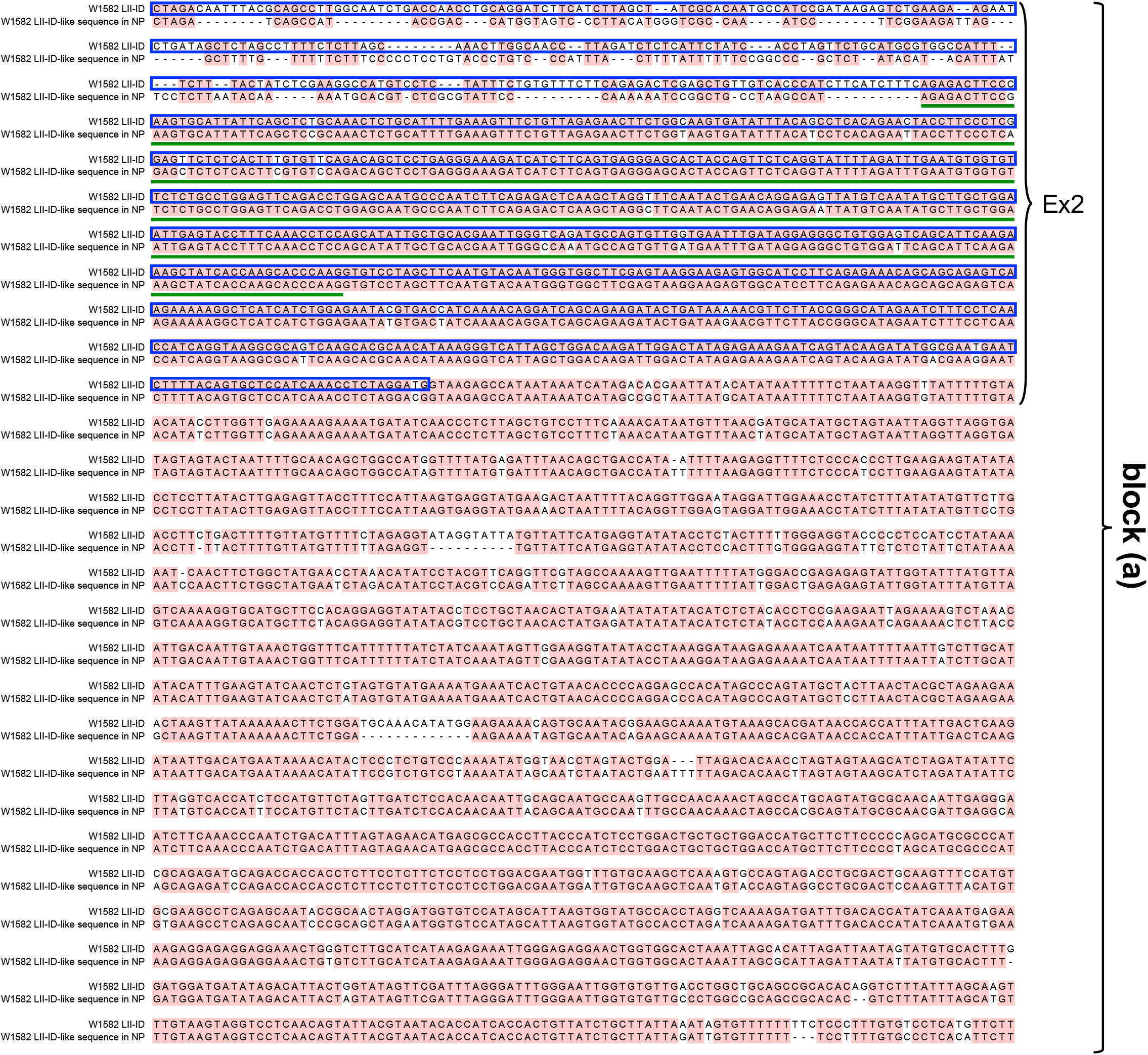
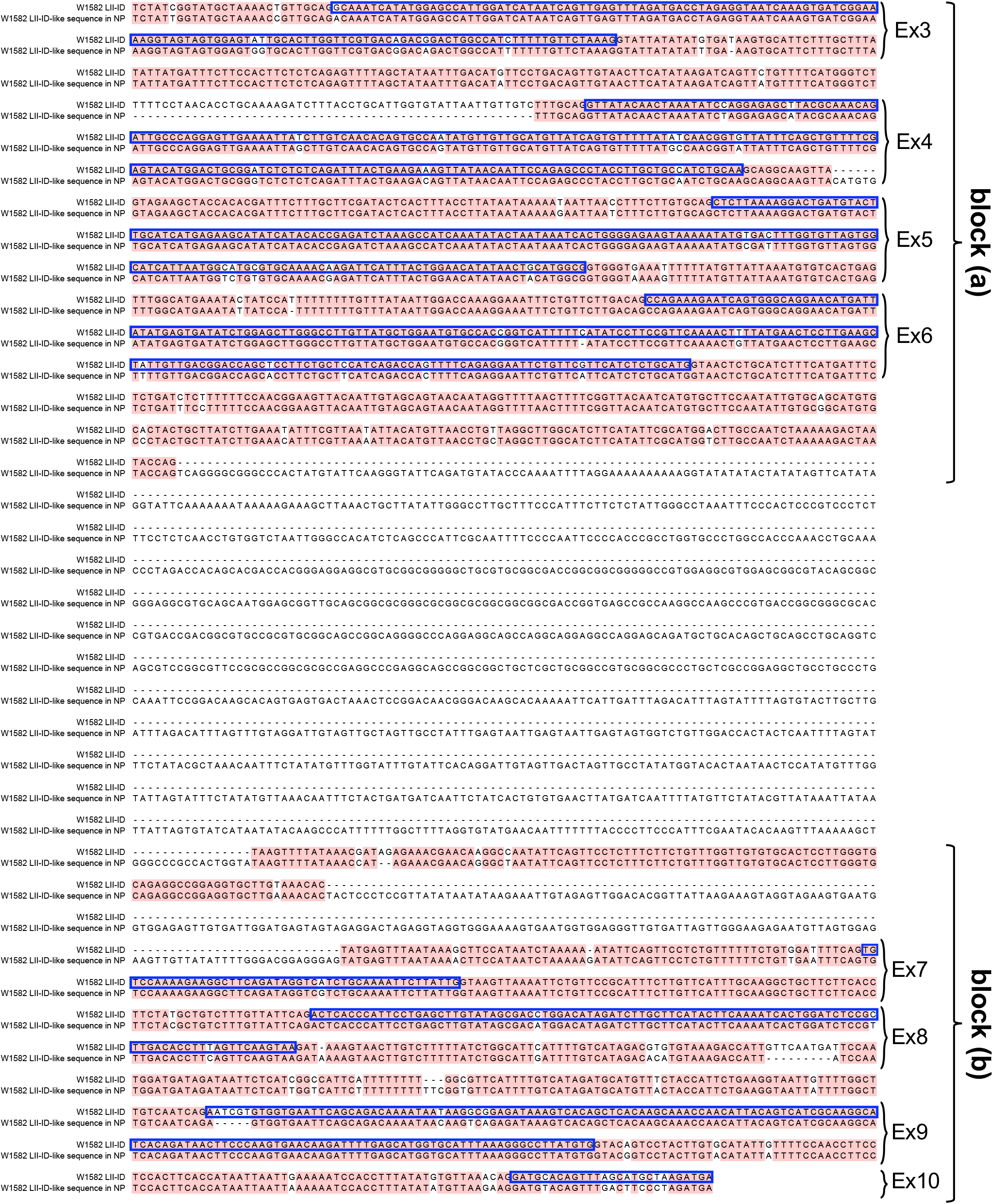
The *O. punctata* W1582 LII-ID sequence is conserved in the downstream sequence of *Pias* in *O. sativa* cv. Nipponbare. Alignment of the DNA sequences of the *O. punctata* W1582 LII-ID and the W1582 LII-ID-like sequence in *N. sativa* Nipponbare. The blue boxes and green lines indicate the exon sequence of *O. punctata* W1582 Pias-2 homolog and the LII region, respectively.

**Suppl. Fig. 12.**
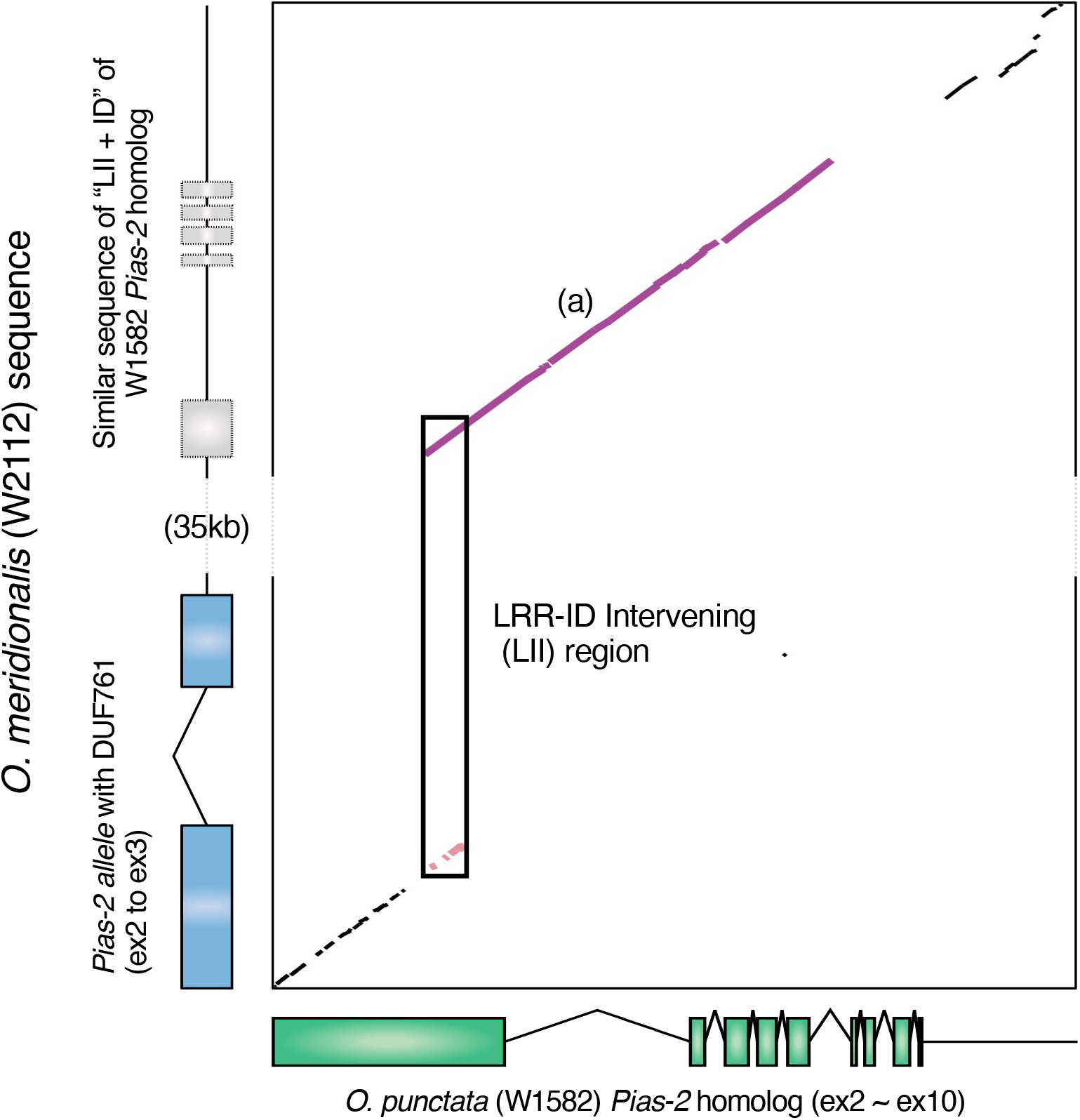
The LII + ID sequence of the *O. punctata* (W1582) *Pias-2* homolog is conserved in the downstream sequence of *O. meridionalis* (W2112) Pias-2. Dot-plot analysis of *O. punctata* (W1582) *Pias-2* homolog and the genome sequence of *O. meridionalis* (W2112). The purple line corresponds to block (a) in Figure 2. The block (b) sequence in Figure 2 is deleted in *O. meridionalis* (W2112).

**Supple. Fig. 13.**
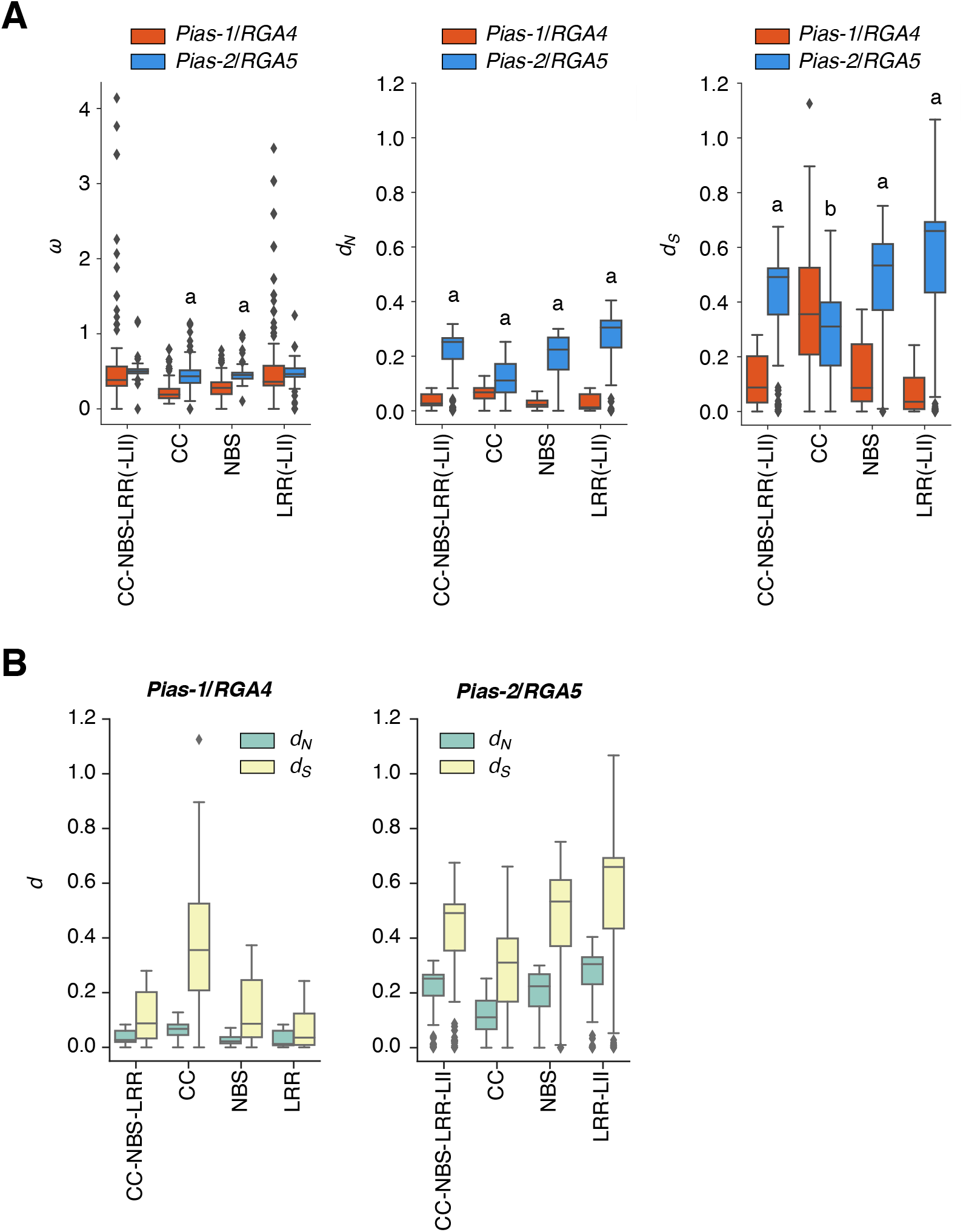
(A) Comparison of *ω*, *d_N_*, and *d_S_* between *Pias-1/RGA4* and *Pias-2/RGA5*. The ‘a’ indicates that *Pias-2/RGA5* is significantly larger than *Pias-1/RGA4* by two-sided Welch’s t-test (*p* < 0.0001). The ‘b’ indicates that *Pias-2/RGA5* is significantly smaller than *Pias-1/RGA4* by two-sided Welch’s t-test (*p* < 0.0001). **(B) Comparison between *d_N_*, and *d_S_* in *Pias-1/RGA4* and *Pias-2/RGA5*.**

**Suppl. Fig. 14.**
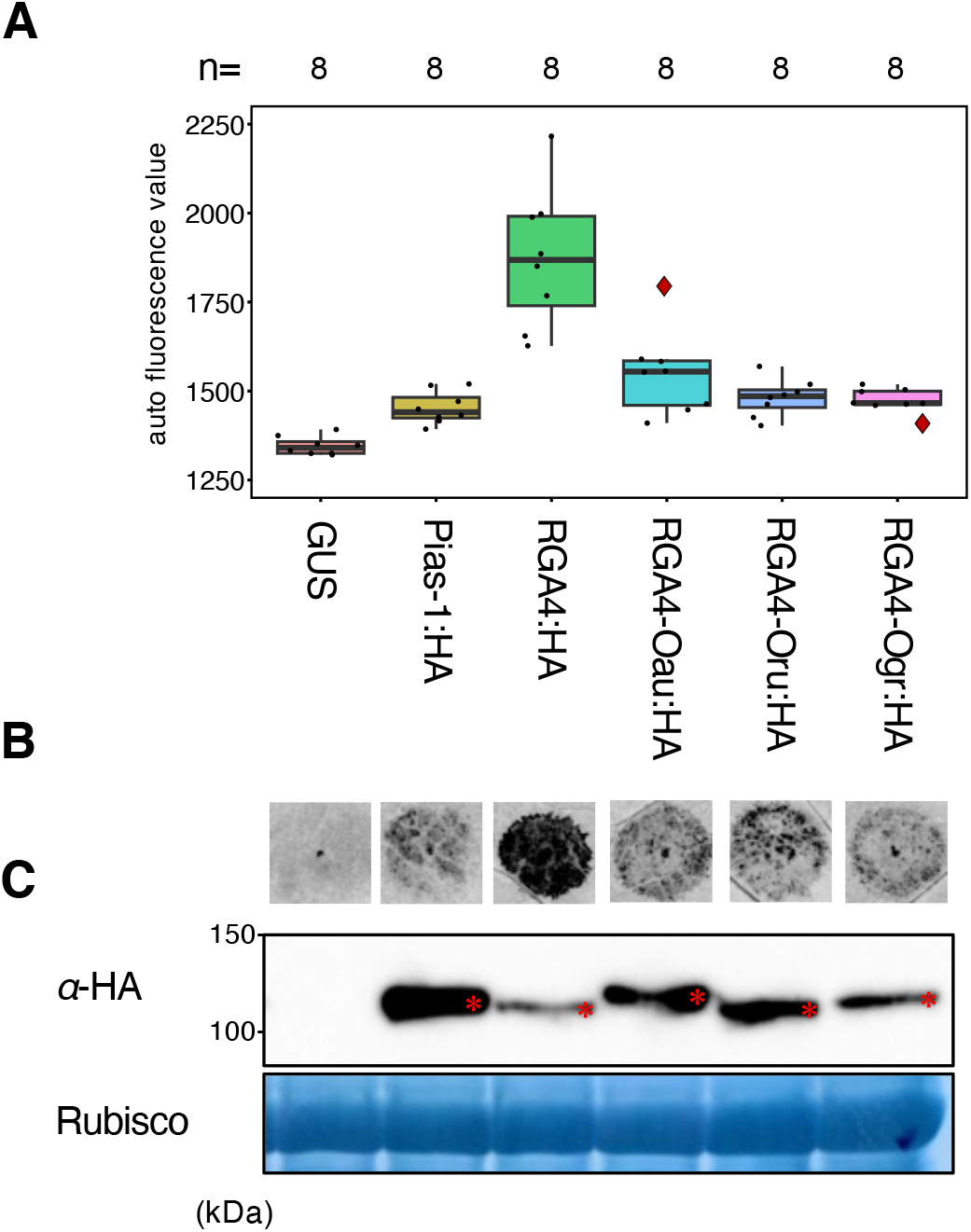
HR-like cell death after overexpression of helper NLRs of the RGA4/Pias-1 lineage in *benthamiana.* (A) Boxplots of autofluorescence values after transient expression of RGA4/Pias-1 homologs. GUS vector was included as a control. The number of spots inoculated with *A. tumefaciens* are indicated at the top of the boxplot. Box bounds represent the 25th and 75th percentile, center line represents the median, and whiskers indicate the range of the maximum or minimum data within a 1.5 x interquartile range (IQR). (B) Representative image of Pias-1/RGA4-meditated HR in *N. benthamiana*. (C) Immunoblot analysis of Pias-1:HA, RGA4:HA, RGA4-Oau:HA (cloned from *O. australiensis* accession W0008), RGA4-Oru:HA (cloned from *O. rufipogon* accession W1943), and RGA4-Ogr:HA (cloned from *O. granulata* accession W0067B) proteins detected by anti-HA (*α*-HA) antibody. Coomassie blue staining of Rubisco small subunit shows equal protein loading. The protein bands expressed from the constructs are marked by red asterisks. The positions of molecular size marker are indicated on the left (kDa)

**Suppl. Fig. 15.**
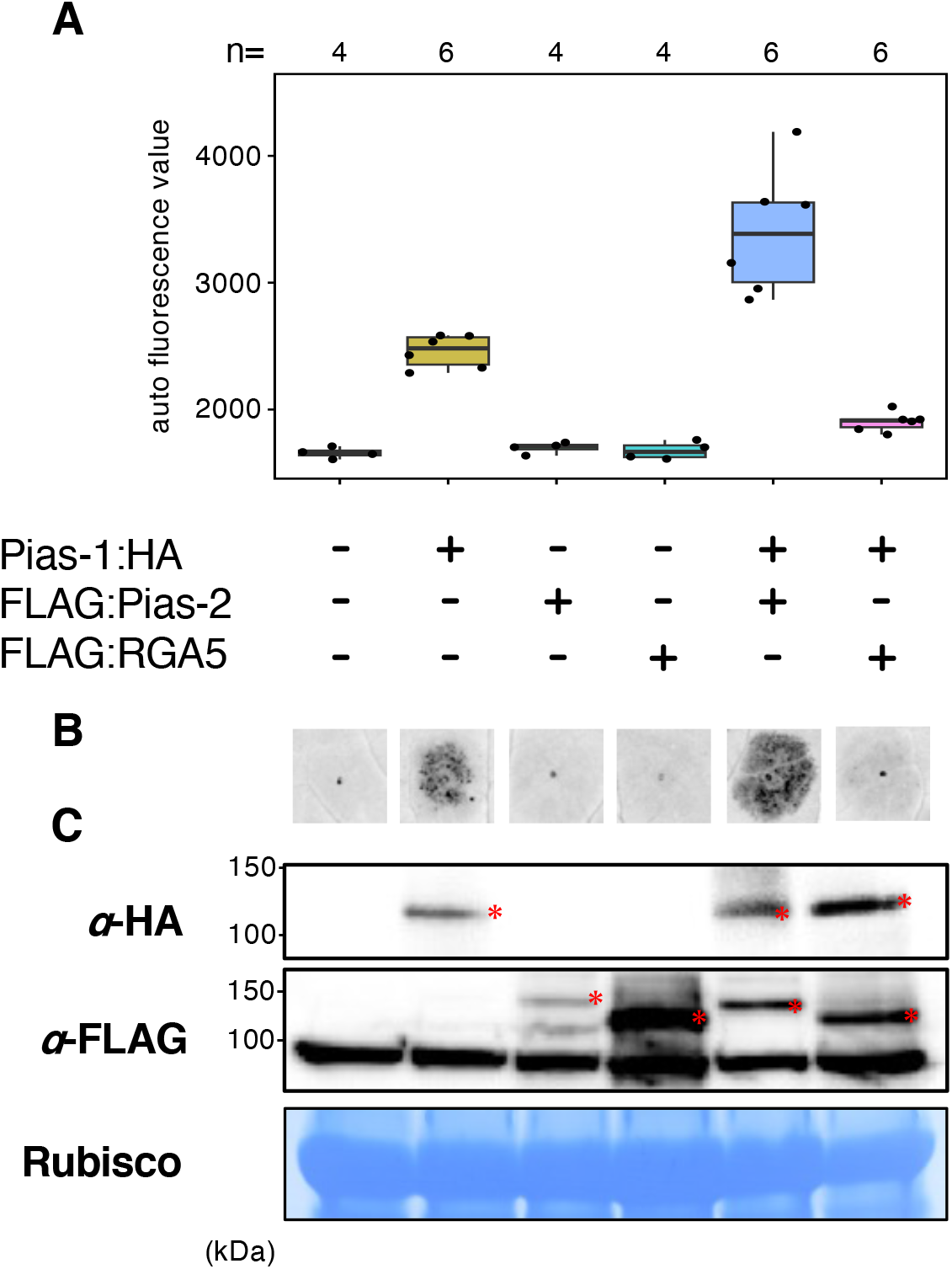
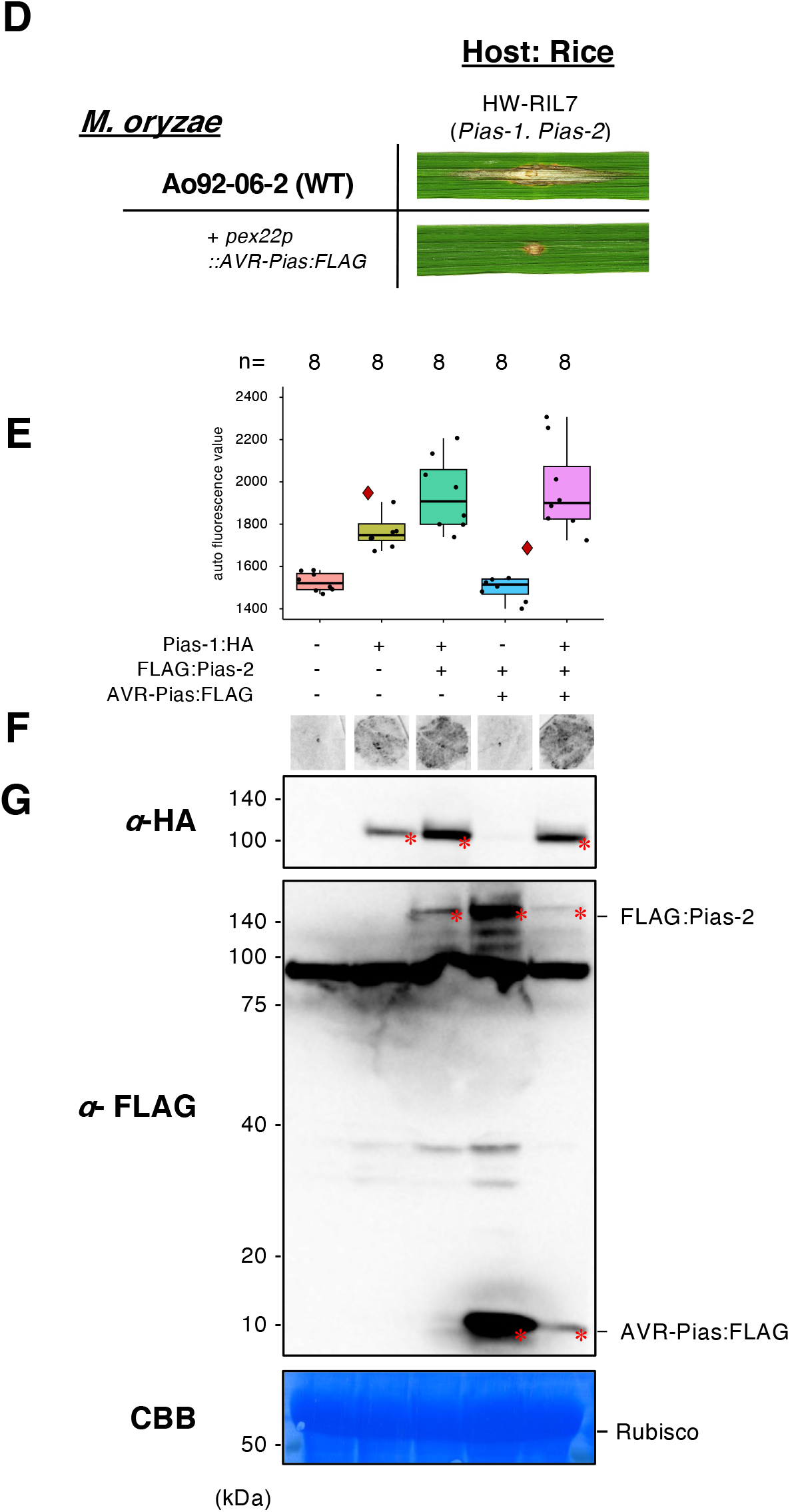
HR-like cell death after overexpression of Pias-1:HA helper and Pias-1/RGA5 sensors in *N. benthamiana*. (A) Boxplots of autofluorescence values after transient expression of Pias-1:HA, FLAG:Pias-2, and FLAG:RGA5 separately or in combination in *N. benthamiana*. The number of spots inoculated with *A. tumefaciens* are indicated at the top of the boxplot. Box bounds represent the 25th and 75th percentile, center line represents the median, and whiskers indicate the range of the maximum or minimum data within a 1.5 x interquartile range (IQR). (B) Representative image of HR after transient expression of Pias-1:HA, FLAG:Pias-2, and FLAG:RGA5 separately or in combination in *N. benthamiana*. (C) Immunoblot analysis of Pias-1:HA protein detected by anti-HA (*α*-HA) antibody and FLAG:Pias-2 and FLAG:RGA5 proteins detected by anti-FLAG antibody. Coomassie blue staining of Rubisco small subunit shows equal protein loading. The protein bands expressed from the constructs are marked by red asterisks. The positions of molecular size marker are indicated on the left (kDa). **HR-like cell death after overexpression of Pias-1 helper, Pias-2 sensor, and AVR-Pias in *N. benthamiana*.** (D) The rice line HW-RIL7 with *Pias* recognizes the *M. oryzae* Ao-92-06-2 isolate with *AVR-Pias:FLAG* (Ao92-06-2+*pex22p*:*AVR-Pias:FLAG*) and shows resistance to this isolate. (E) Boxplots of autofluorescence values after transient expression of Pias-1:HA, FLAG:Pias-2, and AVR-Pias:FLAG separately or in combination in *N. benthamiana*. The number of spots inoculated with *A. tumefaciens* are indicated above the boxplot. Box bounds represent the 25th and 75th percentile, center line represents the median, and whiskers indicate the range of the maximum or minimum data within a 1.5 x interquartile range (IQR). (F) Representative image of HR after transient expression of Pias-1:HA, FLAG:Pias-2, and FLAG:RGA5 separately or in combination in *N. benthamiana*. (G) Immunoblot analysis of Pias-1:HA protein detected by anti-HA (*α*-HA) antibody and FLAG:Pias-2 and AVR-Pias:FLAG proteins detected by anti-FLAG antibody. Coomassie blue staining of Rubisco small subunit shows equal protein loading. The protein bands expressed from the constructs are marked by red asterisks. The positions of the molecular size marker are indicated on the left (kDa).

**Suppl. Fig. 16.**
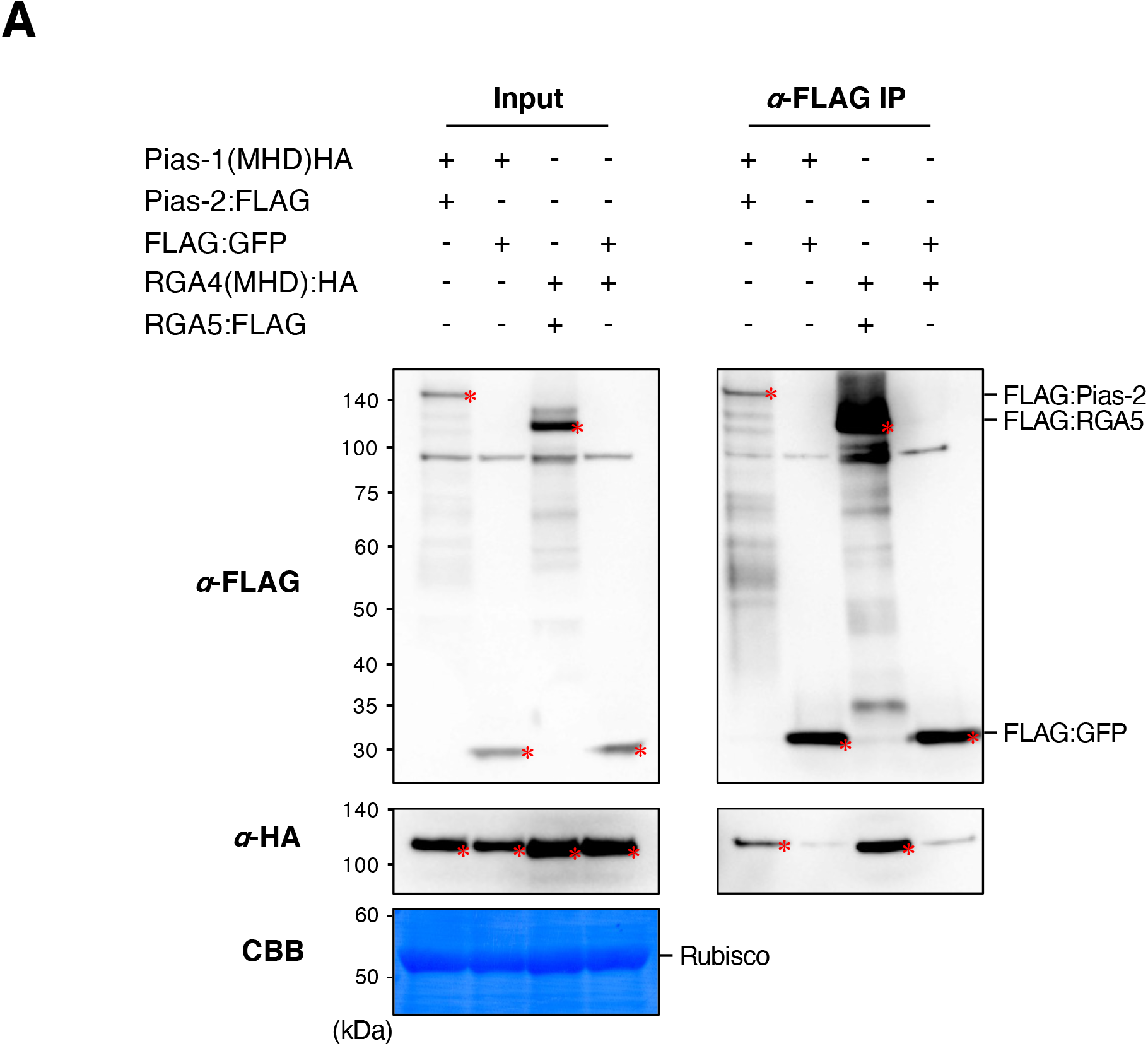
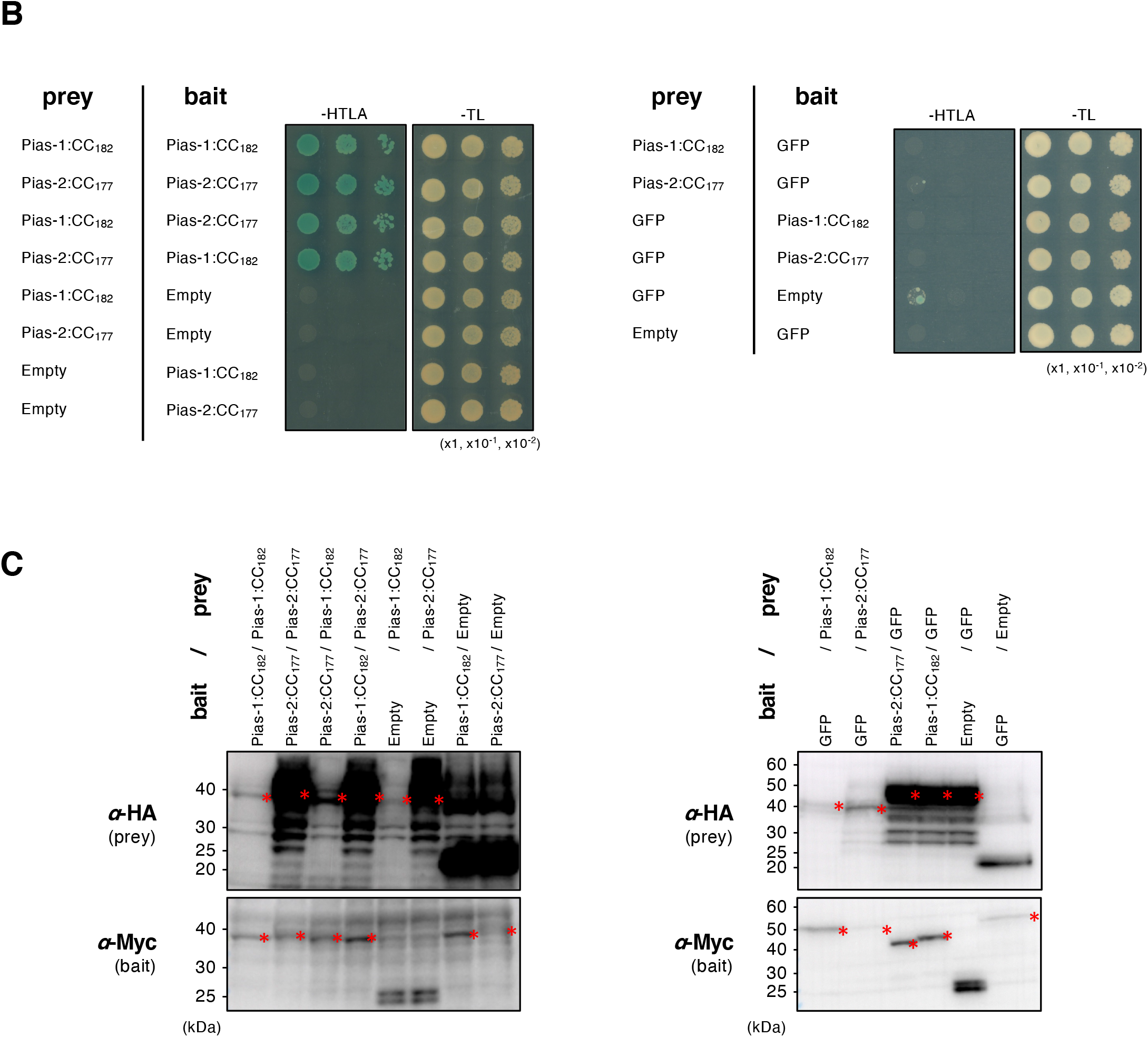
Co-immunoprecipitation shows that Pias-1 interacts with Pias-2. Co-immunoprecipitation (Co-IP) of Pias-1 (MHD mutant):HA with FLAG:Pias-2 or FLAG:GFP (negative control), as well as Co-IP of RGA4 (MHD mutant):HA with FLAG:RGA5 or FLAG:GFP (negative control) were performed. Pias-1 (MHD mutant):HA and FLAG:Pias-2 or FLAG:GFP were transiently co-expressed in *N. benthamiana.* Similarly, RGA4 (MHD mutant):HA and FLAG:RGA5 or FLAG:GFP were transiently co-expressed in *N. benthamiana*. Instead of the wild-type Pias-1 and RGA4, their MHD mutants (TYG to MHD in ARC2 subdomain as described in Cesari et al. 2014) were used to avoid HR-like cell death that reduces protein accumulation. We judge the weak bands detected by anti-HA antibody after Co-IP in Pias-1(MHD):HA+FLAG:GFP (lane 2) and RGA4(MHD):HA+FLAG:GFP (lane 4) are non-specific. Bound fractions were analyzed by immunoblotting using anti-FLAG and anti-HA antibodies. Coomassie blue staining of the Rubisco small subunit shows equal protein loading. The protein bands expressed from the constructs are marked by red asterisks. The positions of the molecular size marker are indicated on the left (kDa). We obtained similar results in three independent experiments. **Yeast two-hybrid assays indicate that the Pias-1 CC domain and Pias-2 CC domain homo- and heterodimerize.** (A) The Pias-1 and Pias-2 CC domains form homo- and heterocomplexes. A dilution series of yeast cells expressing a GAL4-AD and GAL4-BD fusion of Pias-1 1-182 (Pias-1:CC_1-182_) and/or Pias-2 1-177 (Pias-2:CC_1-177_) on selective media lacking Trp, Leu, Ade, and His (–HTLA) and non-selective media lacking Trp and Leu (–TL) with 5-bromo-4-chloro-3-indolyl α-D-galactopyranoside (X-a-gal). GFP was used as the negative control. Photographs were taken 5 days after growth. (C). Immunoblot analysis confirms the protein production in the Y2H assay shown in (B). The bait protein was tagged with the Myc epitope and the prey protein with the HA epitope. The protein bands expressed from each vector are marked by red asterisks. The positions of the molecular size marker are indicated on the right (kDa). We obtained similar results in three independent experiments. We attempted Y2H assay using the full-length Pias-1 and Pias-2 constructs. However, we could not detect interactions between the full-length Pias-1 and Pias-2.

**Suppl. Fig. 17.**
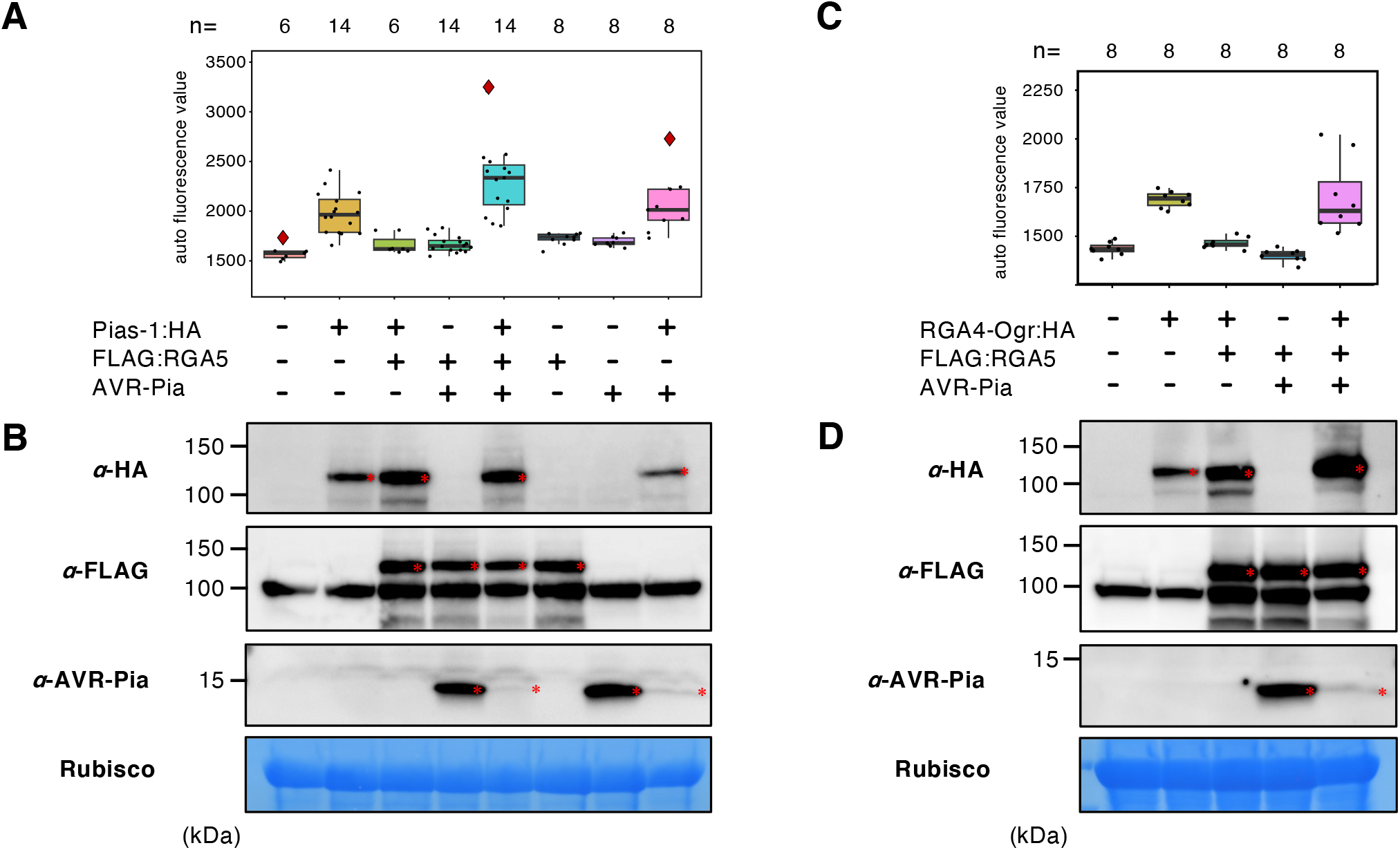
HR-like cell death caused by Pias1 and RGA4-Ogr expression is suppressed by RGA5 expression, and additional AVR-Pia expression induces cell death. (A) Boxplots of autofluorescence values after transient expression of Pias-1:HA, FLAG:RGA5, and AVR-Pia separately or in combination in *N. benthamiana* leaves. The number of spots inoculated with *A. tumefaciens* are indicated at the top of the boxplot. Box bounds represent the 25th and 75th percentile, center line represents the median, and whiskers indicate the range of the maximum or minimum data within a 1.5 x interquartile range (IQR). (B) Immunoblot analysis of Pias-1:HA protein detected by anti-HA (*α*-HA) antibody, FLAG:RGA5 protein detected by anti-FLAG antibody, and AVR-Pia detected by anti-AVR-Pia antibody. Coomassie blue staining of Rubisco small subunit shows equal protein loading. The protein bands expressed from the constructs are marked by red asterisks. The positions of molecular size markers are indicated on the left (kDa). (C) Boxplots of autofluorescence values after transient expression of RGA4-Ogr:HA, FLAG:RGA5, and AVR-Pia separately or in combination in *N. benthamiana* leaves.The number of spots inoculated with *A. tumefaciens* are indicated at the top of the boxplot. Box bounds represent the 25th and 75th percentile, center line represents the median, and whiskers indicate the range of the maximum or minimum data within a 1.5 x interquartile range (IQR). (D) Immunoblot analysis of RGA4-Ogr:HA protein detected by anti-HA (*α*-HA) antibody, FLAG:RGA5 protein detected by anti-FLAG (*α*-FLAG) antibody, and AVR-Pia detected by anti-AVR-Pia (*α*-AVR-Pia) antibody. Coomassie blue staining of Rubisco small subunit shows equal protein loading. The protein bands expressed from the constructs are marked by red asterisks. The positions of molecular size markers are indicated on the left (kDa).

**Suppl. Fig. 18.**
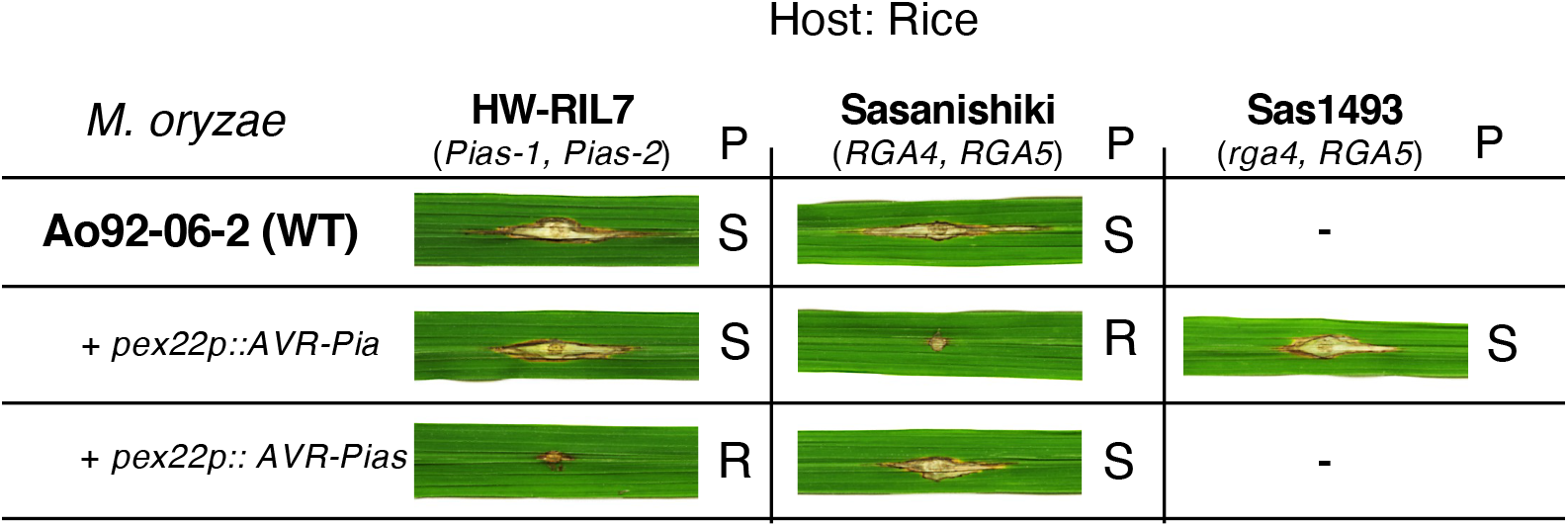
Results of inoculation of HW-RIL7, Sasanishiki, and Sas1493 rice plants with *M. oryzae* isolates Ao92-06-2, Ao92-06-2 +*pex22p:AVR-Pia*, and Ao92-06-2 +*pex22p:AVR-Pias*.

**Suppl. Fig. 19.**
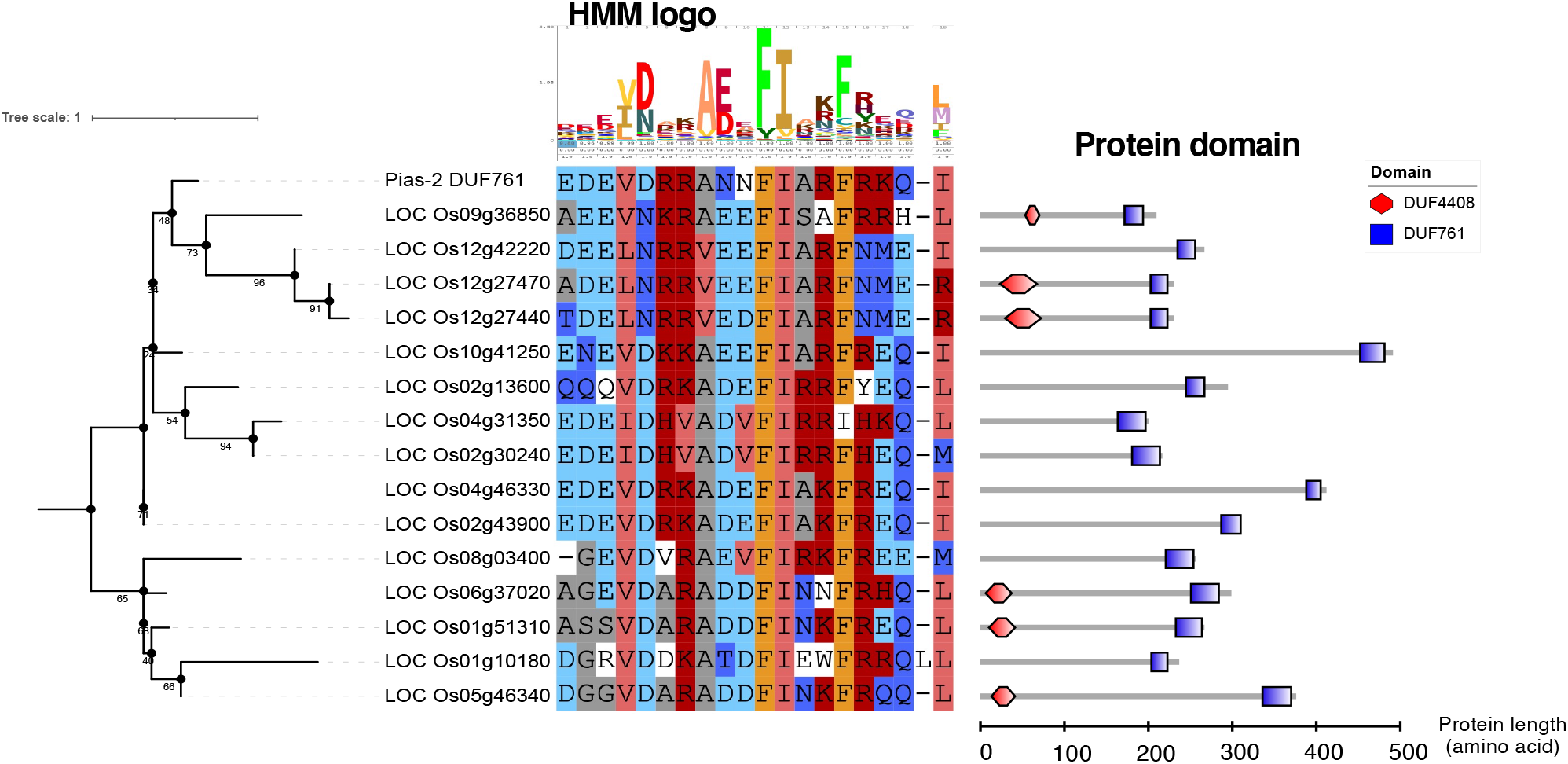
The DUF761-containing gene family in rice. A BLASTP search was performed using the amino acid sequence of the DUF761 domain of Pias-2 as a query. Fifteen DUF761 domain-containing genes were retrieved using a cutoff e-value < 10. The middle panel shows the amino acid sequence alignment, and the right panel shows the domain structure of each protein.

**Suppl. Table 1.** Summary of Illumina sequencing results used for RaIDeN methods.

| Line | Sequencing platform | Trim adapters and quality filter | Total no. of paired-end (PE) reads after filtering | Max length of short reads | Purpose |
| --- | --- | --- | --- | --- | --- |
| **<u>Whole genome sequencing</u>** |  |  |  |  |  |
| **R-parental line** |  |  |  |  |  |
| WRC17 | Miseq | No | 36,047,742 | 250 | for *de novo* assemble |
| **S-parental line** |  |  |  |  |  |
| Hitomebore | Hiseq4000 | Yes | 32,016,561 | 126 | for RaIDeN |
| **S-RILs** |  |  |  |  | for RaIDeN |
| #22 | Hiseq4000 | Yes | 30,830,739 | 126 | for RaIDeN |
| #25 | Hiseq4000 | Yes | 39,330,775 | 126 | for RaIDeN |
| #34 | Hiseq4000 | Yes | 36,109,189 | 126 | for RaIDeN |
| #37 | Hiseq4000 | Yes | 34,054,341 | 126 | for RaIDeN |
| #39 | Hiseq4000 | Yes | 42,282,097 | 126 | for RaIDeN |
| #53 | Hiseq4000 | Yes | 34,013,397 | 126 | for RaIDeN |
| **<u>RNAseq</u>** |  |  |  |  |  |
| *M. oryzae* infected leaf of WRC17 | Nextseq500 | Yes | 39,744,107 | 75 | for RaIDeN |

**Suppl. Table 2.** Summary metrics of genome assembly of the resistance rice line WRC17. Genome assembly was performed by *DISCOVAR* software.

|  |  |
| --- | --- |
| No. of contigs (>1,000bp) | 25,278 |
| Contig N50 (kb) | 55.1 |
| Largest contig (kb) | 459.5 |
| Total size of the assembled genome (Mb) | 356.2 |

**Suppl. Table 3.** Summary of 38 candidate resistance gene analogs (RGAs) expressed in WRC17 leaves and possibly responsible for WRC17 resistance against 2012-1 as predicted by RGAugury (Li et al. 2016). These were selected from the 853 candidate genes as identified by the RaIDeN method. From the 38 RGAs, we selected 18 genes belonging to the categories of CNL (Coiled-Coil, NBS and Leucine rich repeat protein genes: 11 genes) and NL (NBS and Leucine rich repeat protein genes: 7 genes) as the candidate NLRs conferring resistance to WRC17 against *M. oryzae* 2012-1.

| NBS encoding |  |  |  |  |  | TX | Other | RLP | RLK | TM-CC |
| --- | --- | --- | --- | --- | --- | --- | --- | --- | --- | --- |
| NBS | CNL | TNL | CN | TN | NL |  |  |  |  |  |
| 3 | 11 | 0 | 1 | 0 | 7 | 0 | 0 | 1 | 12 | 3 |

CC, C: coiled-coil T: Toll/Interleukin-1 receptor NBS, N: nucleotide-binding site L: leucine rich repeat
TX: TIR-unknown domain RLK: receptor like kinase RLP: receptor like protein TM: transmembrane

**Suppl. Table 4.** Summary of whole genome sequencing of the nine M. oryzae isolates and RNA sequencing of M. oryzae isolate 2012 for identification of AVR-Pias.

| Line | Race code | Sequencing platform | Trim adapters and quality filter | Total no. of paired-end (PE) reads after filtering | Max length of short reads | Purpose |
| --- | --- | --- | --- | --- | --- | --- |
| **<u>Whole genome sequencing</u>** |  |  |  |  |  |  |
| 2012-1 |  | Miseq | No | 3,332,000 | 250 | for *de novo* assemble |
| **S-pathogen for HW-RIL7** |  |  |  |  |  |  |
| Ina87T-56A | **r1** | Miseq | Yes | 1,776,902 | 125 | for RaIDeN |
| Ao92-06-2 | **r2** | Miseq | Yes | 2,691,934 | 125 | for RaIDeN |
| 85-141 | **r3** | Miseq | Yes | 3,034,450 | 125 | for RaIDeN |
| SL91-48D | **r4** | Miseq | Yes | 3,108,538 | 125 | for RaIDeN |
| 2403-1 | **r5** | Miseq | Yes | 3,388,670 | 125 | for RaIDeN |
| H98-315-1 | **r6** | Miseq | Yes | 2,933,673 | 125 | for RaIDeN |
| 0423-1 | **r7** | Miseq | Yes | 2,996,542 | 125 | for RaIDeN |
| Ina85-182 | **r8** | Miseq | Yes | 3,148,657 | 125 | for RaIDeN |
| **<u>RNAseq</u>** |  |  |  |  |  |  |
| *M. oryzae* isolate 2012-1 inoculated to the leaf of *Hordeum vulgare* var. Nigrate |  |  |  | **Total no. of reads (SE) after filtering** |  |  |
| 24H after inoculation |  | Nextseq500 | Yes | 15,658,123 | 75 | for RaIDeN and expression analysis |
| 48H after inoculation |  | Nextseq500 | Yes | 15,770,613 | 75 | for RaIDeN and expression analysis |

**Suppl. Table 5.**
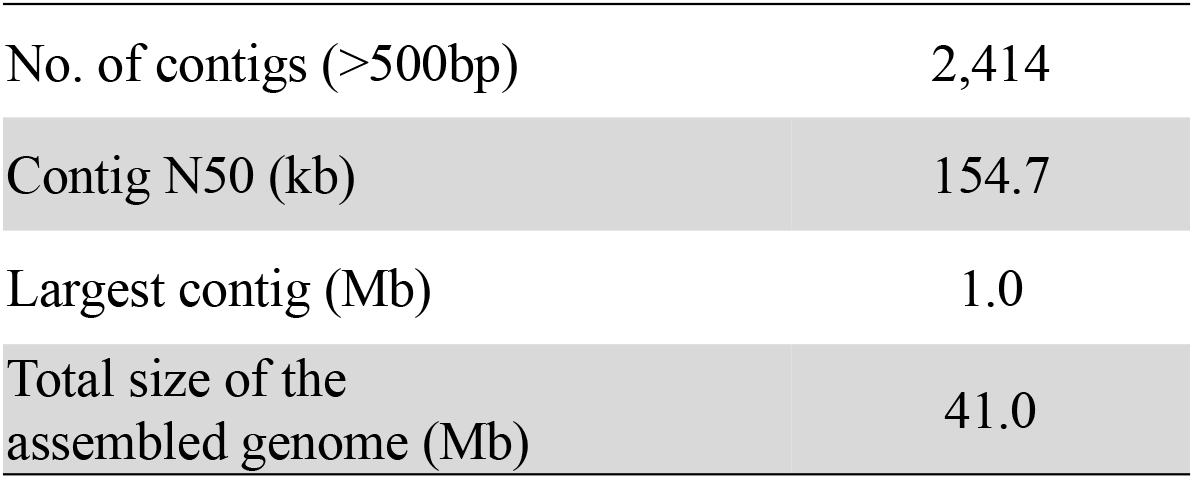
Summary metrics of genome assembly of M. oryzae 2012-1 isolate. Genome assembly was performed by *DISCOVAR* software.

**Suppl. Table 9.**
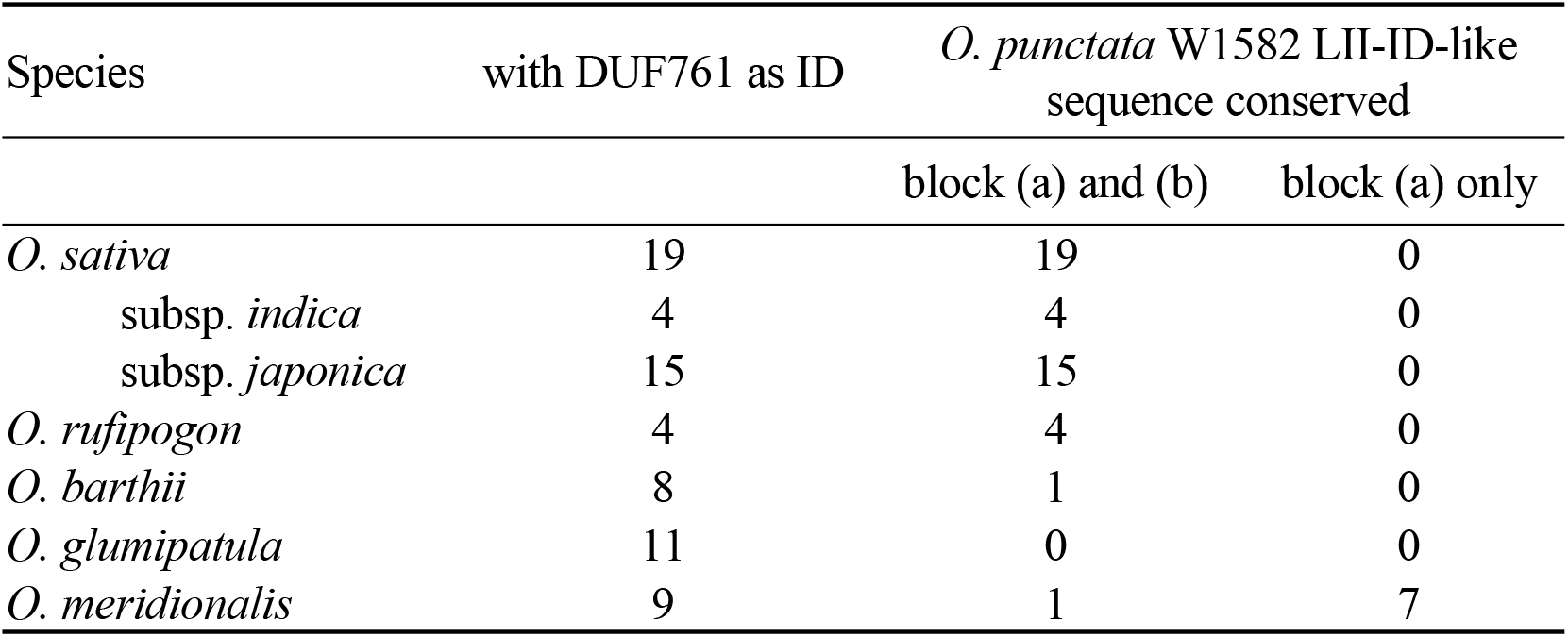
A survey of 51 A-genome *Oryza* accessions with DUF761 ID revealed that the*O. punctata* LII-ID-like downstream sequence is widely conserved. BLASTN search was performed on each genome sequence using *O. punctata* W1582 LII-ID-like sequence of Nipponbare cv. as a query. Block (a) and block (b) region correspond to those in Figure 2.DE.

| Species | with DUF761 as ID | *O. punctata* W1582 LII-ID-like sequence conserved |  |
| --- | --- | --- | --- |
|  |  | block (a) and (b) | block (a) only |
| *O. sativa* | 19 | 19 | 0 |
| subsp. *indica* | 4 | 4 | 0 |
| subsp. *japonica* | 15 | 15 | 0 |
| *O. rufipogon* | 4 | 4 | 0 |
| *O. barthii* | 8 | 1 | 0 |
| *O. glumipatula* | 11 | 0 | 0 |
| *O. meridionalis* | 9 | 1 | 7 |

**Suppl. Table 11.**
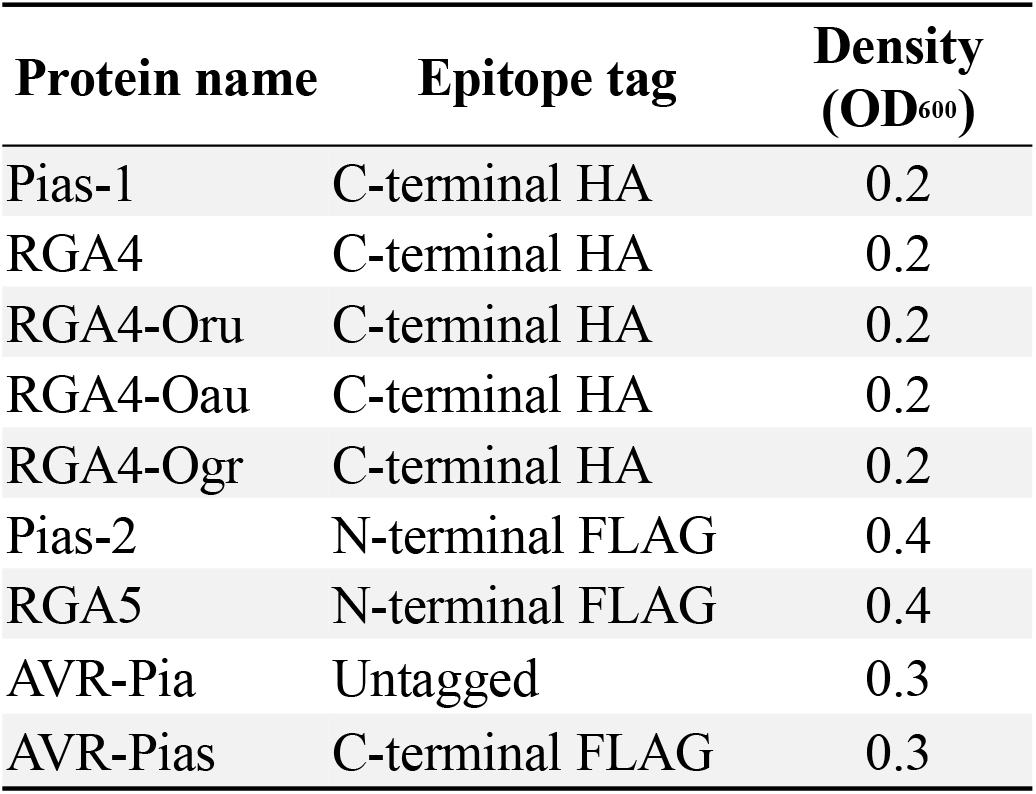
List of NLRs and AVR used in the cell death assay. OD_600_ of *Agrobacterium tumefaciens* used for agroinfiltration to *N. benthamiana* leaves is shown.

| Protein name | Epitope tag | Density<br>(OD<sub>600</sub>) |
| --- | --- | --- |
| Pias-1 | C-terminal HA | 0.2 |
| RGA4 | C-terminal HA | 0.2 |
| RGA4-Oru | C-terminal HA | 0.2 |
| RGA4-Oau | C-terminal HA | 0.2 |
| RGA4-Ogr | C-terminal HA | 0.2 |
| Pias-2 | N-terminal FLAG | 0.4 |
| RGA5 | N-terminal FLAG | 0.4 |
| AVR-Pia | Untagged | 0.3 |
| AVR-Pias | C-terminal FLAG | 0.3 |

